# Sensory neurons sprouting is dependent on osteoclast-derived extracellular vesicles involving the activation of epidermal growth factor receptors

**DOI:** 10.1101/259218

**Authors:** Estrela Neto, Luís Leitão, José Mateus, Daniela M. Sousa, Francisco Conceição, Cecília J. Alves, Richard O. C. Oreffo, Jonathan West, Paulo Aguiar, Meriem Lamghari

## Abstract

The patterning of peripheral innervation is accomplished through the tissue expression, in specific space and timeframe, of attractive or repulsive axonal guidance cues. At the bone microenvironment, neurotrophic factors such as nerve growth factor, brain-derived neurotrophic factor, vascular endothelial growth factor, netrin-1 and others were described to regulate the nerve ingrowth towards the bone compartment, by acting directly on receptors expressed at the nerve terminals. Interestingly, besides the gradient of soluble factors, neurons were described to be responsive to extracellular vesicles (EV) derived from myelinating cells and mesenchymal stem cells.

Here we provide evidence on a new mechanism by which peripheral innervation can be coordinated. We show that sensory nerves outgrowth and electric signal propagation are dependent on the EV secreted by osteoclasts, the bone resorbing cells. Furthermore, we demonstrate that the axonal sprouting is achieved through the activation of epidermal-growth factor receptor (EGFR) family signaling pathway. We proved that the EV-depleted osteoclast secretome leads to a significant decrease of neurons firing rate and axonal sprouting, concomitant with a decrease of EGFR/ErbB2 activation levels. Excitingly, the proteomic analysis of the osteoclast-derived EV cargo shows a high correlation with synaptic components reinforcing the role on sensory neurons/osteoclast crosstalk.

Our findings that osteoclast-derived EV hold effect in axonal outgrowth, contributing actively to the dynamics of the sensory neurons sprouting and electrophysiology, is a step toward unraveling target mechanisms to control electrical signal propagation and nerve fibers sprouting and consequently open new avenues for the development of innovative therapies to control bone pain.

**Significance Statement:** Sensory nerve fibers sprouting in bone pathologies is highly associated with pain. Thus, understanding the mechanisms behind sensory nerves ingrowth, sprouting and electrical activity, within the bone compartment, is essential for improving the strategies to overcome pain in bone disorders. We provide a new mechanism on the sensory nerves sprouting, indicating that the effect is dependent on the extracellular vesicles (EV) released by osteoclasts, through the epidermal growth factor receptor family targeting, by integrin independent pathways. We show different electrophysiology patterns being triggered in the presence of osteoclasts secretome and the abolishment of sensory neurons firing rate in EV-depleted conditions. Overall, our results elucidate novel mechanisms on the peripheral nerves sprouting, essential for pursuing new targets for bone pain therapies.

## Introduction

Nerve fibers present in the bone microenvironment have been directly implicated in bone homeostasis and regeneration (Chenu, 2004; Chenu and Marenzana, 2005; Elefteriou, 2008; Elefteriou et al., 2014; Patel and Elefteriou, 2007). Anatomical mapping of innervation during skeletal development shows co-localization of nerve fibers with a rich capillary network in areas with high osteogenic activity (Gomez et al., 2005; Hukkanen et al., 1993; Li et al., 2007; Sisask et al., 2013; Tomlinson et al., 2016) and the pattern of neuronal dynamics observed during bone development is typically repeated (Hukkanen et al., 1993; Li et al., 2001; Madsen et al., 1998; Nordsletten et al., 1994), highlighting the critical role of neuronal regulation. Several denervation studies have revealed an impairment on fracture healing, emphasizing the impact of the peripheral nervous system on bone turnover (Bove et al., 2009; Cherruau et al., 1999; Edoff et al., 1997; Hill and Elde, 1991; Song et al., 2012). In line with these observations, sensory innervation has attracted significant interest concerning bone formation. Beyond their role in bone nociception, evidence demonstrates that the lack of sensory nerves significantly impairs bone mass accrual (Chen et al., 2019; Fukuda et al., 2013; Hayashi et al., 2012), indicating a key role in bone homeostasis. In the opposite way, undifferentiated bone marrow stromal cells are acknowledged to hold a neurotrophic potential. Both the bone marrow cells and/or their secretome are widely used to improve nerve regeneration (Brohlin et al., 2012; Chen et al., 2002; Gu et al., 2010; Han et al., 2016; Lu et al., 2005; Neuhuber et al., 2005; Novikova et al., 2011; Wilkins et al., 2009), mainly through the secretion of neurotrophic factors such as nerve growth factor (NGF), brain-derived neurotrophic factor (BDNF), vascular endothelial growth factor (VEGF), platelet-derived growth factor (PDGF) or epidermal growth factor (EGF) (Han et al., 2016). These growth factors are known to activate their respective receptors in axonal terminals, ultimatelly promoting axonal regeneration, elongation and guidance (Chao, 2003).

Interestingly, in bone pathological conditions such as bone cancer, ectopic sprouting of nerve fibers in the bone marrow, mineralized bone, and periosteum is detected (Bloom et al., 2011; Chartier et al., 2014; Ghilardi et al., 2010; Jimenez-Andrade et al., 2010). Osteoclasts, the bone resorbing cells, play a major role in the activation of nociceptors. Matrix resorption in the bone compartment results in active proton secretion into the extracellular space and a decrease in the pH milieu (Mantyh, 2014; Middlemiss et al., 2011; Yoneda et al., 2011), stimulating the acid-sensing ions channels in nerve endings and causing pain. Notably, clinical data showed that osteoclast inhibitors significantly reduced cancer-associated bone pain (Cleeland et al. 2013; Kohno et al., 2005). While such studies provide support for the role of osteoclasts in the activation of nociceptors in diseased bone (Yoneda et al., 2015b), recent reports start to reveal how osteoclasts contribute to the changes in the innervation pattern in normal or diseased bone. Zhu *et al*. have shown that osteoclasts can actively promote axonal growth on subchondral bone, through netrin-1 signaling, in an osteoarthritis model (Zhu et al., 2019). Moreover, osteoclast-derived netrin-1 was also linked to an increase in the sensory innervation of porous endplates of intervertebral disc, causing an low back pain (Ni et al., 2019). Therefore, deciphering the mechanisms by which peripheral innervation can be controlled by the osteoclasts constitute a major advantage for developing new strategies to tackle bone pain.

The current study has examined the effect of osteoclasts secretome on the growth and electrical signal propagation on sensory nerve fibers. We demonstrate new mechanisms by which peripheral innervation is controlled, namely indicating that the effects are dependent on the extracellular vesicles (EV) released by osteoclasts, through the epidermal growth factor receptor family targeting, by integrin independent pathways.

## Materials and methods

All animal procedures were approved by the IBMC/INEB ethics committee and by the Portuguese Agency for Animal Welfare (*Direção-Geral de Alimentação e Veterinária*) in accordance with the EU Directive (2010/63/EU) and Portuguese law (DL 113/2013). Mice were housed at 22°C with a 12 h light/ dark cycle with *ad libitum* access to water and food. Adult C57Bl/6 male mice (6-8 weeks-old) were sacrificed in a carbon dioxide chamber to obtain bone marrow cells, while pregnant females were sacrificed by cervical dislocation.

### Cell culture

#### Bone marrow cells cultures

Bone marrow stromal cells (BMSC) were isolated from tibiae and femur by flushing the bone marrow with α-MEM (Gibco, Thermo Fisher Scientific, Waltham, MA, USA) containing 10% (v/v) heat-inactivated (30 min at 56°C) foetal bovine serum (FBS, Gibco, Thermo Fisher Scientific) and 1% (v/v) penicillin/streptomycin (P/S, Gibco, Thermo Fisher Scientific). Non-adherent cells were removed after 3 days, and fresh medium was added. Cells were expanded from the colony-forming units for one week. Afterward, the medium conditioned for 3 days was collected, centrifuged at 140 g, 4°C, 5 min, and stored at −80°C. The conditioned medium was divided into small aliquots (500 μL - 1 mL) before freezing to avoid repeated freeze/thaw cycles.

#### Osteoclast cultures

Bone marrow cells were isolated from tibiae and femur by flushing the bone marrow with α-MEM containing 10% (v/v) FBS and 1% (v/v) P/S. To generate primary osteoclast precursors, the bone marrow mononuclear cell suspension was treated with red blood cells lysis buffer (ACK lysing buffer, #A1049201, Gibco, Thermo Fisher Scientific) for 1 min at room temperature (RT) and, after centrifugation, cells were plated in 10 cm diameter Petri dishes with 10 ng/mL macrophage colony-stimulating factor (M-CSF, PeproTech, London, UK) for 24 h. Afterward the M-CSF concentration was increased to 30 ng/mL for an additional 3 days (pre-osteoclasts). Conditioned medium from osteoclasts precursors was collected, centrifuged at 140 g, 4°C, 5 min, and stored at −80°C. Adherent cells were then detached with a cell scrapper and seeded at a density of 5 × 10^4^ cells/cm^2^ in the presence of 30 ng/mL M-CSF and 100 ng/mL receptor activator of nuclear factor kappa-B ligand (RANKL, PeproTech) (Bartell et al., 2014; Sousa et al., 2016). Cells were also seeded on top of bone slices (boneslices.com, Denmark) to collect the secretome from active resorbing osteoclasts. Cells were maintained at 37°C in a humidified atmosphere of 5% CO_2_ for 3-4 days and then osteoclast conditioned medium was collected and stored at −80°C. Evaluation of the osteoclast differentiation in supplemental data (Figure S1).

#### Dorsal root ganglia (DRG) culture

Embryonic DRG were obtained from 16 to 18 days-old (E16-18) murine embryos. Embryos were harvested and maintained on ice-cold Hank’s balanced salt solution (HBSS, Invitrogen). Embryonic ganglia were accessed through the dorsal side of the embryo after spinal cord removal. The meninges were cleaned from the isolated DRG, and the roots were cut. The ganglia were kept in cold HBSS until seeding. DRG were isolated under the stereoscopic magnifier and seeded into the lower wells of a 15-well μ-Slide Angiogenesis plate (#81506, Ibidi, Martinsried, Germany). Fibrin hydrogels were formed in the plates by applying equal volumes of a solution of plasminogen-free fibrinogen, pooled from human plasma, and a thrombin solution containing CaCl_2_ and aprotinin (final concentration of fibrin components: 6 mg/mL fibrinogen (Sigma-Aldrich); 2 NIH U/mL thrombin from human plasma (Sigma-Aldrich); 2.5 mM CaCl_2_ (Sigma-Aldrich); 10 μg/ mL aprotinin (Sigma-Aldrich)). Before being used in the preparation of fibrin gels, fibrinogen was dissolved in ultrapure water, dialyzed against tris-buffered saline (TBS, pH 7.4), sterile-filtered and diluted to 12 mg/mL with sterile TBS. The fibrin gel was allowed to polymerize for 15 min at 37°C in a 5% CO_2_ humidified incubator, before the addition of culture media. DRG were cultured with neurobasal medium supplemented with 2% v/v B-27 Serum-Free Supplement® (B-27, Invitrogen), 60 μM 5-fluoro-2’-deoxyuridine (FDU, Sigma-Aldrich), 25 mM glucose (Sigma-Aldrich), 1 mM pyruvate (Sigma-Aldrich), 50 ng/mL 7S NGF (Calbiochem, Merck Millipore), 2 mM glutamine (BioWittacker, Lonza, Basel, Switzerland) and 1% P/S. Embryonic DRG explant cultures were left undisturbed for 24 h. Afterward, neurobasal media was replaced by the conditioned media collected from BMSC and osteoclast cultures at different stages of differentiation (200 mg of total protein/well) for an additional three days. Controls with neurobasal supplemented with NGF and α-MEM supplemented with M-CSF and RANKL (no contact with cells) were performed.

### Quantification of axonal growth

Axonal outgrowth was quantified after 72 h of treatment with conditioned medium and controls. The embryonic DRG samples were fixed with 2% PFA in PBS for 10 min followed by 10 min at 37°C with 4% of PFA in PBS with 4% sucrose. Ganglia were permeabilized with 0.25% (v/v) Triton X-100 in PBS and incubated, for 30 min at RT, with blocking solution composed of 5% v/v normal goat serum (Invitrogen) and 5% v/v FBS in PBS. DRG cultures were incubated with an antibody directed against the neuronal-specific marker βIII tubulin (Promega, Madison, WI, United States) diluted 1:2000 in blocking solution, overnight at 4°C. Afterward, cells were washed and incubated for 1 h at RT with the secondary antibody (Alexafluor 488/568, Invitrogen). Images were captured either with a confocal laser scanning microscope (CLSM) Leica SP2 AOBS SE equipped with LCS 2.61 software (Leica Microsystems, Wetzlar, Germany) at the i3S Advanced Light Microscopy Unit or with an IN Cell Analyzer 2000 equipped with IN Cell Investigator software (GE Healthcare, Little Chalfont, UK) at the i3S BioSciences Screening Unit. Radial outgrowth, defined as the area between the ganglion edge and the outgrowth front, was determined. The outgrowth area was computed according to Bessa *et al.* (Bessa et al., 2013).

### Quantification of secretome-neurotrophins by enzyme-linked immunosorbent assay (ELISA)

Conditioned media from osteoclasts and BMSC were concentrated 10 times by centrifugation at 4000 g for 45 min, using 3 kDa MW cut-off filter units (Merck Millipore). In order to detect and quantify the amount of NGF, BDNF, neurotrophin-3 (NT-3) and NT4/5 in the conditioned medium, a multi-neurotrophin rapid screening ELISA kit (#BEK-2231, Tebu-bio, France) was used according to the manufacturers’ protocol. The concentration of netrin-1 in the concentrated conditioned media was measured using the ELISA development kit (EKC37454, Biomatik), also according to the manufacturer’s instructions.

### Analysis of mature osteoclast total secretome by information dependent acquisition (IDA) / liquid chromatography – mass spectrometry (LC-MS)

Protein identification was performed by nanoLC-MS/MS. This equipment comprises an Ultimate 3000 liquid chromatography system coupled to a Q-Exactive Hybrid Quadrupole-Orbitrap mass spectrometer (Thermo Scientific, Bremen, Germany). Samples were loaded onto a trapping cartridge (Acclaim PepMap C18 100Å, 5 mm x 300 μm i.d., 160454, Thermo Scientific) in a mobile phase of 2% ACN, 0.1% FA at 10 μL/min. After 3 min loading, the trap column was switched in-line to a 50 cm by 75μm ID EASY-Spray column (ES803, PepMap RSLC, C18, 2 μm, Thermo Scientific, Bremen, Germany) at 300 nL/min. Separation involved mixing A: 0.1% FA, and B: 80% ACN, with the following gradient: 5 min (2.5% B to 10% B), 120 min (10% B to 30% B), 20 min (30% B to 50% B), 5 min (50% B to 99% B) and 10 min (hold 99% B). Subsequently, the column was equilibrated with 2.5% B for 17 min. Data acquisition was controlled by Xcalibur 4.0 and Tune 2.9 software (Thermo Scientific, Bremen, Germany).

The mass spectrometer was operated in data-dependent (dd) positive acquisition mode alternating between a full scan (m/z 380-1580) and subsequent HCD MS/MS of the 10 most intense peaks from full scan (normalized collision energy of 27%). The ESI spray voltage was 1.9 kV. Global settings: use lock masses best (m/z 445.12003), lock mass injection Full MS, full-width half-maxima (FWHM) 15s. Full scan settings: 70k resolution (m/z 200), AGC target 3e6, maximum injection time 120 ms. dd settings: minimum AGC target 8e3, intensity threshold 7.3e4, charge exclusion: unassigned, 1, 8, >8, peptide match preferred, exclude isotopes on, dynamic exclusion 45 s. MS2 settings: microscans 1, resolution 35k (m/z 200), AGC target 2e5, maximum injection time 110 ms, isolation window 2.0 m/z, isolation offset 0.0 m/z, spectrum data type profile.

The raw data was processed using the Proteome Discoverer software (Thermo Scientific) and compared against the UniProt database for *Mus musculus* taxonomic selection. The Sequest HT search engine was used to identify tryptic peptides. The ion mass tolerance was 10 ppm for precursor ions and 0.02 Da for fragment ions. Maximum allowed missing cleavage sites was set to 2. Cysteine carbamidomethylation was defined as constant modification. Methionine oxidation and protein N-terminus acetylation were defined as variable modifications. Peptide confidence was set to high. The processing node Percolator was enabled with the following settings: maximum delta Cn 0.05; decoy database search target FDR 1%, validation based on q-value.

Network analysis was performed using the Cytoscape software (version v3.7.1) (Cytoscape Consortium, San Diego, CA, US)) with the plugins ClueGO (version v2.5.4) and CluePedia (version v1.5.4). We used ClueGO’s default settings: merge redundant groups with >50.0% overlap; the minimum GO level used was 0 and the maximum GO level was 2; statistical test used was “Enrichment/Depletion (Two-sided hypergeometric test)”, Bonferroni step down p-value correction; Kappa Score Threshold was 0.5; and number of genes was set at 3 with a minimum percentage at 4.0.

### Analysis of Phospho-Receptor Tyrosine Kinase (RTK) activation

A proteome profiler mouse phospho-RTK array kit (#ARY014, R&D system, Minneapolis, MN, USA) was used to quantify the phosphorylation level of 39 RTKs. After 72 h of treatment with conditioned medium and controls, protein lysate of DRG was quantified and analyzed. The protein content (from 6-10 DRG) of the same condition, from each independent experiment, was pooled together. According to the manufacturer’s instructions, for the array analysis, the same amount of protein was added to each membrane. Each array membrane was exposed to X-ray film using a chemiluminescence detection system (Amersham, GE Healthcare). The film was scanned using Molecular Imager GS800 calibrated densitometer (Bio-Rad, Hercules, CA, USA) and pixel density was quantified using Quantity One 1-D Analysis Software, v 4.6 (Bio-Rad). The results were presented as the mean spot intensity which corresponds to the mean of the two spots for each receptor within the same membrane array.

### Pharmacological inhibition of epidermal growth factor receptor (EGFR) and ErbB2

Embryonic DRG were cultured in 15-well slides for 24 h as previously described. Erlotinib, an EGFR inhibitor, was added to the conditioned medium at 10 nM, 100 nM, 1 μM, 10 μM and 100 μM and tested on DRG cultures during 72 h. Trastuzumab (Herceptin®), a monoclonal antibody against the ErbB2 receptor, was kindly donated by the Instituto Português de Oncologia do Porto (IPO, Porto, Portugal). Trastuzumab was tested in DRG cultures at a concentration of 10 μg/mL in conditioned medium. Afterward, axonal outgrowth was measured as previously described.

### β1 integrin subunit function blocking

To assess the contribution of the β1 integrin subunit on the phosphorylation of EGFR and osteoclast-induced axonal sprouting, stimulation of DRG with osteoclast secretome was conducted in the absence or presence of β1 integrin function-blocking antibodies, as previously described in (J. Silva et al., 2017). For this purpose, monoclonal antibodies against β1 (clone Ha2/5, BD Pharmigen; 10 μg/mL) integrin subunits were added to the osteoclast secretome and incubated with DRG at 37°C during 72 h. To evaluate the contribution of non-specific antibody interactions, incubation with isotype-matched control β1 antibodies (hamster IgM λ1, Pharmigen; 10 μg/mL) was also performed. At the end of this period, the wells were washed and fixed for immunocytochemistry or the protein collected for RTK phosphorylation array experiments.

### Extracellular vesicles (EV) depletion from the osteoclast secretome and characterization

Osteoclasts were isolated and differentiated as described in previous sections. After seeding on 48 well plates, cells were cultured with 10% EV-depleted FBS. 24h prior media collection, FBS was reduced to 1%. All steps for the EV depletion were conducted under sterile conditions and in line with the published *Minimal information for studies of extracellular vesicles 2018* guidelines (Théry et al., 2018) and as described in (Théry et al., 2006). Briefly, the secretome was collected, centrifuged 1 000g 10 min to clear the cell debris, 2 000g for 10 min, followed by 10 000g for 30 min. The supernatant was then centrifuged at 100 000g using 70Ti rotor (Beckman Coulter Genomics), for 120 min. The pellet containing exosomes was then resuspended in PBS and stored at −80°C. All centrifugation steps were performed at 4°C (Tassew et al., 2017). The supernatant was stored −80°C to perform the experiments comprising the EV-depleted osteoclast secretome.

Western blot (WB) analysis was performed to the protein content from the EV enriched fraction. Protein was quantified by DC Protein Assay kit (Bio-Rad). The same amount of protein (15 μg) of the EV enriched fraction and the corresponding supernatant were prepared in non-reducing loading buffer, denatured at 95°C for 5 min, and loaded in 10% polyacrylamide SDS-PAGE gels. Resolved proteins were wet-transferred to nitrocellulose membranes, and membranes blocked with non-fat dry milk 5% solution. Membranes were probed overnight at 4°C with hamster anti-mouse CD81 antibody, clone Eat2 (MCA1846GA, Bio-Rad). Membranes were probed with HRP-conjugated secondary antibody (GE Healthcare), incubated with ECL substrate, and chemiluminescence signal was detected with autoradiographic films (all from GE Healthcare). Films were scanned on a GS −800 calibrated imaging densitometer (Bio-Rad).

The osteoclast-derived EV enriched fraction was further characterized by nanoparticle tracking analysis (NTA) and transmission electron microscopy (TEM), as previously described (A. M. Silva et al., 2017). Briefly, for size and particle concentration evaluation, EV suspensions were diluted 1:500 in filtered PBS 1x and analyzed by NTA in a NanoSight NS300 device with NTA3.0 software. For the TEM negative staining, 10 μL of samples were mounted on Formvar/carbon film-coated mesh nickel grids (Electron Microscopy Sciences, Hatfield, PA, USA) and left standing for 2 min. The liquid in excess was removed with filter paper, and 10 μL of 1% uranyl acetate were added on to the grids and left standing for 10 s, after which, liquid in excess was removed with filter paper. Visualization was carried out on a JEOL JEM 1400 TEM at 120 kV (Tokyo, Japan). Images were digitally recorded using a CCD digital camera Orious 1100W (Tokyo, Japan) at the i3S Scientific Platforms Histology and Electron Microscopy.

### Microelectrode–microfluidic cultures and electrophysiology recordings

The microfluidic devices were fabricated by mixing the polydimethylsiloxane (PDMS) elastomer (Sylgard® 184, DowCorning) with curing agent (10:1, w/w), degassed and cured over the mold at 65°C for 2 h. Custom microElectrode-microFluidic (μEF) devices were prepared as described previously (Lopes et al., 2018). Briefly, PDMS microfluidic chambers were aligned on top of microelectrode array (MEA) chips (MultiChannel Systems MCS GmbH, Germany), with 252 recording electrodes (30 μm in diameter and pitch of 100 μm) organized in a 16×16 grid. Microfluidic chambers had an appropriate microchannel spacing for compartmentalization and probing of axonal activity. Microfluidic chambers were also adapted by adding an extra smaller reservoir (Ø 3 mm), which allowed the seeding of the DRG in a central position to the electrode matrix (Neto et al., 2014). μEF devices were composed of two separate compartments connected by 16 microchannels of 700 μm length × 9.6 μm height × 10 μm width. Each microchannel was probed with 7 electrodes, thus, every axon extending to the axonal compartment was electrophysiologically monitored. After mounting, μEFs were sequentially coated with poly-D-lysine (PDL) (0.01 mg/ml) (Corning) and laminin (0.01 mg/ml) (Sigma-Aldrich). The unbound laminin was aspirated, and chambers were refilled with complete neurobasal medium and left to equilibrate at 37 °C. Isolated embryonic DRG explants were placed and cultured as described before. DRG explants extended axons to the axonal compartment within the first 5 days *in vitro* (DIV). Then, treatments and recordings were performed at 6-7 DIV. This time point was chosen following preliminary studies that showed adequate electrophysiological maturation and culture viability at this stage (Heiney et al., 2019).

Besides the positive control composed of complete neurobasal supplemented with NGF, three conditions with osteoclast secretome were tested. The total mature osteoclast secretome, the EV-depleted osteoclast secretome and secretome from osteoclasts cultured in mineralized substrates.

Recordings at a sampling rate of 20 kHz were performed using a MEA2100 recording system (MCS GmbH, Germany). In every recording session, the temperature was maintained at 37ºC by an external temperature controller. After removing the cultures from the incubator, recordings only started after 5 minutes of habituation to avoid an effect due to mechanical perturbation. Then, a baseline recording (5 minutes) was obtained. Afterward, the medium from the axonal compartment was gently aspirated and replaced by 100 μl of treatment medium. The larger volume present on the somal compartment maintained a hydrodynamic pressure difference, inhibiting any flow from the axonal to the somal compartment.

Post-treatment (timepoint 0h) recordings (20-30 minutes) were started as soon as the baseline stabilized following liquid flow perturbation (less than 1 minute). Cultures were returned to the incubator until 3 hours after treatment, when a post-treatment recording (timepoint 3h) of 5-10 minutes was performed. The day after, a final post-treatment recording (timepoint 24h) was performed for 5-10 minutes.

Raw signals were high-pass filtered (200 Hz) and spikes were detected by a threshold set to 5× SD of the electrode noise. Spike data analysis was carried out in MATLAB R2018a (The MathWorks Inc., USA) using custom scripts. The mean firing rate (MFR) of each microchannel was calculated by averaging the MFR of the 5 inner electrodes (typically electrode rows 10-14), due to their superior signal-to-noise ratio. Microchannels with an MFR of at least 0.1 Hz in a given time point at 6 DIV were considered as active and included in the analysis.

### Statistical analysis

All experiments were run in triplicate and repeated at least three times. Data analysis was performed using GraphPad Prism 8.20 for Windows (GraphPad Software, San Diego CA, USA). Statistical differences between groups were calculated using one-way analysis of variance, more precisely, the non-parametric Kruskal-Wallis test followed by Dunns post test for multiple comparisons. The non-parametric Mann-Whitney t-test was used to identify statistical differences when only two groups where being compared. Differences between groups were considered statistically significant when *0.01 < p < 0.05, ** 0.001 < p < 0.01, *** p < 0.001, **** p < 0.0001.

## Results

### Sensory neurons outgrowth under osteoclasts effect is neurotrophin-independent

To understand the response of peripheral sensory nerves to the cells present in the bone compartment we assessed the effect of the secretome from osteoclasts lineage, at different stages of differentiation, in axonal growth and sprouting of dorsal root ganglia (DRG). We compared these effects with the neurotrophic effect of bone marrow stromal cells (BMSC).

Organotypic explants of embryonic DRG cultures were exposed to the secretome derived from BMSC, osteoclast precursors (pre-OC) and mature osteoclasts (OC) populations. The mature osteoclast secretome demonstrated an approximate 3-fold stronger influence on sensory neurons growth than the secretome from BMSC (Figure 1A-B), previously demonstrated to have a positive neurotrophic potential (Gu et al., 2012, 2010). The DRG treated with osteoclastic lineage conditioned media, independently of the osteoclasts maturation status (pre-OC and mature OC), showed significantly higher axonal network when compared to the neurobasal-nerve growth factor (neurobasal (+NGF)) positive control (Figure 1A-B). Explant DRG cultures were also treated with osteoclast differentiation medium (OCm) to rule out the effect of receptor activator of nuclear factor kappa-Β ligand (RANKL) and macrophage colony-stimulating factor (M-CSF) present in the medium and known to modulate axonal outgrowth (Gutierrez et al., 2013; Imai and Kohsaka, 2002). No significant differences were observed when compared to neurobasal control media.

**Figure 1:**
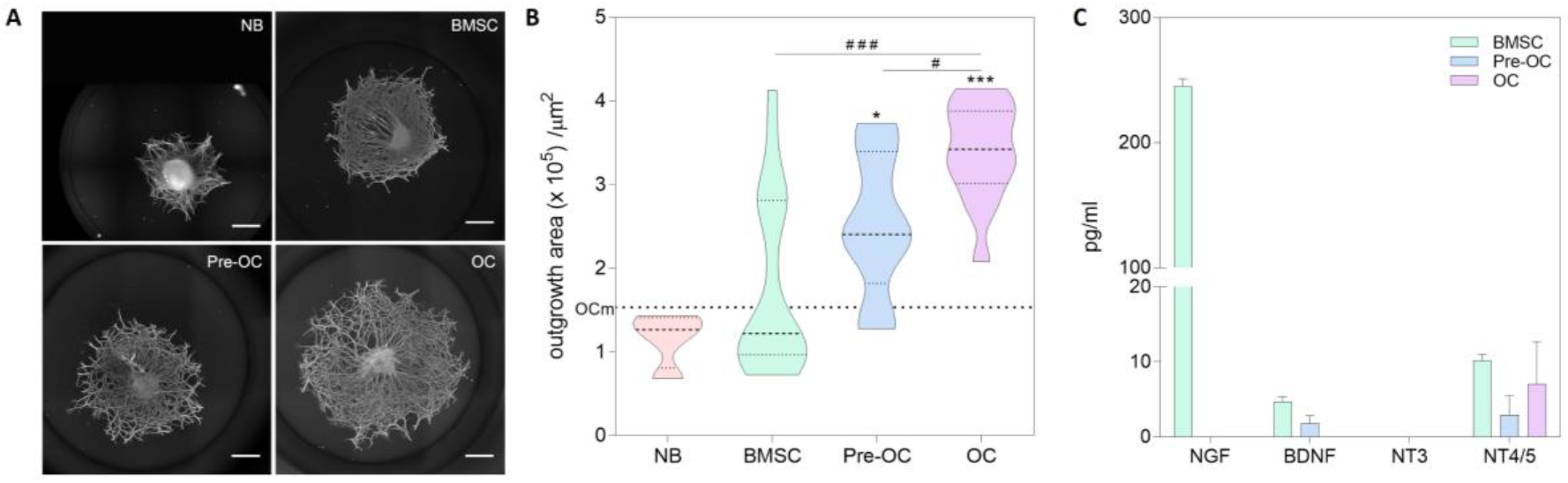
Sensory neurons axonal outgrowth under the effect of bone cells secretome. **A.** Representative images of DRG outgrowth after treatment (stained for βIII-tubulin, scale bar 500 μm). Neurobasal supplemented with nerve growth factor (NB); bone marrow stromal cells conditioned medium (BMSC); Pre-osteoclast conditioned media (Pre-OC), and mature osteoclast conditioned media (OC) was collected and added to embryonic DRG culture. **B.** Automatic axonal outgrowth quantification of DRG. Data represented as a violin plot. Non-conditioned osteoclast medium (OCm) represented by the dashed line. Statistical analysis was performed *vs.* controls – NB and OCm (*) and between groups (#). **C.** Analysis of secretome neurotrophins. Quantification of NGF, BDNF, NT-3, and NT-4/5 by ELISA on BMSC, Pre-OC and mature OC conditioned media. Data represented as mean±SD.

BMSC are known to promote axon growth through the secretion of neurotrophins (Cantinieaux et al., 2013; Gu et al., 2012, 2010) and osteoclasts where described to induce nerve outgrowth through netrin-1 action (Ni et al., 2019; Zhu et al., 2019). To assess if the osteoclast secretome observed effect follow similar mechanisms, we measured in the conditioned medium the levels of NGF, BDNF, NT-3, NT-4/5 and netrin-1, by enzyme-linked immunosorbent assay (ELISA). NGF, BDNF, and NT-3 were not detected in the mature osteoclast secretome, despite our observation of higher neurite outgrowth on embryonic DRG explant cultures. NT-4/5 was detected in very small concentrations for the three different secretomes, while NGF and BDNF were detected in the BMSC secretome (Figure 1C). Netrin-1 was not detected in the different secretomes (data not shown).

### Epidermal growth factor receptor family signaling pathway is involved in the axonal sprouting of DRG organotypic cultures under osteoclasts secretome stimulation

To understand the mechanisms activated in the context of axonal outgrowth under osteoclasts stimulation, we determined the phosphorylation level (and therefore, activation level) of receptors tyrosine kinase (RTK) which have been implicated in neuronal development, growth, survival and axonal regeneration (Vigneswara et al., 2012).

Epidermal growth factor receptor (EGFR), ErbB2, and platelet-derived growth factor receptor alpha (PDGFRα) displayed higher activation levels (Figure 2). Interestingly, under osteoclast stimulation, there was increased activation of the ErbB2 receptor in DRG neurons when compared to treatment with BMSC (Figure 2B). ErbB2, an orphan receptor with no characterized ligand, can be activated by spontaneous homodimer formation or by heterodimerization with another ligand-bound or epidermal growth factor (EGF) family transactivated receptor (Macdonald-Obermann et al., 2013; Olayioye, 2000). The high levels of EGFR phosphorylation suggest heterodimerization of EGFR/ErbB2 leading to pathway activation. Low levels of TrkA, TrkB and TrkC phosphorylation were observed further confirming the low contribution of NGF, BDNF, NT-3 and NT-4/5 neurotrophins (Figure 2A). In the context of nerve repair, these results are in agreement with the literature where ErbB receptors expression was shown to be increased in DRG upon lesion (Mizobuchi and Kanzaki, 2013). Therefore, osteoclasts might promote axonal outgrowth through EGFR family signaling, described to be involved in neuronal repair.

**Figure 2:**
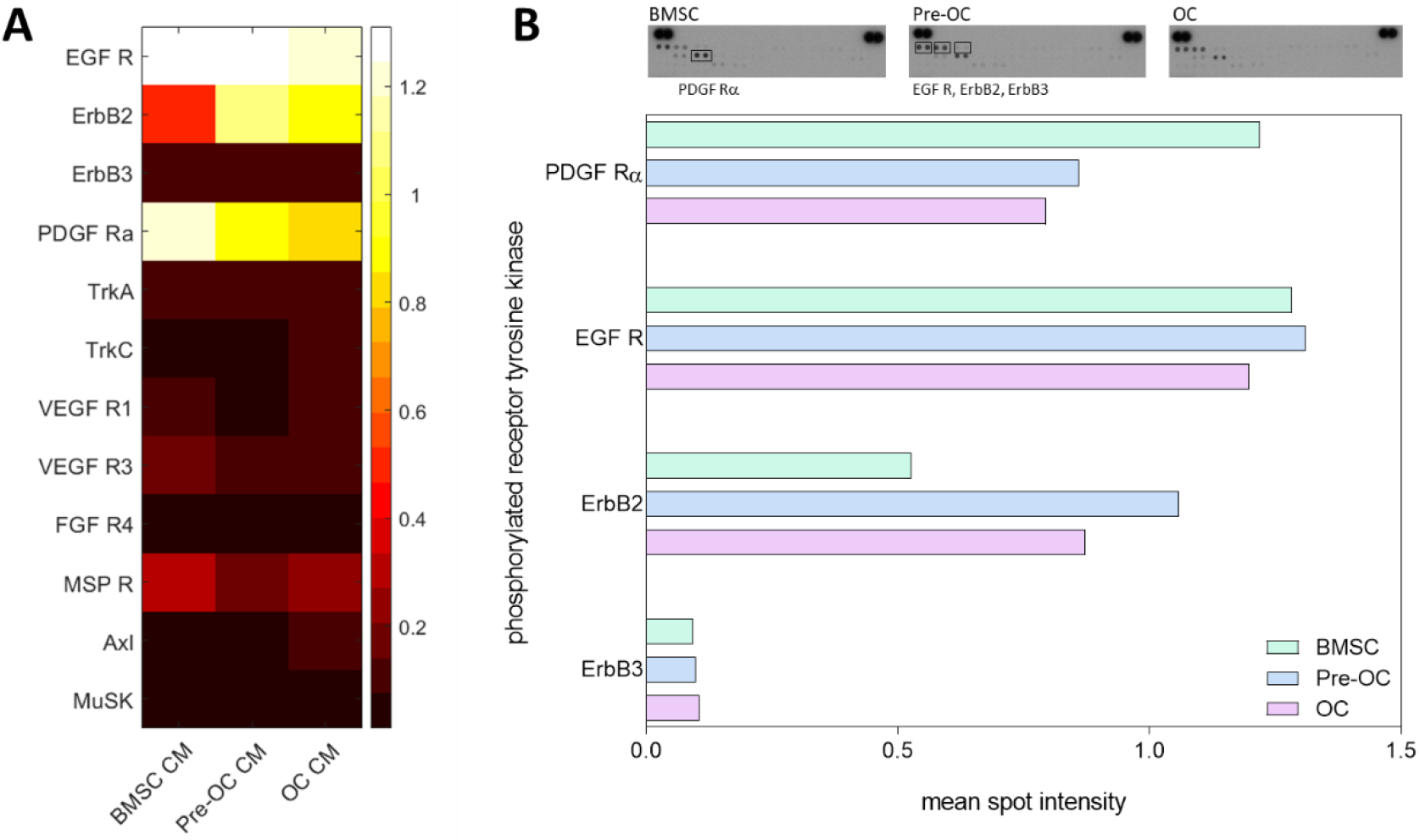
Analysis of activated receptor tyrosine kinases (RTK) in DRG cultures. **A.** Phosphorylation levels of RTK on DRG after treatment with with different secretomes – BMSC; Pre-OC and mature OC. Heatmap representing the relative spot intensity for the activated receptors calculated from the pixel density in the X-ray films. **B.** X-ray films of the arrays performed. The graphic represents the mean spot intensity of the most phosphorylated receptors in embryonic DRG treated with different secretomes – BMSC; Pre-OC and mature OC, showing the major activation of two different families: epidermal-growth factor receptors (EGFR), platelet-derived growth factor (PDGF) receptor alpha. For array analysis, protein lysate from a 6–10 DRG in each condition, from three independent experiments (n = 3), was pooled. One array was used per condition.

To understand the contribution of EGFR and ErbB2 signaling in osteoclast-induced axonal outgrowth, receptor-mediated inhibition using pharmacological blockers was performed (Figure 3).

**Figure 3:**
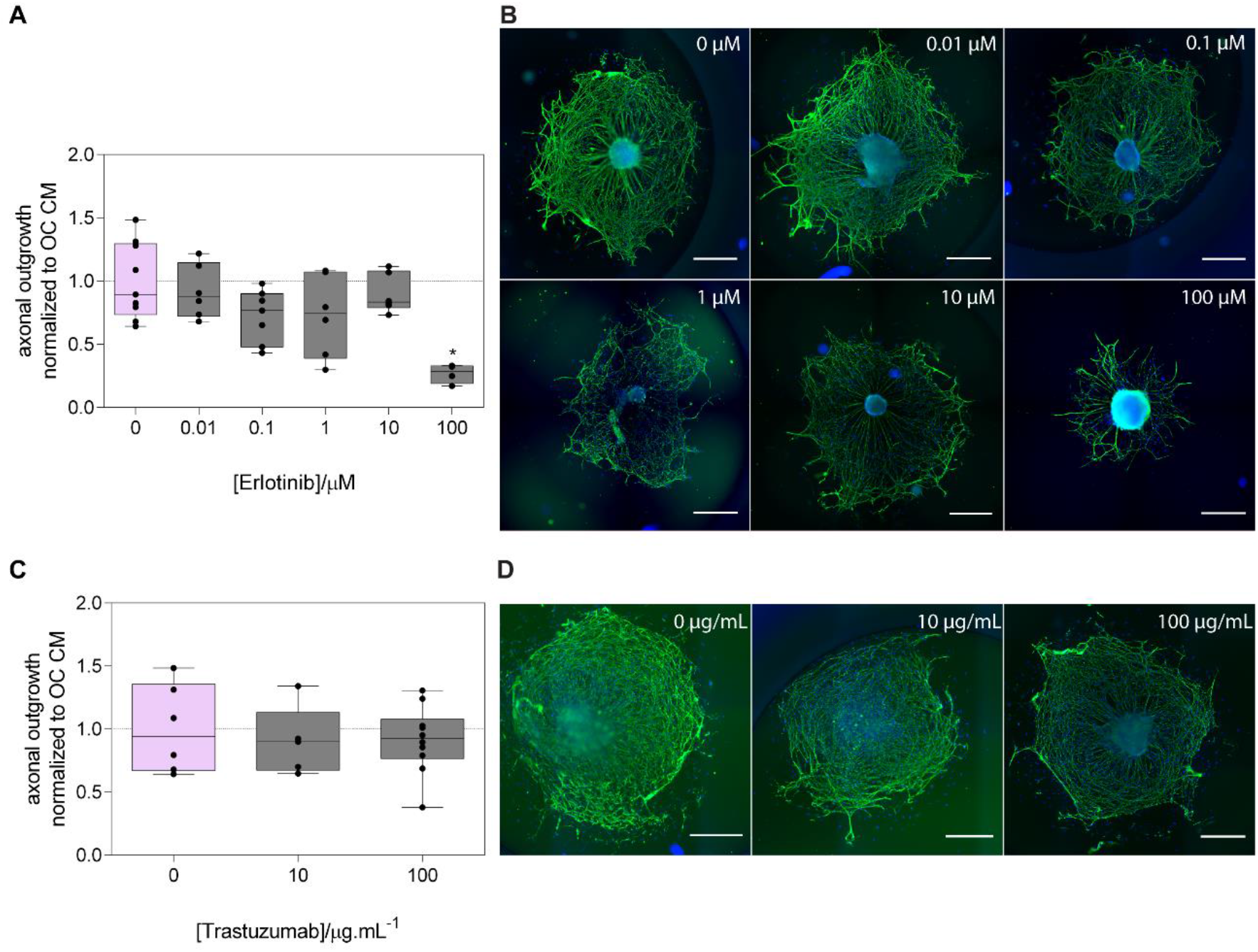
Pharmacological inhibition of EGFR and ErbB2. **A.** Quantification of axonal outgrowth of sensory neurons blocked with epidermal growth factor receptor (EGFR) inhibitor – Erlotinib. **B.** Representative images of DRG treated with different concentrations of Erlotinib (βIII tubulin in green and nuclei in blue, scale bar 500 μm). **C.** Quantification of axonal outgrowth of sensory neurons blocked with ErbB2 receptor inhibitor – Trastuzumab. **D.** Representative images of DRG treated with different concentrations of Trastuzumab (βIII tubulin in green and nuclei in blue, scale bar 500 μm). Data represented as box & whiskers (centerline is plotted at the median. Whiskers represent minimum to maximum range) *p≤0.05.

Erlotinib is an EGFR inhibitor that reversibly binds to the intracellular tyrosine kinase domain of the receptor, blocking EGFR over ErbB2. At high concentrations it inhibits both receptor kinases (Hynes and Lane, 2005; Macdonald-Obermann et al., 2013; Wood et al., 2004). Our results show that the neurotrophic effect of osteoclasts was reduced in the presence of the high Erlotinib concentration (Figure 3A-B), without compromising the cell viability and metabolic activity (supplemental data Figure S2), suggesting that both EGFR/EGFR homodimers and EGFR/ErbB2 heterodimers might contribute to the osteoclast-mediated effect in axonal outgrowth.

To unravel the input of ErbB2 in osteoclast neurotrophic effect the monoclonal antibody Trastuzumab (also known as Herceptin®) was used against the extracellular domain (Nielsen et al., 2013). Upon treatment with Trastuzumab, no differences in the outgrowth area of the sensory neurons were observed (Figure 3C-D), indicating that sole inhibition of ErbB2 signaling fails to circumvent the osteoclast secretome stimuli. These results show that osteoclast-mediated axonal outgrowth required both EGFR- and ErbB2 receptor-dependent pathways.

### EGFR phosphorylation is independent of β1 integrin subunit activation

Integrin-mediated adhesion can enhance signaling pathways by direct phosphorylation of growth factor receptors. β1 integrin cytoplasmic domain was demonstrated to trigger integrin-dependent EGFR phosphorylation (Moro et al., 2002). Therefore, we decided to explore further if β1 integrin activation could contribute to EGFR family phosphorylation and synergistically potentiate axonal growth.

An antibody-based β1 integrin subunit blocker and the respective isotype were added to the osteoclast secretome, and axonal sprouting was measured. No differences in the axonal outgrowth was observed when the anti-β1 integrin isotype was added to the osteoclast secretome (supplemental Figure S3). Regarding the functional blocker, the treatment induced a significant decrease in the axonal outgrowth area when compared to the control (Figure 4A-B). Nevertheless, no alterations of EGFR family phosphorylation were observed when the activity of β1 integrin was reduced (Figure 4C). These results were further supported by immunocytochemistry where phosphorylated EGFR was detected independently on the condition analyzed (Figure 4D).

**Figure 4:**
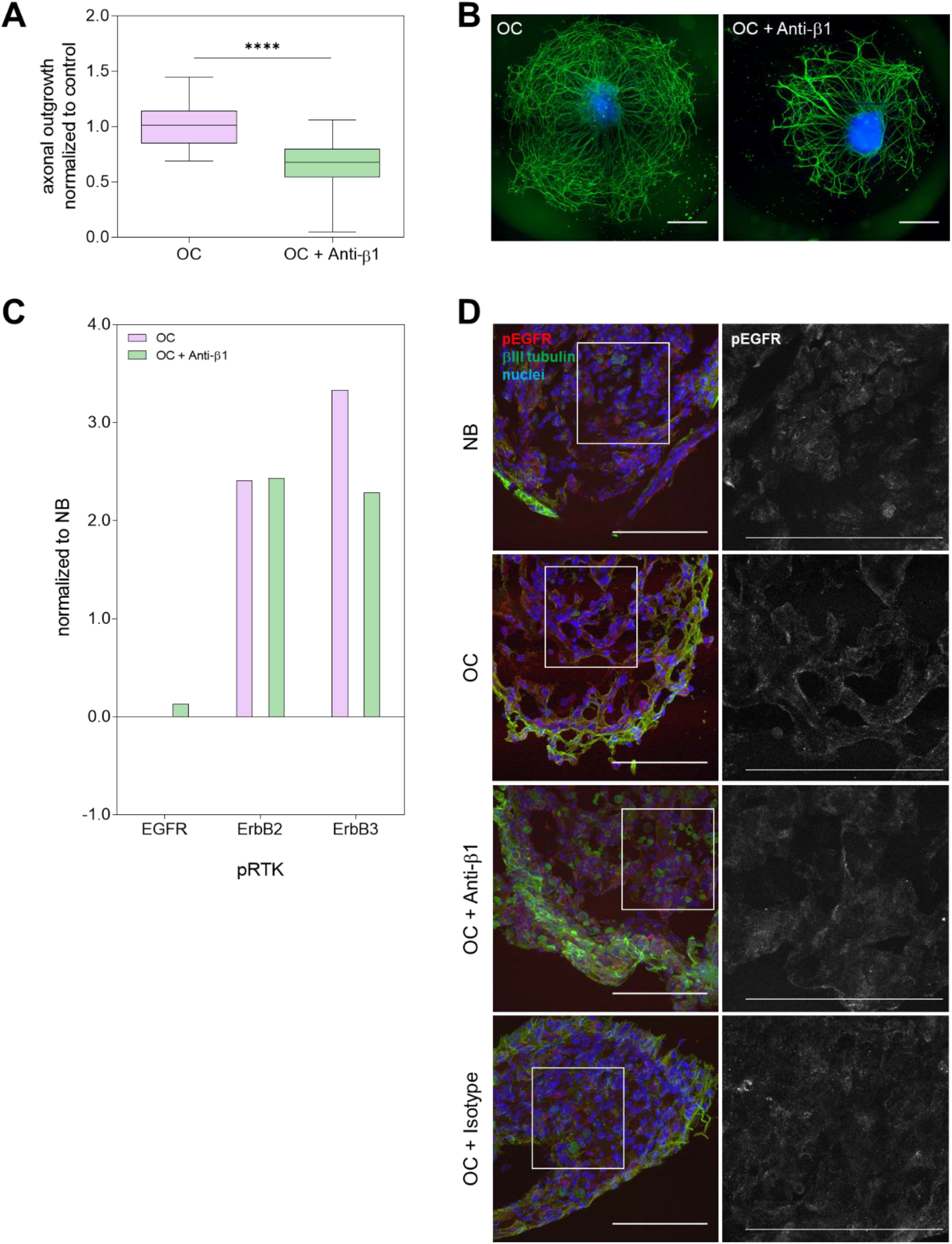
Effect of β1 integrin blocker in axonal sprouting and EGFR phosphorylation. **A.** Quantification of axonal sprouting of DRG stimulated with osteoclast secretome and osteoclast secretome with the functional anti-β1 integrin. Data represented as box & whiskers (median, whiskers represent minimum to maximum range), ****p≤0.0001. **B.** Representative images of DRG treated with osteoclast secretome and osteoclast secretome with the functional anti-β1 integrin. Staining for βIII tubulin in green and nuclei in blue, scale bar 500 μm. **C.** Levels of EGFR family phosphorylation assessed by RTK array. Values normalized by the neurobasal control. **D.** Representative images of EGFR phosphorylation by immunocytochemistry after treatment with NGF-supplemented neurobasal, osteoclast secretome, osteoclast secretome with the functional anti-β1 integrin and osteoclast secretome with anti-β1 integrin isotype. Phosphorylated EGFR (pEGFR) in red, βIII tubulin in green and nuclei in blue (left column); zoom in images of the isolated channel for the pEGFR (right column). Scale bar 100 μm.

### Soluble EGFR/ErbB2 activators were undetected in the osteoclast-derived secretome

EGF and TGFα ligands, that preferentially bind to the EGFR homodimers and EGFR/ErbB2 heterodimers (Hynes and Lane, 2005), activating the signaling cascade, were not detected in the osteoclasts secretome by ELISA (data not shown). Therefore, to understand which ligands were present in the osteoclast secretome that could directly or indirectly activate the signaling cascade, we proceed to a higher sensitive technique and performed proteomic analysis by liquid chromatography-tandem mass spectrometry (LC-MS/MS) of osteoclast serum-free secretome collected at two different time points: 6 h and 24 h.

Over four hundred proteins were identified in the osteoclast serum-free secretome. The complete set of identified proteins is available as supplemental data (supplemental Table S1). Protein identification revealed oscillations in the relative concentration of the soluble factors throughout the conditioning period. As expected, the higher amount of the identified proteins at 24 h were increased in relative concentration when compared to 6 h.

Interestingly, the identified proteins were not only assigned to the extracellular region (extracellular matrix proteins, secreted soluble factors, extracellular vesicles-associated proteins, etc.) but also to synaptic part (synaptic structural proteins, downstream effectors, cytoskeletal components) in equal percentage, according to Gene Ontology cellular component terms (ClueGO v2.5.4 + CluePedia (v1.5.4) from the software Cytoscape (v3.7.1) (Figure 5A). By restricting the analysis to pathways with a higher significance (p<0.01) and increasing the specificity level of the cellular component terms, we observed that the majority of the proteins remained associated to the initial detected clusters (Figure 5B and supplemental table S2). Importantly, neither EGF nor TGFα were identified among the secreted proteins, which is in accordance with a previous study reporting briefly the analysis of the osteoclast secretome where no EGFR ligands were detected (Rody et al., 2017). Nevertheless, our thorough proteomic analysis of the secretome revealed that the osteoclasts secrete molecules to interact with neuronal and non-neuronal cells to directly influence neuronal processes. Dihydropyrimidinase-related protein 2 (Dpysl2), also known as collapsin response mediator protein 2 (Crmp2), here found linked to the synaptic cluster. Moreover, in the extracellular region cluster, we detected the protein delta homolog 1 (Dlk1) and Serpin F1 (SerpinF1), also known as pigment epithelium-derived factor (PEDF). Still, none of the identified proteins are described to contribute directly or indirectly to the EGFR/ErbB2 signaling cascade.

**Figure 5:**
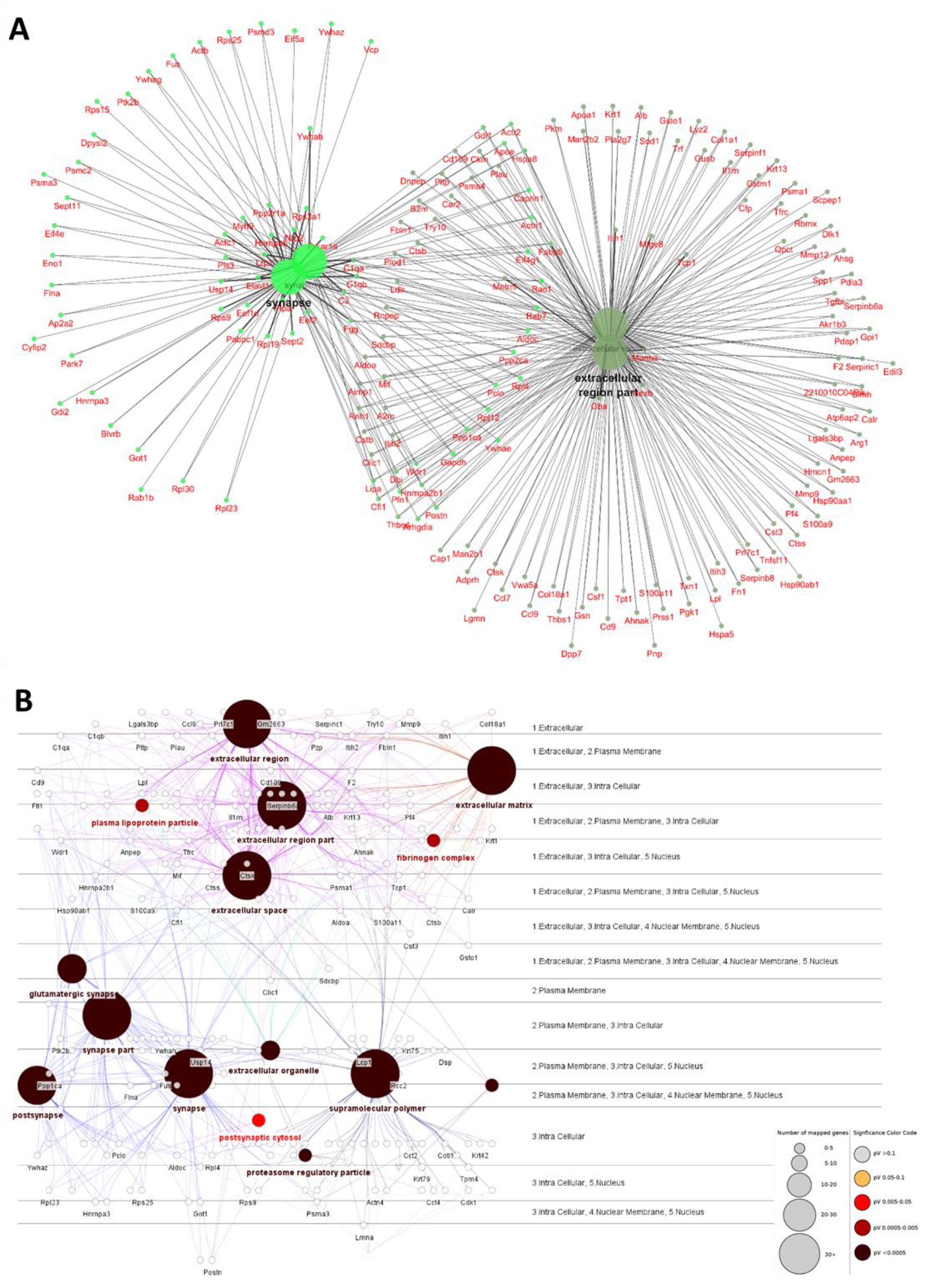
Cytoscape network of the main cellular components associated to osteoclasts-secreted proteins at 6 h and 24 h. **A.** Network analysis using the plugin ClueGO (v2.5.4) + CluePedia (v1.5.4) from the software Cytoscape (v3.7.1) to analyze the cellular components represented by the identified proteins from osteoclast secretome. Gene ontology (GO) term fusion was applied. Pathways with p value lower than 0.05 were considered. Tree interval was applied from 0 to 1 level. Genes from the different cellular components are shown in red. **B.** Protein network (cerebral view layout) representing the most significant clusters associated with the identified proteins. Gene ontology (GO) term fusion was applied. Pathways with p value lower than 0.01 were considered. Tree interval was applied from 1 to 2 level. The size of the nodes is indicative of the number of associated proteins. The increase in red color gradient represents higher statistical significance.

### Extracellular vesicles secreted by osteoclasts carry mediators of the EGFR/ErbB2-dependent axonal sprouting

Neurotrophic proteins, are generally believed to be released from cells in a soluble form and act locally or move through the extracellular milieu, which lead us to perform the proteomic profiling in the total secretome. However, it is increasingly appreciated that growth factors can be released from cells in or on the surface of extracellular vesicles (EV)/exosomes (Jaiswal and Sedger, 2019). We, therefore, hypothesized that osteoclast-derived EV could play a crucial role in the axonal outgrowth. To test this, DRG were exposed to EV-depleted osteoclast secretome and the axonal sprouting was measured. The EV enriched fraction was characterized by NTA, WB and by TEM. The analysis of the size and concentration of the vesicles showed a normal distribution with a mean size of 141.8 ± 2.7 nm and a concentration of 4.90 × 10^11^ particles/ml (supplemental Figure S4). The analyzed fraction stained positive for the CD81 EV-specific marker (Figure 6A) and by negative staining for TEM, it was possible to visualize the EV present in the OC-derived secretome (Figure 6B, white arrowheads), ranging from 40 to 200 nm.

**Figure 6:**
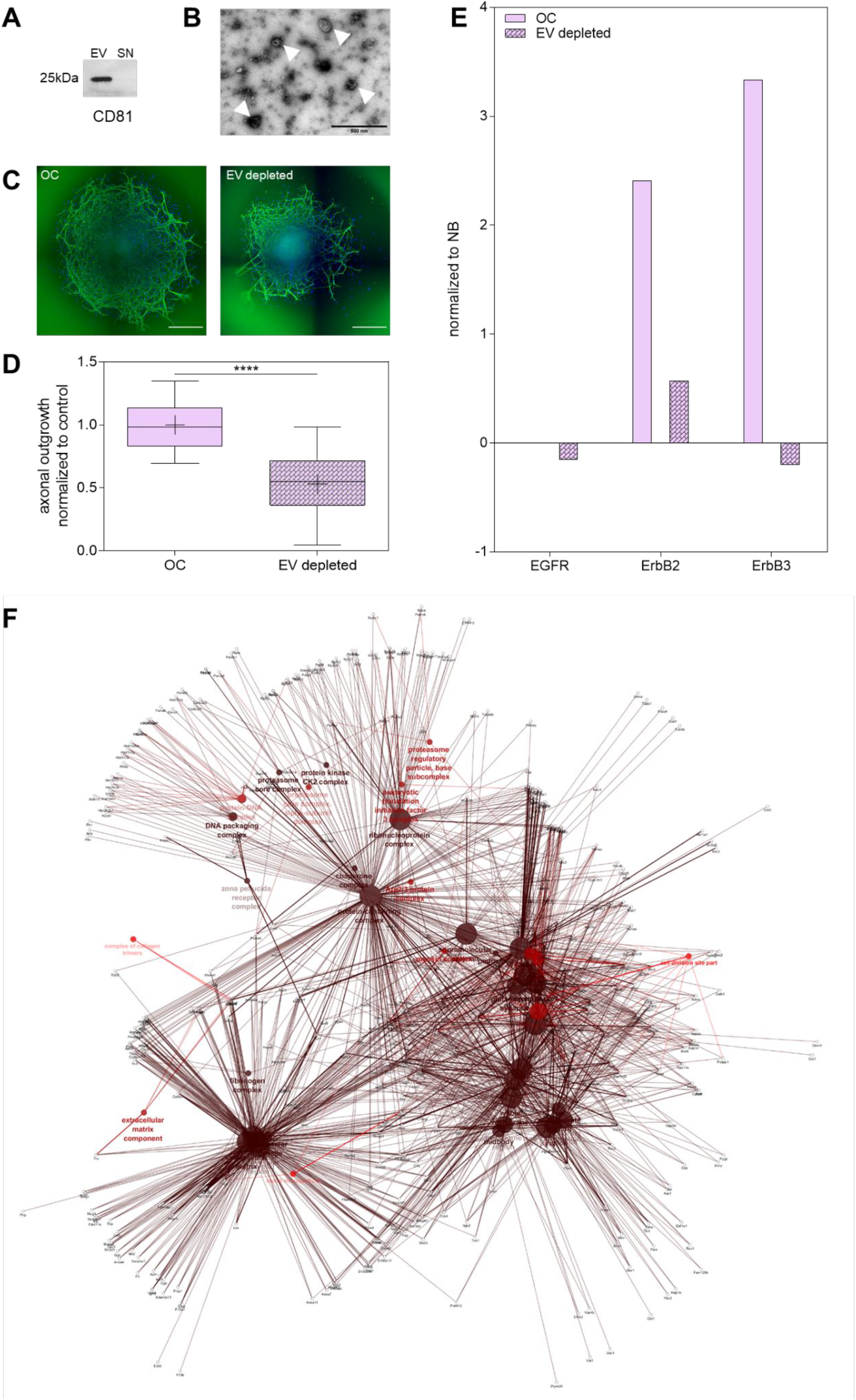
Dorsal root ganglia (DRG) axonal network growth is dependent on the osteoclasts extracellular vesicles (EV). **A.** Characterization of osteoclasts-derived EV by Western blot using CD81 membrane marker (EV *vs.* Supernatant (SN) EV depleted). **B.** Transmission electron microscopy of osteoclasts-derived EV (white arrows) by negative staining (100 000x). **C.** Representative images of DRG treated with osteoclast secretome and EV-depleted osteoclast secretome. Staining for βIII tubulin in green and nuclei in blue, scale bar 500 μm. **D.** Quantification of axonal sprouting of DRG stimulated with osteoclast secretome vs. osteoclast secretome lacking the EV. Data represented as box & whiskers (median, whiskers represent minimum to maximum range), *p≤0.05. **E.** Levels of RTK phosphorylation assessed by RTK array. Values normalized to the neurobasal control. **F.** Network analysis of osteoclast-derived EV enriched fraction according to the cellular components. Protein network (edge-weighted spring embedded layout) representing the most significant clusters associated with the identified proteins. Gene ontology (GO) term fusion was applied. Pathways with p value lower than 0.01 were considered. Tree interval was applied from 1 to 2 level. The size of the nodes is indicative of the number of associated proteins. The increase in red color gradient represents higher statistical significance. High resolution networks provided as supplemental material.

A significant decrease in axonal sprouting was observed in the absence of EV (Figure 6C-D). This suggests that the EV cargo plays a role in the neurotrophic potential of the osteoclast secretome. To understand if EV depletion modulates the EGFR/ErbB2 phosphorylation levels, that we previously demonstrated to be involved in the osteoclast-induced axonal outgrowth, we evaluated the phosphorylation state of the EGFR family. Remarkably, a substantial decrease in EGFR family phosphorylation was observed in the absence of osteoclast-derived EV (Figure 6E), confirming that the EGFR family activator is present in the EV cargo and reinforcing the contribution of this signaling pathway to the osteoclast-mediated axonal growth.

Therefore, the EV enriched fraction was further analyzed by LC-MS/MS for the identification of potential proteins that can directly or indirectly regulate the EGFR/ErbB2 signaling cascade. Over five hundred proteins were identified (supplemental table S3). New clusters were detected, when comparing to the previous analysis performed for the whole secretome. Highly significant clusters containing proteins related to DNA binding, cell-substrate interactions, proteic complexes and membrane-associated proteins were revealed (supplemental table S4). Very interestingly, the nodes with higher protein contribution remained the synaptic part (27%) and extracellular region (15%). In addition, according to the proteomic analysis, SH3 domain-binding glutamic acid-rich-like protein (SH3BGRL) previously described to interact with the EGFR signalling cascade (Schulze et al., 2005; Taguchi et al., 2011) and promote the dowstream activation of c-Src (Wang et al., 2016) was detected. The proteins present in the nodes with SH3BGRL were closely related to several biological processes including developmental growth with high contribution to positive regulation of cell growth (Figure 6F and supplemental table S5).

### Electrical activity induced by osteoclast secretome exposure is eradicated upon EV depletion

To unravel the electrophysiological implications of the axonal exposure to the osteoclast secretome, a combination of substrate-integrated microelectrode arrays (MEAs) with custom-made microfluidic chambers (Lopes et al., 2018) were used. MEAs enable non-invasive, thus repeatable, recordings of extracellular action potentials. Although recordings of random DRG cultures have been previously demonstrated (Black et al., 2019), these are not adapted to the study of peripheral innervation. Here, we employed microElectrode-microFluidic (μEF) devices (Figure 7A), which allowed us to monitor axonal activity with great fidelity (Figure 7B) (Heiney et al., 2019; Lopes et al., 2018).

**Figure 7:**
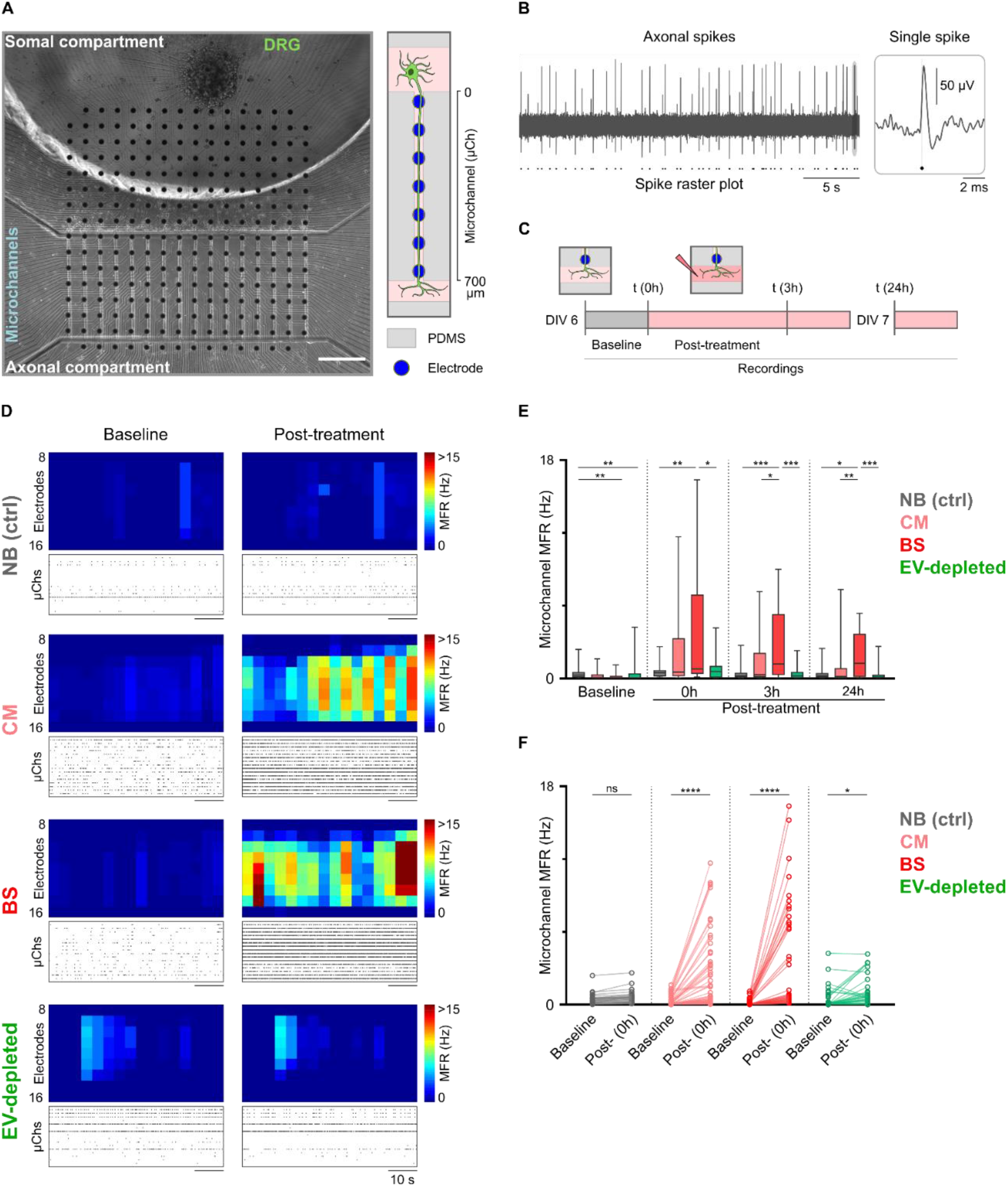
Electrophysiology studies on dorsal root ganglia (DRG) neurons in microfluidic devices stimulated with osteoclasts secretome. **A.** Phase-contrast microscopy image mosaic of an organotypic dorsal root ganglia (DRG) culture at 7 days *in vitro* (DIV). The whole microelectrode array (MEA) (1.5 × 1.5 mm) active area is shown by a combination of 9 mosaic images (10× objective) from different parts of the culture. A PDMS device composed of 16 microchannels (10 μm width; 700 μm length) is aligned to encompass 7 microelectrodes. Details of axonal morphology can be seen in the somal compartment, microchannels and axonal compartment (scale bar = 200 μm). A schematized version of a microchannel is shown on the right. **B.** Electrophysiological trace of 30 seconds of baseline activity from an electrode (within a microchannel) at 6 DIV and corresponding spike raster plot. Inset shows a single spike. **C.** Schematic diagram of experiments and timeline of recordings. **D.** Representative activity maps of baseline and post-treatment (time 0h) activity for each condition (microchannel area only, electrode rows 9-15). Each pixel corresponds to an electrode and the mean firing rate (MFR) is color-coded. Corresponding raster plots of 60 seconds of activity are shown below each activity map. Each row corresponds to the spike raster plot from the central electrode of every microchannel. **E.** Box-and-whiskers plots of the active microchannels’ MFR at baseline and 0h, 3h and 24 post-treatment. Data from 35-61 microchannels from 3-5 independent μEFs. **F.** Before-after plot of every active microchannel after treatment. ns = not significant, *0.01 < p < 0.05, ** 0.001 < p < 0.01, *** p < 0.001, **** p < 0.0001.

Unlike central nervous system neurons in culture (e.g. hippocampal neurons), DRG did not fire in bursts but rather exhibited sporadic spontaneous spiking at baseline. Under normal conditions, this relatively low level of spontaneous activity also occurs *in vivo* (Weng et al., 2012). Regardless of the explant position (outside or inside the array area), in most cases, we could only detect activity within the microchannels (Figure 7D). Still, this allowed us to directly compare baseline and post-treatment levels of axonal activity, as most axons within the microchannels are expected to have extended to the axonal compartment.

DRG terminals were exposed to three conditions comprising osteoclast secretome: i) mature, non-resorbing. osteoclasts (CM), ii) resorbing osteoclasts on mineralized substrates (bone slices – BS) and iii) EV-depleted osteoclasts secretome and compared to NGF-containing neurobasal control (NB).

Electrophysiology recordings show that the DRG exposed to osteoclasts secretome present an increased mean firing rate (MFR) when compared to the baseline recording immediately before treatment (Figure 7D-F). While the control treatment, supplemented with NGF, did not produce a significant effect in the MFR, the DRG treated with secretome from resorbing osteoblasts (BS) presented a highly significative effect. Previous reports have shown that medium acidification and the molecules released by osteoclasts and matrix itself, upon matrix degradation, contribute to the activation of nociceptors on the nerve terminals (Hiasa et al., 2017; Yoneda et al., 2015a). Here we show that the secretome from mature, non-resorbing, osteoclasts, also contribute to the electrical activation and signal propagation across sensory neurons. Excitingly, the effect on the firing rate of the sensory neurons was dramatically reduced upon EV depletion from the osteoclast secretome (Figure 7D-F). The effect produced by EV-depleted secretome, when compared to the pre-treatment baseline, was 598% lower than the parallel effect produced by osteoclasts CM and 1990% lower than the BS secretome (Figure 7F). Therefore, we postulate that the sensory neurons electrophysiological activity is dependent on the osteoclasts secreted EV, still, the mechanisms that support this effect deserve further elucidation.

## Discussion

We provide evidence on a new mechanism by which peripheral innervation can be coordinated. We show that sensory nerves outgrowth and electric signal propagation are dependent on the EV secreted by osteoclasts, the bone resorbing cells. Furthermore, we demonstrate that the axonal sprouting is achieved through the activation of epidermal-growth factor receptor (EGFR) and ErbB2 receptors, highlighting their involvement in the crosstalk, independent of β1 integrin function. Furthermore, inhibition of EGFR/EGFR and EGFR/ErbB2 dimers uncovers the contribution of this signaling cascade to the osteoclast-mediated axonal outgrowth. We proved that the EV-depleted osteoclast secretome leads to a significant decrease of neurons firing rate and axonal sprouting, concomitant with a decrease of EGFR/ErbB2 activation levels. Excitingly, the proteomic analysis of the osteoclast-derived EV cargo shows a high correlation with synaptic components reinforcing the role on sensory neurons/osteoclast crosstalk.

Different studies point to a parallel between osteoclast activity and sensory neurons outgrowth in physiological and pathological conditions (Cheung et al., 2009; Clarke, 2008; Gajda et al., 2005; Kumar et al., 2011; Mediati et al., 2014; Nagae et al., 2006; Ni et al., 2019; Reddy et al., 1999; Riminucci et al., 2003; Sisask et al., 1996, 2013; Zhu et al., 2019). Several reports have described that during embryonic long bone development, osteoclasts are found in the process of endochondral ossification, degrading the newly formed trabecular bone, creating the bone marrow space (Clarke, 2008). In another hand, longitudinal studies on the innervation process showed nerves appearing in areas with high osteogenic and chondrogenic activity during embryonic development (Gajda et al., 2005; Sisask et al., 2013, 1996). Thus, the presence of osteoclasts in stages where innervation occurs suggests a role of these cells in the innervation pattern. Bone pathologies with increased osteoclast number (Paget’s disease (Kumar et al., 2011; Reddy et al., 1999), fibrous dysplasia (Riminucci et al., 2003), *osteogenesis imperfecta* (Cheung et al., 2009)), and with increased osteoclast activity (osteoporosis and metastatic bone disease), are frequently associated with activation of sensory nerves (Mantyh, 2013; Mediati et al., 2014; Nagae et al., 2006). Although the pattern of innervation in these pathological conditions has only been described in metastatic bone disease, where exuberant sprouting of sensory nerve fibers has been reported, several studies indicate osteoclasts play a fundamental role in bone pain (Julius and Basbaum, 2001; Mantyh, 2014; Nagae et al., 2006; Yoneda et al., 2015b). Recently, Zhu *et al*. have shown that osteoclasts can actively promote axonal growth on subchondral bone, through netrin-1 signaling, in an osteoarthritis model (Zhu et al., 2019). Moreover, osteoclasts-derived netrin-1 was also linked to an increase in the sensory innervation of porous endplates of intervertebral disc (Ni et al., 2019). We were unable to detect NGF, BDNF, NT-3, NT4/5 nor netrin-1 in the osteoclast secretome by using comparable settings for osteoclast *in vitro* differentiation and netrin-1 quantification (mass spectrometry analysis and ELISA (data not shown)). In light of our results, we hypothesize that other signaling mechanisms were taking place. Herein we aimed to understand the main activation mechanisms on DRG explant cultures. Our work highlights new key players in this dynamic crosstalk. We detected high levels of EGFR and ErbB2 activation on DRG upon osteoclast secretome stimuli.

EGFR activation is involved in neuron differentiation and survival and has been implicated in responses to neuronal damage (Mizobuchi and Kanzaki, 2013; Nilsson and Kanje, 2005; Tsai et al., 2010; Xian and Zhou, 1999). Nevertheless, there is an extensive discussion in the literature concerning the direct involvement of EGFR activation/inhibition in axonal regeneration (Berry et al., 2011; Douglas et al., 2009; Liu et al., 2006; Wong and Guillaud, 2004). EGFR phosphorylation has been implicated in signaling inhibition of axonal growth in central nervous system (Berry et al., 2011; Koprivica et al., 2005; Liu et al., 2006). Koprivica *et al.* showed that EGFR inhibitors effectively promoted neurite outgrowth from cultured DRG (Koprivica et al., 2005). In contrast, others reported that EGFR knockdown did not induce neurite outgrowth of retinal ganglion cells, and inhibitors of EGFR were shown to enhance neurite outgrowth in the absence of the receptor (Berry et al., 2011). We and others previously demonstrated an increased expression of the EGFR family in DRG after lesion (Mizobuchi and Kanzaki, 2013; Neto et al., 2017; Xian and Zhou, 1999). Our data indicate that EGFR signaling has a role in axonal outgrowth promoted by osteoclast secretome. Upon EGFR inhibition with Erlotinib, described to inhibit both EGFR and ErbB2 receptor kinases at high concentrations (Macdonald-Obermann et al., 2013; Wood et al., 2004), we observed a significant reduction in the axonal outgrowth area. These findings are consistent with prior observations that the phosphorylation of EGFR enhances neurite outgrowth (Evangelopoulos et al., 2009; Goldshmit et al., 2004a, 2004b; Nilsson and Kanje, 2005; Tsai et al., 2010; Xu et al., 2014). The activation of EGFR in an integrin-dependent and EGFR ligand-independent manner has been described (Bill et al., 2004), where β1 integrin activation can enhance direct phosphorylation of EGFR (Balanis et al., 2011; Bill et al., 2004; Moro et al., 2002; Morozevich et al., 2012). We detected a significant decrease in axonal sprouting upon β1 integrin blocking, yet the EGFR family phosphorylation was not impaired, suggesting that activation of EGFR signaling cascade is not dependent on the cell-matrix or cell-cell mediated interactions.

Neurons have been described to be highly responsive to EV and exosomes derived from myelinating cells (oligodendrocytes and Schwann cells) (Budnik et al., 2016; Frühbeis et al., 2013; Lopez-Verrilli et al., 2013). Microglial-derived EV were also reported to promote synaptic refinement and instructing neurons upon inflammatory stimuli (Paolicelli et al., 2019). More recently, neurons were shown to be reactive to EV derived from other cellular populations, as seen for the response of cortical neurons to mesenchymal stem cells-derived EV (Xin et al., 2012; Zhang et al., 2017) and neuroblastoma cell line to EV from adipocyte derived Schwann cell-like (Ching et al., 2018). Osteoclasts secrete EV at the bone microenvironment in both physiological and pathological conditions (Yuan et al., 2018), which have been associated as key players underlying the osteoclast-osteoblast communication (Collison, 2017; Huynh et al., 2016a), tumor proliferation, angiogenesis, and tumor cell survival (Huynh et al., 2016b; Lee et al., 2007; Rossi et al., 2018).

Here, we firstly described that sensory neurons are responsive to osteoclast-derived EV. Our results strengthen the importance on the communication between sensory terminals and osteoclasts and unravels new mechanism underlying the interplay. We indicate that the EV signaling mechanism underlie sensory processing and nociception signal propagation, largely unknow in the literature (Fowler, 2019). Remarkably, EV-depleted osteoclast secretome produced not only a significant decrease in axonal growth, but also a striking reduction in EGFR family phosphorylation. Once again, the proteome of EV cargo revealed no presence of EGF or TGFα, which is in accordance with recent data reporting the proteomic profile of the osteoclasts secreted vesicles upon different substrates (Rody et al., 2019). Importantly, the proteomic screening of the EV cargo, revealed the presence of SH3BGRL, proved to bind to the intracellular EGFR domain (Chiang et al., 2015; Schulze et al., 2005; Taguchi et al., 2011). More interestingly, phosphotyrosine interactome of the ErbB-receptor kinase family showed that ErbB2 has few interaction partners being SH3BGRL one of them on residue Y0923 (Schulze et al., 2005). Noteworthy, SH3BGRL promote the dowstream activation of c-Src (Wang et al., 2016), which is central to multiple signal transduction pathways including axonal elongation, adhesion, growth, and cell survival (Irby and Yeatman, 2000; Robles et al., 2005). The main mechanism underlying the EV interaction with sensory neurons require further studies, still it has been reported that mesenchymal stem cells-derived EV are capable of entering cortical neurons both through cell body and axonal terminals (Zhang et al., 2017).

We also show that different electrophysiological patterns are revealed depending on the osteoclast secretome tested. The greater effect observed in the increasing of the MFR was related to the secretome from osteoclasts cultured on mineralized substrates, as expected. Osteoclasts were shown to increase the expression of sodium channel Na_v_ 1.8 in sensory nerve endings in the endplates leading to spinal hypersensitivity (Ni et al., 2019). Protons released by the osteoclasts upon matrix degradation are known to stimulate the acid-sensing ions channels contributing to the activation of nociceptors on the nerve terminals (Hiasa et al., 2017; Yoneda et al., 2015a). We demonstrate that, not only the products released through matrix degradation process are mediating the sensory fibers activation, but also the products secreted by osteoclasts in non-degrading scenario. The EV-depletion resulted in a major decrease on the MFR reflecting a possible pathway to be addressed to understand the nerves sensitization on bone microenvironment.

In conclusion, we provide a new mechanism on the sensory nerves sprouting, indicating that the effect is dependent on the extracellular vesicles (EV) released by osteoclasts, through the epidermal growth factor receptor family targeting, by integrin independent pathways. We show different electrophysiology patterns being triggered in the presence of osteoclasts secretome and the abolishment of sensory neurons firing rate in EV-depleted conditions. Clearly, besides the activation of nociceptors through microenvironment acidification or soluble molecules, osteoclasts secreted vesicles can be implicated in cancer-associated bone pain through the control of nerve sprouting and electrical signal propagation, consequently remodeling the pathological innervation pattern and activity. Furthermore, studies aiming to manipulate osteoclast-derived EV cargo can be employed to further dissect the bone-peripheral nervous system communication. This is a subject that deserves further investigation to i) control/reduce the formation of neuroma-like structures and ii) specify new candidates as potential targets for bone pain therapies.

## Author contributions

Conceptualization, Estrela Neto, Meriem Lamghari and Paulo Aguiar; Methodology, Estrela Neto, Luís Leitão, José Mateus, Daniela Monteiro Sousa and Francisco Conceição; Investigation, Estrela Neto, Luís Leitão, José Mateus, Daniela Monteiro Sousa and Cecília Juliana Alves; Writing – Original Draft, Estrela Neto, José Mateus and Meriem Lamghari; Writing – Review & Editing, all authors; Funding Acquisition, Meriem Lamghari and Paulo Aguiar; Supervision, Richard Oreffo, Jonathan West and Meriem Lamghari.

## Acknowledgments

The authors would like to acknowledge to João H. Morais-Cabral from i3S for the fruitful scientific discussions and input for this work. To Susana Santos, Andreia Silva and José Teixeira for the help with EV isolation protocol. To Maria José Oliveira, José Carlos Machado and Ana Paula Pêgo groups from i3S for the phospho-EGFR antibody, Erlotinib and anti-β1 integrin antibodies, respectively. The microfluidic design adapted to the commercial microelectrode arrays used in this study were fabricated at INESC - Microsystems and Nanotechnologies, Portugal, under the supervision of João Pedro Conde and Virginia Chu.

This work was financed by FEDER - Fundo Europeu de Desenvolvimento Regional funds through the COMPETE 2020 - Operacional Programme for Competitiveness and Internationalization (POCI), Portugal 2020, and by Portuguese funds through FCT/MCTES in the framework of the project SproutOC: POCI-01-0145-FEDER-030158, PTDC/MED-PAT/30158/2017. LL, JM and FC are recipients of Ph.D. fellowships (SFRH/BD/109686/2015, PD/BD/135491/2018 and SFRH/BD/128771/2017, respectively). DMS is recipient of Post-Doc fellowship (SFRH/BPD/115341/2016). The authors acknowledge the support of the i3S Scientific Platforms BioSciences Screening, Bioimaging, and Histology and Electron Microscopy, members of the national infrastructure PPBI - Portuguese Platform of Bioimaging. The mass spectrometry technique was performed at the Proteomics i3S Scientific Platform with the assistance of Hugo Osório.

The authors declare no competing interests.

