## Supplemental data for "Sensory neurons sprouting is dependent on osteoclast-derived extracellular vesicles involving the activation of epidermal growth factor receptors"

### Supplemental information

#### S1. Evaluation of osteoclast differentiation

Evaluation of osteoclast differentiation was performed by qRT-PCR analysis. RNA was isolated and purified as previously described in (Neto *et al.*, 2017) and the mRNA expression of differentiation markers (tartrate-resistant acid phosphatase (TRAP, Fw: CGACCATTGTTAGCCACATACG; Rv: TCGTCCTGAAGATACTGCAGGTT, Invitrogen, Carlsbad, CA, USA), cathepsin K (CTSK, Fw: ATATGTGGGCCAGGATGAAAGTT; Rv: TCGTTCCCCACAGGAATCTCT, Invitrogen), osteoclast associated immunoglobulin-like receptor (OscAR, Fw: TGGCGGTTTGCACTCTTCA; Rv: GATCCGTTACCAGCAGTTCCAGA, Invitrogen) and glyceraldehyde 3-phosphate dehydrogenase (GAPDH, Fw: GCCTTCCGTGTTCCTACC; Rv: AGAGTGGGAGTTGCTGTTG, Invitrogen) was performed.

Immunocytochemistry was performed to assess the morphology of the differentiated osteoclasts and the possible presence of macrophages in culture. Briefly, cells were fixed with 4% paraformaldehyde (PFA, Merck Millipore, Kenilworth, NJ, USA) in PBS during 10 min followed by permeabilization with 0.25% (v/v) Triton X-100 (Sigma-Aldrich, St. Louis, MO, USA) in PBS and incubated, for 30 min at RT, with blocking solution composed of 1% (w/v) Bovine Serum Albumin (BSA, Sigma-Aldrich) in PBS for 30 min at RT to block antibody unspecific binding. The cells were incubated with macrophages specific marker – anti-F4/80 (#14-4801-85, eBioscience, Thermo Fisher Scientific), 1:250 in blocking solution overnight at 4 °C. Afterwards cells were washed with PBS and incubated with Alexafluor 488-Phalloidin (Life Technologies, Thermo Fisher Scientific) 1:100 in PBS for 30 minutes at RT in the dark. At the end of the incubation time, the cells were washed with PBS and mounted using Vectashield with DAPI (Vector Labs, Burlingame, California, USA). Images were captured using the Inverted Fluorescence Microscope (Zeiss AxioVert, Carl Zeiss, Oberkochen, Germany), equipped with AxioVision SE64 Rel. 4.8 software. TRAP staining was also performed using an Acid phosphatase, Leukocyte (TRAP) kit (Sigma-Aldrich) according to manufacturer's instructions. Images were acquired using a stereomicroscope (SZX10, Olympus, Shinjuku, Tokyo, Japan) coupled to a digital camera (DP21, Olympus).

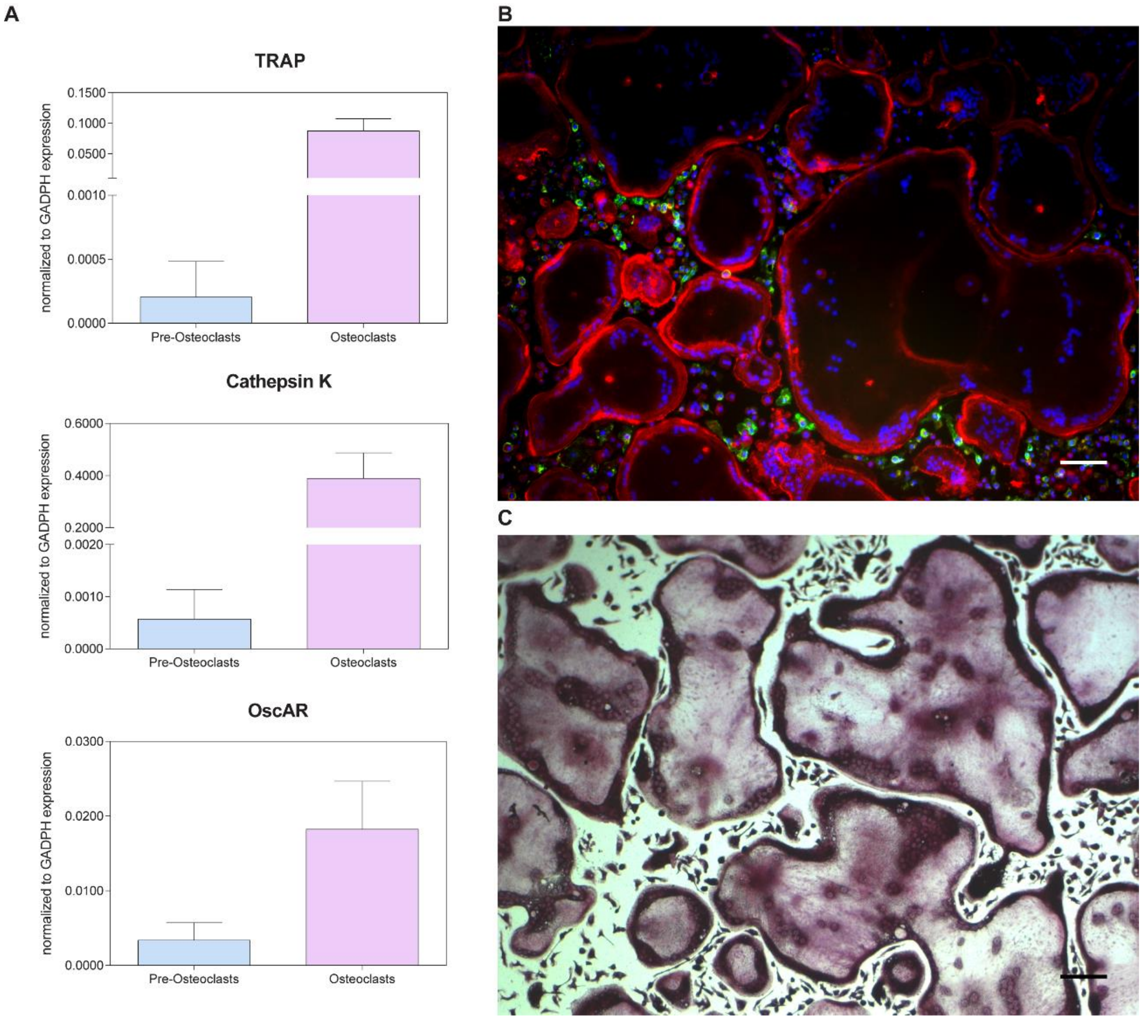


**Figure S1: Characterization of osteoclasts differentiation *in vitro*.** Osteoclasts were differentiated from mouse bone marrow hematopoietic lineage. **A.** Differentiation markers (tartrate-resistant acid phosphatase (TRAP), cathepsin K (CTSK) and osteoclast associated immunoglobulin-like receptor (OscAR)) were evaluated by qPCR. Data represented as mean±S.D. **B.** Image of multinucleated osteoclasts stained for F-actin (red), nuclei (blue) and macrophages stained with F4/80 (green), revealed a large population of multinucleated osteoclasts with the low contribution of positive F4/80 macrophages present in the culture. **C.** TRAP staining of mature osteoclasts differentiated *in vitro*. Scale bar 100 µm.

#### S2. Drug toxicity assay – EGFR inhibitor

Cell viability assay was performed for the DRG treated with the highest concentration of Erlotinib to rule out the possible drug toxicity. Briefly, DRG were incubated with Calcein AM (Invitrogen) in PBS for 30 min at 37˚C. Calcein AM was washed out and DRG were incubated with propidium iodide (Sigma Aldrich) for 10 min at 37˚C. Images were acquired using IN Cell Analyzer 2000 equipped with IN Cell Investigator software. To address the DRG metabolic activity, resazurin solution (Sigma) was added to wells at a final concentration of 10% (v/v). After 4 h at 37 °C, 100 µL (pooled from two wells) were transferred to a 96-well black plate and fluorescence was measured (530 nm/590 nm) in a Spectra Max Gemini XS (Molecular Devices).

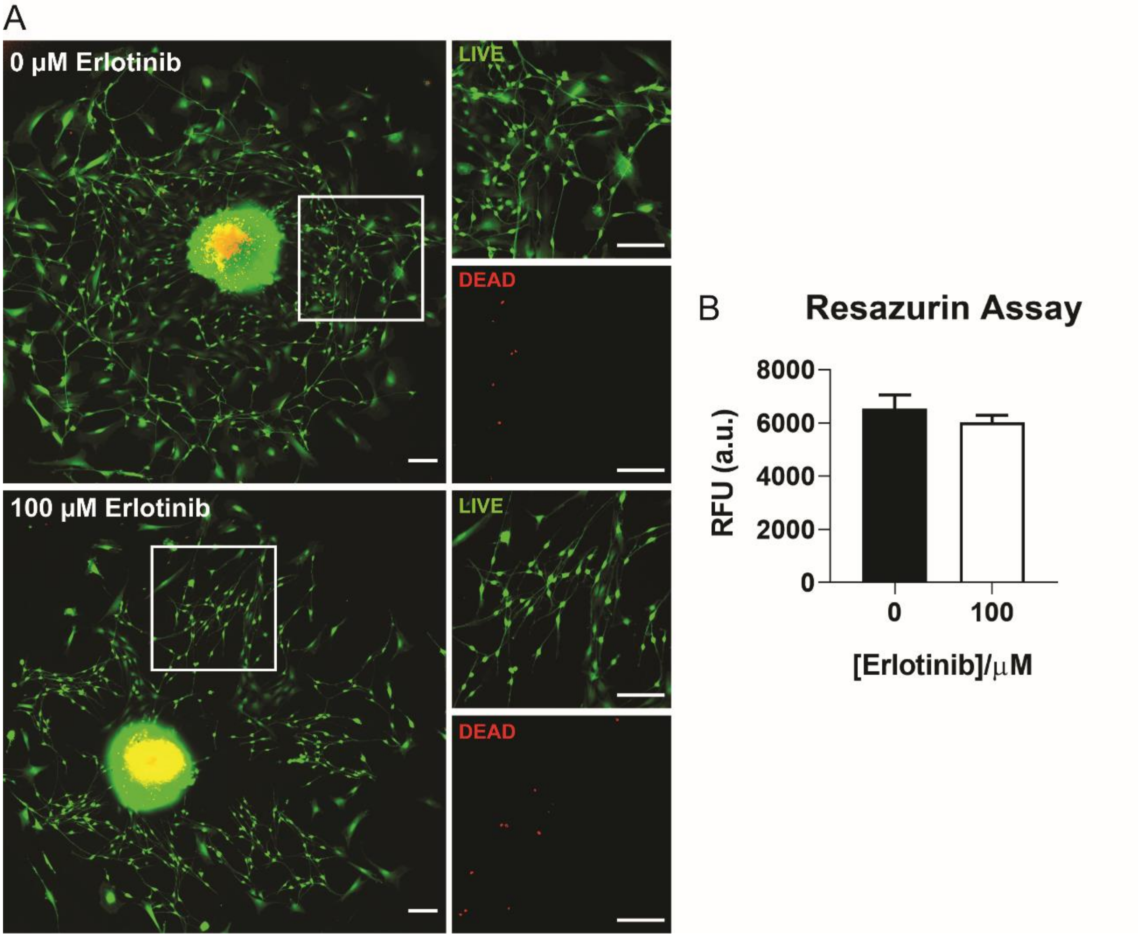


**Figure S2: Cell viability and metabolic activity of DRG upon treatment with 100 µM of Erlotinib. A.** Live/dead assay showing the live cells in green and dead cells in red for control conditions (neurobasal+NGF) and 100 µM of Erlotinib. Scale bar 100 µm. **B.** Evaluation of the metabolic activity of the DRG in control conditions (neurobasal+NGF) and 100 µM of Erlotinib. Data presented as relative fluorescence units (RFU).

#### S3. Evaluation of axonal outgrowth in the presence of β1 integrin blocker and its isotype

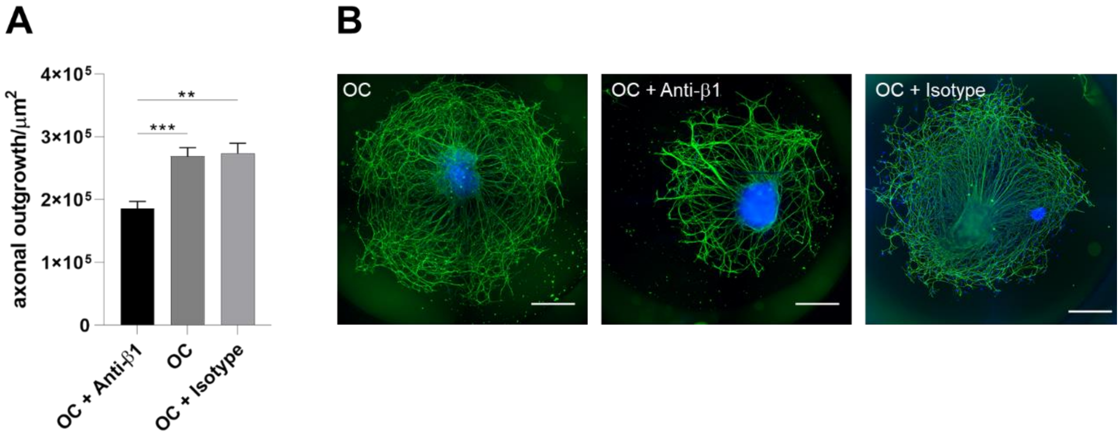


**Figure S3**: **Evaluation of axonal outgrowth in the presence of β1 integrin blocker and its isotype. A.** Quantification of axonal sprouting of DRG stimulated for 72h with osteoclast secretome with the functional anti-β1 integrin, osteoclast secretome and osteoclast secretome with the isotype of anti-β1 integrin. $^{**}p \leq 0.01$; $^{***}p \leq 0.001$. **B.** Representative images of DRG treated for 72h with osteoclast secretome, osteoclast secretome with the functional anti-β1 integrin and osteoclast secretome with the isotype of anti-β1 integrin. Staining for βIII tubulin in green and nuclei in blue, scale bar 500 µm.

**S4. Nanoparticle tracking analysis (NTA) of the osteoclast-derived EV enriched fraction**

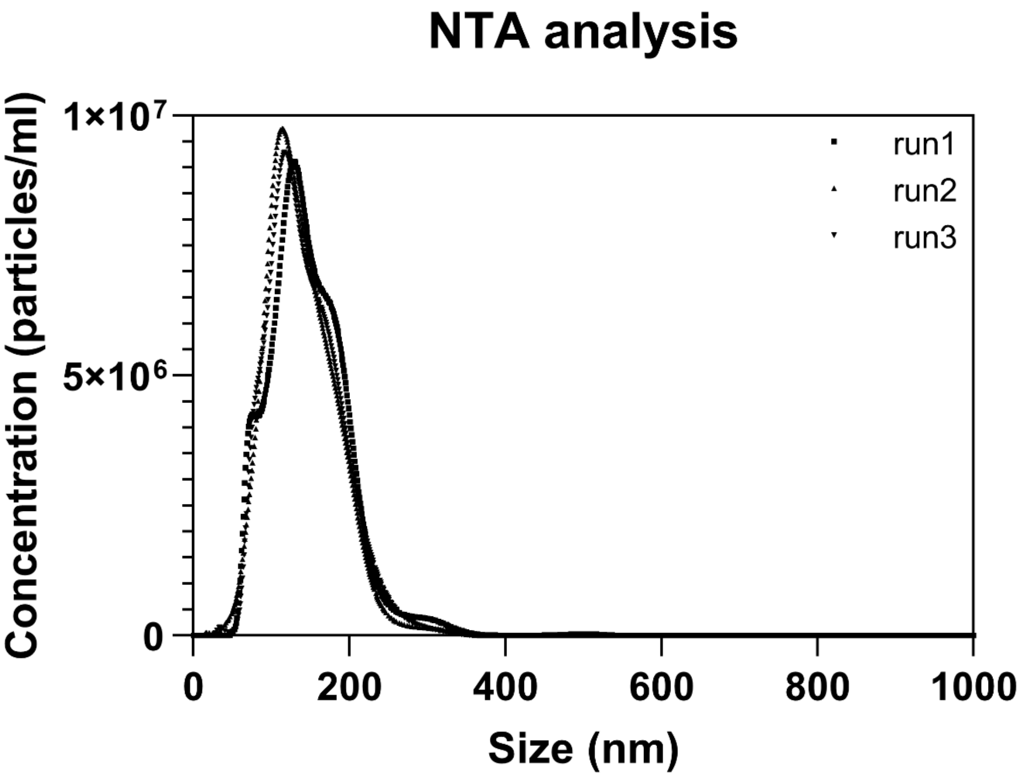


**Figure S4**: Characterization of concentration *versus* size distribution of osteoclast-derived EV enriched fraction (diluted in filtered PBS 1:500), by nanoparticle tracking analysis (NTA). Plot is representative of EV size and concentration analysis with NanoSight NS300.
