## Supplemental table S1 for "Sensory neurons sprouting is dependent on osteoclast-derived extracellular vesicles involving the activation of epidermal growth factor receptors"

Table 1: LC-MS/MS analysis of protein content from serum-free osteoclasts secretome at 6h and 24h.

| ACCESSION | DESCRIPTION | SUM PEP SCORE | # UNIQUE PEPTIDES | MW [KDA] | ABUNDANCE RATIO: (24H) / (6H) |
|---|---|---|---|---|---|
| **Q3THB4** | L-lactate dehydrogenase OS=Mus musculus OX=10090 GN=Ldha PE=2 SV=1 | 106,046 | 2 | 36,5 | 100 |
| **G3X8T3** | Carboxypeptidase OS=Mus musculus OX=10090 GN=Ctsa PE=1 SV=1 | 37,76 | 5 | 55,7 | 100 |
| **P68372** | Tubulin beta-4B chain OS=Mus musculus OX=10090 GN=Tubb4b PE=1 SV=1 | 22,581 | 1 | 49,8 | 100 |
| **Q3TEU8** | Coronin OS=Mus musculus OX=10090 GN=Coro1c PE=2 SV=1 | 21,46 | 3 | 53,1 | 100 |
| **P80316** | T-complex protein 1 subunit epsilon OS=Mus musculus OX=10090 GN=Cct5 PE=1 SV=1 | 18,961 | 3 | 59,6 | 100 |
| **P08730** | Keratin, type I cytoskeletal 13 OS=Mus musculus OX=10090 GN=Krt13 PE=1 SV=2 | 16,625 | 1 | 47,7 | 100 |
| **Q3TAW4** | Protein transport protein SEC23 OS=Mus musculus OX=10090 GN=Sec23b PE=2 SV=1 | 16,539 | 4 | 86,3 | 100 |
| **P50518** | V-type proton ATPase subunit E 1 OS=Mus musculus OX=10090 GN=Atp6v1e1 PE=1 SV=2 | 15,042 | 2 | 26,1 | 100 |
| **Q69Z71** | MKIAA1905 protein (Fragment) OS=Mus musculus OX=10090 GN=Scin PE=2 SV=1 | 14,823 | 6 | 83 | 100 |
| **P08226** | Apolipoprotein E OS=Mus musculus OX=10090 GN=Apoe PE=1 SV=2 | 13,845 | 2 | 35,8 | 100 |
| **P99026** | Proteasome subunit beta type-4 OS=Mus musculus OX=10090 GN=Psmb4 PE=1 SV=1 | 13,665 | 2 | 29,1 | 100 |
| **P97371** | Proteasome activator complex subunit 1 OS=Mus musculus OX=10090 GN=Psme1 PE=1 SV=2 | 13,461 | 3 | 28,7 | 100 |
| **O08992** | Syntenin-1 OS=Mus musculus OX=10090 GN=Sdcbp PE=1 SV=1 | 12,747 | 3 | 32,4 | 100 |
| **Q8C338** | Isocitrate dehydrogenase [NADP] OS=Mus musculus OX=10090 GN=Idh1 PE=2 SV=1 | 12,598 | 3 | 47,5 | 100 |
| **Q8VDC3** | Cytoplasmic aconitase OS=Mus musculus OX=10090 GN=aco1 PE=3 SV=1 | 12,473 | 3 | 99 | 100 |
| **Q80X90** | Filamin-B OS=Mus musculus OX=10090 GN=Flnb PE=1 SV=3 | 12,205 | 3 | 277,7 | 100 |
| **Q544Z7** | DNA-(apurinic or apyrimidinic site) lyase OS=Mus musculus OX=10090 GN=Apex1 PE=1 SV=1 | 12,151 | 2 | 35,5 | 100 |

| | | | | | |
|---|---|---|---|---|---|
| **P46664** | Adenylosuccinate synthetase isozyme 2 OS=Mus musculus OX=10090 GN=Adss PE=1 SV=2 | 12,045 | 4 | 50 | 100 |
| **Q9QUM9** | Proteasome subunit alpha type-6 OS=Mus musculus OX=10090 GN=Psma6 PE=1 SV=1 | 11,346 | 3 | 27,4 | 100 |
| **O55234** | Proteasome subunit beta type-5 OS=Mus musculus OX=10090 GN=Psmb5 PE=1 SV=3 | 10,957 | 1 | 28,5 | 100 |
| **Q3U536** | Uncharacterized protein OS=Mus musculus OX=10090 GN=Npm1 PE=2 SV=1 | 10,437 | 2 | 32,5 | 100 |
| **A0A0G2JEA5** | Predicted gene 43738 OS=Mus musculus OX=10090 GN=Gm43738 PE=4 SV=1 | 9,964 | 2 | 95,2 | 100 |
| **Q6P8R3** | C-X-C motif chemokine OS=Mus musculus OX=10090 GN=Pf4 PE=2 SV=1 | 9,633 | 1 | 11,2 | 100 |
| **Q3U6A3** | Monocyte differentiation antigen CD14 OS=Mus musculus OX=10090 GN=Cd14 PE=2 SV=1 | 9,403 | 3 | 39,1 | 100 |
| **A0A1G5SJA5** | OTTMUSWSBP00075795 OS=Mus musculus domesticus OX=10092 GN=OTTMUSWSBG00059279 PE=4 SV=1 | 9,13 | 1 | 27 | 100 |
| **Q8BG13** | RNA-binding protein 3 OS=Mus musculus OX=10090 GN=Rbm3 PE=1 SV=1 | 9,129 | 2 | 16,8 | 100 |
| **Q3UP42** | S100 calcium binding protein A9 (Calgranulin B), isoform CRA_a OS=Mus musculus OX=10090 GN=S100a9 PE=1 SV=1 | 8,99 | 1 | 13 | 100 |
| **Q7TMR0** | Lysosomal Pro-X carboxypeptidase OS=Mus musculus OX=10090 GN=Prcp PE=1 SV=2 | 8,947 | 2 | 55 | 100 |
| **Q3UK83** | Uncharacterized protein OS=Mus musculus OX=10090 GN=Hnrnpa1 PE=2 SV=1 | 8,94 | 2 | 38,8 | 100 |
| **B2CQD6** | Fibulin-1 OS=Mus musculus OX=10090 GN=Fbln1 PE=2 SV=1 | 8,759 | 3 | 78 | 100 |
| **Q8BP47** | Asparagine--tRNA ligase, cytoplasmic OS=Mus musculus OX=10090 GN=Nars PE=1 SV=2 | 8,673 | 5 | 64,2 | 100 |
| **A0A0G2JGL0** | Ubiquitin-conjugating enzyme E2 D3 OS=Mus musculus OX=10090 GN=Ube2d3 PE=1 SV=1 | 8,64 | 1 | 16,8 | 100 |
| **Q3TL79** | Uncharacterized protein OS=Mus musculus OX=10090 GN=Ahsa1 PE=2 SV=1 | 8,319 | 1 | 38,1 | 100 |
| **Q61081** | Hsp90 co-chaperone Cdc37 OS=Mus musculus OX=10090 GN=Cdc37 PE=1 SV=1 | 8,253 | 2 | 44,6 | 100 |
| **Q3U5N4** | ADP-ribosylarginine hydrolase, isoform CRA_a OS=Mus musculus OX=10090 GN=Adprh PE=1 SV=1 | 8,109 | 1 | 40 | 100 |
| **A2AE89** | Glutathione S-transferase Mu 1 OS=Mus musculus OX=10090 GN=Gstm1 PE=1 SV=2 | 7,829 | 1 | 28,5 | 100 |
| **I6L9I7** | Cbx3 protein OS=Mus musculus OX=10090 GN=Cbx3 PE=2 SV=1 | 7,365 | 1 | 20,7 | 100 |

| | | | | | |
|---|---|---|---|---|---|
| **Q3UA53** | Uncharacterized protein OS=Mus musculus OX=10090 GN=Ppa1 PE=2 SV=1 | 7,256 | 1 | 32,6 | 100 |
| **O70435** | Proteasome subunit alpha type-3 OS=Mus musculus OX=10090 GN=Psma3 PE=1 SV=3 | 7,254 | 2 | 28,4 | 100 |
| **P08030** | Adenine phosphoribosyltransferase OS=Mus musculus OX=10090 GN=Aprt PE=1 SV=2 | 7,112 | 1 | 19,7 | 100 |
| **Q3TYK4** | Uncharacterized protein OS=Mus musculus OX=10090 GN=Prkar1a PE=2 SV=1 | 7,079 | 2 | 57,2 | 100 |
| **E9PYD5** | Transcription elongation factor A protein 1 OS=Mus musculus OX=10090 GN=Tcea1 PE=1 SV=1 | 6,901 | 1 | 35 | 100 |
| **Q8CIJ3** | Eukaryotic translation initiation factor 3 subunit B OS=Mus musculus OX=10090 GN=Eif3b PE=2 SV=1 | 6,674 | 2 | 108,9 | 100 |
| **P42208** | Septin-2 OS=Mus musculus OX=10090 GN=Sept2 PE=1 SV=2 | 6,621 | 2 | 41,5 | 100 |
| **Q99KP6-2** | Isoform 2 of Pre-mRNA-processing factor 19 OS=Mus musculus OX=10090 GN=Prpf19 | 6,59 | 1 | 57,3 | 100 |
| **B2CY77** | Laminin receptor (Fragment) OS=Mus musculus OX=10090 GN=Rpsa PE=2 SV=1 | 6,579 | 1 | 32,8 | 100 |
| **B7ZP20** | Ankyrin repeat and FYVE domain containing 1 OS=Mus musculus OX=10090 GN=Ankfy1 PE=2 SV=1 | 6,405 | 2 | 128,6 | 100 |
| **Q8BJY1** | 26S proteasome non-ATPase regulatory subunit 5 OS=Mus musculus OX=10090 GN=Psmd5 PE=1 SV=4 | 6,167 | 2 | 55,9 | 100 |
| **P97351** | 40S ribosomal protein S3a OS=Mus musculus OX=10090 GN=Rps3a PE=1 SV=3 | 6,163 | 2 | 29,9 | 100 |
| **O70251** | Elongation factor 1-beta OS=Mus musculus OX=10090 GN=Eef1b PE=1 SV=5 | 6,047 | 1 | 24,7 | 100 |
| **Q8C483** | Serine--tRNA ligase, cytoplasmic OS=Mus musculus OX=10090 GN=Sars PE=1 SV=1 | 6,03 | 2 | 61,1 | 100 |
| **Q9R059** | Four and a half LIM domains protein 3 OS=Mus musculus OX=10090 GN=Fhl3 PE=1 SV=2 | 5,951 | 1 | 31,8 | 100 |
| **P98086** | Complement C1q subcomponent subunit A OS=Mus musculus OX=10090 GN=C1qa PE=1 SV=2 | 5,814 | 2 | 26 | 100 |
| **Q7TSV4** | Phosphoglucomutase-2 OS=Mus musculus OX=10090 GN=Pgm2 PE=1 SV=1 | 5,7 | 3 | 68,7 | 100 |
| **P62315** | Small nuclear ribonucleoprotein Sm D1 OS=Mus musculus OX=10090 GN=Snrpd1 PE=1 SV=1 | 5,645 | 1 | 13,3 | 100 |
| **Q3TI61** | 26S proteasome non-ATPase regulatory subunit 2 OS=Mus musculus OX=10090 GN=Psmd2 PE=2 SV=1 | 5,632 | 2 | 100,2 | 100 |

| | | | | | |
|---|---|---|---|---|---|
| **F8WJ05** | Inter-alpha-trypsin inhibitor heavy chain H1 OS=Mus musculus OX=10090 GN=Itih1 PE=1 SV=1 | 5,554 | 1 | 101,6 | 100 |
| **C5H0E8** | Rap1A-retro1 OS=Mus musculus OX=10090 GN=Gm9392 PE=2 SV=1 | 5,516 | 1 | 21 | 100 |
| **E9QK48** | Echinoderm microtubule-associated protein-like 2 OS=Mus musculus OX=10090 GN=Eml2 PE=1 SV=1 | 5,412 | 1 | 90,7 | 100 |
| **J3QN31** | Adenylosuccinate synthetase isozyme 1 OS=Mus musculus OX=10090 GN=Adssl1 PE=1 SV=1 | 5,374 | 1 | 52,7 | 100 |
| **P62317** | Small nuclear ribonucleoprotein Sm D2 OS=Mus musculus OX=10090 GN=Snrpd2 PE=1 SV=1 | 5,313 | 1 | 13,5 | 100 |
| **P62746** | Rho-related GTP-binding protein RhoB OS=Mus musculus OX=10090 GN=Rhob PE=1 SV=1 | 5,302 | 1 | 22,1 | 100 |
| **Q9DBG5** | Perilipin-3 OS=Mus musculus OX=10090 GN=Plin3 PE=1 SV=1 | 5,281 | 1 | 47,2 | 100 |
| **Q8BT90** | Uncharacterized protein (Fragment) OS=Mus musculus OX=10090 GN=Rps17 PE=2 SV=1 | 5,242 | 1 | 16,1 | 100 |
| **F8WIP8** | Osteopontin OS=Mus musculus OX=10090 GN=Spp1 PE=1 SV=1 | 5,135 | 1 | 32,6 | 100 |
| **Q571F9** | MKIAA4115 protein (Fragment) OS=Mus musculus OX=10090 GN=G3bp1 PE=2 SV=1 | 5,037 | 1 | 56,1 | 100 |
| **Q920A5** | Retinoid-inducible serine carboxypeptidase OS=Mus musculus OX=10090 GN=Scpep1 PE=1 SV=2 | 4,973 | 1 | 50,9 | 100 |
| **P34022** | Ran-specific GTPase-activating protein OS=Mus musculus OX=10090 GN=Ranbp1 PE=1 SV=2 | 4,963 | 2 | 23,6 | 100 |
| **O35474** | EGF-like repeat and discoidin I-like domain-containing protein 3 OS=Mus musculus OX=10090 GN=Edil3 PE=1 SV=2 | 4,955 | 1 | 53,7 | 100 |
| **Q9WTP6** | Adenylate kinase 2, mitochondrial OS=Mus musculus OX=10090 GN=Ak2 PE=1 SV=5 | 4,869 | 2 | 26,5 | 100 |
| **B2RXY7** | Carbonyl reductase 1 OS=Mus musculus OX=10090 GN=Cbr1 PE=1 SV=1 | 4,867 | 2 | 30,6 | 100 |
| **P35951** | Low-density lipoprotein receptor OS=Mus musculus OX=10090 GN=Ldlr PE=1 SV=2 | 4,826 | 1 | 94,9 | 100 |
| **Q9QYB1** | Chloride intracellular channel protein 4 OS=Mus musculus OX=10090 GN=Clic4 PE=1 SV=3 | 4,813 | 2 | 28,7 | 100 |
| **Q9JIF7** | Coatomer subunit beta OS=Mus musculus OX=10090 GN=Copb1 PE=1 SV=1 | 4,774 | 1 | 107 | 100 |

| | | | | | |
|---|---|---|---|---|---|
| **P11087** | Collagen alpha-1(I) chain OS=Mus musculus OX=10090 GN=Col1a1 PE=1 SV=4 | 4,663 | 1 | 137,9 | 100 |
| **A0A068BFR3** | RAS oncogene family protein OS=Mus musculus OX=10090 GN=Rab11b PE=2 SV=1 | 4,646 | 1 | 24,5 | 100 |
| **Q50HX0** | RAB14 protein OS=Mus musculus OX=10090 GN=Rab14 PE=2 SV=1 | 4,611 | 1 | 23,9 | 100 |
| **Q3TJF2** | Obg-like ATPase 1 OS=Mus musculus OX=10090 GN=Ola1 PE=2 SV=1 | 4,588 | 1 | 44,8 | 100 |
| **Q0VBA8** | Plasminogen activator, urokinase OS=Mus musculus OX=10090 GN=Plau PE=1 SV=1 | 4,448 | 1 | 48,2 | 100 |
| **Q9DCL9** | Multifunctional protein ADE2 OS=Mus musculus OX=10090 GN=Paics PE=1 SV=4 | 4,335 | 1 | 47 | 100 |
| **Q3TIN2** | Uncharacterized protein OS=Mus musculus OX=10090 GN=Qars PE=2 SV=1 | 4,302 | 1 | 87,7 | 100 |
| **Q3U1J4** | DNA damage-binding protein 1 OS=Mus musculus OX=10090 GN=Ddb1 PE=1 SV=2 | 4,147 | 1 | 126,8 | 100 |
| **Q3TG93** | Uncharacterized protein OS=Mus musculus OX=10090 GN=Ssb PE=2 SV=1 | 4,12 | 1 | 47,7 | 100 |
| **Q3TXS7** | 26S proteasome non-ATPase regulatory subunit 1 OS=Mus musculus OX=10090 GN=Psmd1 PE=1 SV=1 | 4,112 | 2 | 105,7 | 100 |
| **Q3UBK2** | Uncharacterized protein OS=Mus musculus OX=10090 PE=2 SV=1 | 3,994 | 1 | 24,9 | 100 |
| **F6X9I3** | Peptidylprolyl isomerase (Fragment) OS=Mus musculus OX=10090 GN=Fkbp1a PE=1 SV=1 | 3,886 | 1 | 18,4 | 100 |
| **A0A1B0GRP7** | Pyridoxal phosphate homeostasis protein (Fragment) OS=Mus musculus OX=10090 GN=Plpbp PE=1 SV=1 | 3,857 | 1 | 36,1 | 100 |
| **P62245** | 40S ribosomal protein S15a OS=Mus musculus OX=10090 GN=Rps15a PE=1 SV=2 | 3,748 | 1 | 14,8 | 100 |
| **P14106** | Complement C1q subcomponent subunit B OS=Mus musculus OX=10090 GN=C1qb PE=1 SV=2 | 3,744 | 1 | 26,7 | 100 |
| **D3Z1V4** | Phosphatidylethanolamine-binding protein 1 OS=Mus musculus OX=10090 GN=Pebp1 PE=1 SV=1 | 3,703 | 1 | 23 | 100 |
| **O08997** | Copper transport protein ATOX1 OS=Mus musculus OX=10090 GN=Atox1 PE=1 SV=1 | 3,678 | 2 | 7,3 | 100 |
| **Q69ZW4** | MKIAA0899 protein (Fragment) OS=Mus musculus OX=10090 GN=Ap2a2 PE=2 SV=1 | 3,638 | 2 | 107,3 | 100 |
| **P29391** | Ferritin light chain 1 OS=Mus musculus OX=10090 GN=Ftl1 PE=1 SV=2 | 3,624 | 1 | 20,8 | 100 |

| | | | | | |
|---|---|---|---|---|---|
| **E9PV22** | Leucine-rich repeat-containing protein 47 OS=Mus musculus OX=10090 GN=Lrrc47 PE=1 SV=1 | 3,59 | 2 | 65,9 | 100 |
| **Q3TXK6** | Uncharacterized protein OS=Mus musculus OX=10090 GN=Tsnax PE=2 SV=1 | 3,447 | 1 | 32,9 | 100 |
| **Q4VWZ5** | Acyl-CoA-binding protein OS=Mus musculus OX=10090 GN=Dbi PE=1 SV=1 | 3,435 | 2 | 15,2 | 100 |
| **Q8JZZ5** | Phosphatidylinositol transfer protein beta isoform OS=Mus musculus OX=10090 GN=Pitpnb PE=1 SV=1 | 3,421 | 1 | 31,6 | 100 |
| **Q3UER8** | Fibrinogen gamma chain OS=Mus musculus OX=10090 GN=Fgg PE=1 SV=1 | 3,418 | 1 | 50,3 | 100 |
| **Q3TDX7** | Extracellular matrix protein 1 OS=Mus musculus OX=10090 GN=Ecm1 PE=2 SV=1 | 3,392 | 1 | 62,8 | 100 |
| **A2AGH5** | Predicted gene 20489 OS=Mus musculus OX=10090 GN=Gm20489 PE=4 SV=1 | 3,392 | 1 | 51,9 | 100 |
| **O54782** | Epididymis-specific alpha-mannosidase OS=Mus musculus OX=10090 GN=Man2b2 PE=1 SV=2 | 3,304 | 1 | 115,5 | 100 |
| **Q3TVK3** | Aspartyl aminopeptidase OS=Mus musculus OX=10090 GN=Dnpep PE=1 SV=1 | 3,283 | 1 | 52,4 | 100 |
| **P57759** | Endoplasmic reticulum resident protein 29 OS=Mus musculus OX=10090 GN=Erp29 PE=1 SV=2 | 3,265 | 1 | 28,8 | 100 |
| **D3Z3G6** | Mitogen-activated protein kinase OS=Mus musculus OX=10090 GN=Mapk3 PE=1 SV=1 | 3,248 | 1 | 42,4 | 100 |
| **F6R7E8** | Predicted gene 2663 OS=Mus musculus OX=10090 GN=Gm2663 PE=3 SV=1 | 3,198 | 1 | 26,6 | 100 |
| **Q3UJ70** | 3-hydroxy-3-methylglutaryl coenzyme A synthase OS=Mus musculus OX=10090 GN=Hmgcs1 PE=2 SV=1 | 3,197 | 1 | 57,5 | 100 |
| **A0A1S6GWH1** | Uncharacterized protein OS=Mus musculus OX=10090 GN=Psmc5 PE=2 SV=1 | 3,196 | 2 | 46,6 | 100 |
| **Q8VIJ6** | Splicing factor, proline- and glutamine-rich OS=Mus musculus OX=10090 GN=Sfpq PE=1 SV=1 | 3,158 | 1 | 75,4 | 100 |
| **Q3UB60** | Uncharacterized protein OS=Mus musculus OX=10090 GN=Ppid PE=2 SV=1 | 3,152 | 2 | 40,7 | 100 |
| **Q9QZE5** | Coatomer subunit gamma-1 OS=Mus musculus OX=10090 GN=Copg1 PE=1 SV=1 | 3,112 | 1 | 97,5 | 100 |
| **Q8BFY9** | Transportin-1 OS=Mus musculus OX=10090 GN=Tnpo1 PE=1 SV=2 | 3,048 | 1 | 102,3 | 100 |
| **P62305** | Small nuclear ribonucleoprotein E OS=Mus musculus OX=10090 GN=Snrpe PE=1 SV=1 | 3,003 | 1 | 10,8 | 100 |
| **P62843** | 40S ribosomal protein S15 OS=Mus musculus OX=10090 GN=Rps15 PE=1 SV=2 | 2,993 | 1 | 17 | 100 |
| **A0A087WQT6** | Caspase-8 OS=Mus musculus OX=10090 GN=Casp8 PE=1 SV=1 | 2,985 | 1 | 57,8 | 100 |

| | | | | | |
|---|---|---|---|---|---|
| **A2A7S7** | Tyrosine--tRNA ligase OS=Mus musculus OX=10090 GN=Yars PE=1 SV=1 | 2,937 | 1 | 63 | 100 |
| **Q9CQW1** | Synaptobrevin homolog YKT6 OS=Mus musculus OX=10090 GN=Ykt6 PE=1 SV=1 | 2,91 | 1 | 22,3 | 100 |
| **P36536** | GTP-binding protein SAR1a OS=Mus musculus OX=10090 GN=Sar1a PE=1 SV=1 | 2,88 | 1 | 22,4 | 100 |
| **P61164** | Alpha-centractin OS=Mus musculus OX=10090 GN=Actr1a PE=1 SV=1 | 2,725 | 1 | 42,6 | 100 |
| **Q3T9Y8** | Eukaryotic translation initiation factor 3 subunit I OS=Mus musculus OX=10090 GN=Eif3i PE=2 SV=1 | 2,714 | 2 | 36,4 | 100 |
| **F8WHL2** | Coatomer subunit alpha OS=Mus musculus OX=10090 GN=Copa PE=1 SV=1 | 2,698 | 1 | 139,3 | 100 |
| **A2RTI3** | Legumain OS=Mus musculus OX=10090 GN=Lgmn PE=1 SV=1 | 2,681 | 1 | 49,3 | 100 |
| **A7VMV2** | Crystallin, lamda 1, isoform CRA_a OS=Mus musculus OX=10090 GN=Cryl1 PE=1 SV=1 | 2,665 | 1 | 35,2 | 100 |
| **P07310** | Creatine kinase M-type OS=Mus musculus OX=10090 GN=Ckm PE=1 SV=1 | 2,652 | 1 | 43 | 100 |
| **P32921** | Tryptophan--tRNA ligase, cytoplasmic OS=Mus musculus OX=10090 GN=Wars PE=1 SV=2 | 2,647 | 1 | 54,3 | 100 |
| **Q5SQX6** | Cytoplasmic FMR1-interacting protein 2 OS=Mus musculus OX=10090 GN=Cyfip2 PE=1 SV=2 | 2,618 | 1 | 145,6 | 100 |
| **Q4KMM3** | Oxidation resistance protein 1 OS=Mus musculus OX=10090 GN=Oxr1 PE=1 SV=3 | 2,58 | 2 | 95,9 | 100 |
| **P19157** | Glutathione S-transferase P 1 OS=Mus musculus OX=10090 GN=Gstp1 PE=1 SV=2 | 2,558 | 1 | 23,6 | 100 |
| **Q80UT7** | Rpl7a protein (Fragment) OS=Mus musculus OX=10090 GN=Rpl7a PE=2 SV=1 | 2,518 | 1 | 30,5 | 100 |
| **Q61704** | Inter-alpha-trypsin inhibitor heavy chain H3 OS=Mus musculus OX=10090 GN=Itih3 PE=1 SV=3 | 2,507 | 1 | 99,3 | 100 |
| **Q8VE43** | Meteorin-like protein OS=Mus musculus OX=10090 GN=Metrnl PE=1 SV=1 | 2,484 | 1 | 34,5 | 100 |
| **Q3TI98** | Uncharacterized protein OS=Mus musculus OX=10090 GN=Farsb PE=2 SV=1 | 2,465 | 1 | 65,8 | 100 |
| **Q3TKM9** | Actin-related protein 2/3 complex subunit 5 OS=Mus musculus OX=10090 GN=Arpc5 PE=2 SV=1 | 2,442 | 1 | 16,2 | 100 |
| **Q5M9L1** | 60S ribosomal protein L36 OS=Mus musculus OX=10090 GN=Rpl36 PE=2 SV=1 | 2,439 | 1 | 12,3 | 100 |
| **Q60866** | Phosphotriesterase-related protein OS=Mus musculus OX=10090 GN=Pter PE=1 SV=1 | 2,396 | 1 | 39,2 | 100 |

| | | | | | |
|---|---|---|---|---|---|
| **P61979-2** | Isoform 2 of Heterogeneous nuclear ribonucleoprotein K OS=Mus musculus OX=10090 GN=Hnrnpk | 2,363 | 1 | 51 | 100 |
| **Q61166** | Microtubule-associated protein RP/EB family member 1 OS=Mus musculus OX=10090 GN=Mapre1 PE=1 SV=3 | 2,32 | 1 | 30 | 100 |
| **Q3UWW9** | Uncharacterized protein OS=Mus musculus OX=10090 GN=Psmd11 PE=2 SV=1 | 2,296 | 1 | 47,3 | 100 |
| **Q99L45** | Eukaryotic translation initiation factor 2 subunit 2 OS=Mus musculus OX=10090 GN=Eif2s2 PE=1 SV=1 | 2,142 | 1 | 38,1 | 100 |
| **Q3TK95** | Uncharacterized protein OS=Mus musculus OX=10090 GN=Eif4e PE=1 SV=1 | 2,101 | 1 | 25 | 100 |
| **Q790Y8** | Glucose-6-phosphate 1-dehydrogenase OS=Mus musculus OX=10090 GN=G6pdx PE=1 SV=1 | 2,072 | 1 | 59,2 | 100 |
| **Q80X81** | Acetyl-Coenzyme A acetyltransferase 3 OS=Mus musculus OX=10090 GN=Acat3 PE=1 SV=1 | 2,008 | 1 | 41,4 | 100 |
| **P97484** | Leukocyte immunoglobulin-like receptor subfamily B member 3 OS=Mus musculus OX=10090 GN=Lilrb3 PE=1 SV=1 | 1,992 | 1 | 93 | 100 |
| **Q3TED1** | S-adenosylmethionine synthase OS=Mus musculus OX=10090 GN=Mat2a PE=2 SV=1 | 1,988 | 1 | 43,6 | 100 |
| **O89023** | Tripeptidyl-peptidase 1 OS=Mus musculus OX=10090 GN=Tpp1 PE=1 SV=2 | 1,888 | 1 | 61,3 | 100 |
| **Q571J7** | Serine/threonine-protein phosphatase 2A 55 kDa regulatory subunit B (Fragment) OS=Mus musculus OX=10090 GN=Ppp2r2a PE=2 SV=1 | 1,882 | 1 | 58,4 | 100 |
| **Q3TW74** | Uncharacterized protein OS=Mus musculus OX=10090 GN=Mthfd1 PE=2 SV=1 | 1,873 | 1 | 101,1 | 100 |
| **P26443** | Glutamate dehydrogenase 1, mitochondrial OS=Mus musculus OX=10090 GN=Glud1 PE=1 SV=1 | 1,859 | 2 | 61,3 | 100 |
| **Q3U3T6** | Uncharacterized protein OS=Mus musculus OX=10090 GN=Stk24 PE=2 SV=1 | 1,848 | 1 | 47,9 | 100 |
| **P61202-2** | Isoform 2 of COP9 signalosome complex subunit 2 OS=Mus musculus OX=10090 GN=Cops2 | 1,812 | 1 | 52,4 | 100 |
| **P97315** | Cysteine and glycine-rich protein 1 OS=Mus musculus OX=10090 GN=Csrp1 PE=1 SV=3 | 1,812 | 1 | 20,6 | 100 |
| **Q8K2Q9** | Shootin-1 OS=Mus musculus OX=10090 GN=Shtn1 PE=1 SV=1 | 1,785 | 1 | 71,3 | 100 |
| **D3YWJ3** | 40S ribosomal protein S2 OS=Mus musculus OX=10090 GN=Rps2 PE=1 SV=1 | 1,775 | 1 | 32,1 | 100 |
| **A1A4T2** | Alpha glucosidase 2 alpha neutral subunit OS=Mus musculus OX=10090 GN=Ganab PE=2 SV=1 | 1,754 | 1 | 109,3 | 100 |

| | | | | | |
|---|---|---|---|---|---|
| **P54823** | Probable ATP-dependent RNA helicase DDX6 OS=Mus musculus OX=10090 GN=Ddx6 PE=1 SV=1 | 1,719 | 1 | 54,2 | 100 |
| **Q8BG07** | Phospholipase D4 OS=Mus musculus OX=10090 GN=Pld4 PE=1 SV=1 | 1,678 | 1 | 56,1 | 100 |
| **Q7TMK9** | Heterogeneous nuclear ribonucleoprotein Q OS=Mus musculus OX=10090 GN=Syncrip PE=1 SV=2 | 1,671 | 1 | 69,6 | 100 |
| **Q6NZJ6** | Eukaryotic translation initiation factor 4 gamma 1 OS=Mus musculus OX=10090 GN=Eif4g1 PE=1 SV=1 | 1,534 | 1 | 176 | 100 |
| **Q3UZG3** | Uncharacterized protein OS=Mus musculus OX=10090 GN=Hnrnpa3 PE=2 SV=1 | 1,517 | 1 | 39,7 | 100 |
| **O89079** | Coatomer subunit epsilon OS=Mus musculus OX=10090 GN=Cope PE=1 SV=3 | 1,49 | 1 | 34,5 | 100 |
| **Q8VCF1-3** | Isoform 3 of Soluble calcium-activated nucleotidase 1 OS=Mus musculus OX=10090 GN=Cant1 | 1,444 | 1 | 49,5 | 100 |
| **P99029** | Peroxiredoxin-5, mitochondrial OS=Mus musculus OX=10090 GN=Prdx5 PE=1 SV=2 | 1,436 | 1 | 21,9 | 100 |
| **A0A1B0GS58** | Glutaredoxin-3 OS=Mus musculus OX=10090 GN=Glrx3 PE=1 SV=1 | 1,424 | 1 | 41,7 | 100 |
| **P38647** | Stress-70 protein, mitochondrial OS=Mus musculus OX=10090 GN=Hspa9 PE=1 SV=3 | 1,406 | 1 | 73,4 | 100 |
| **Q32KG4-2** | Isoform 2 of Retrotransposon Gag-like protein 9 OS=Mus musculus OX=10090 GN=Rtl9 | 1,382 | 1 | 141,7 | 100 |
| **F6XI62** | 60S ribosomal protein L7 (Fragment) OS=Mus musculus OX=10090 GN=Rpl7 PE=1 SV=1 | 1,344 | 1 | 32,5 | 100 |
| **Q03366** | C-C motif chemokine 7 OS=Mus musculus OX=10090 GN=Ccl7 PE=3 SV=1 | 1,339 | 1 | 11 | 100 |
| **Q3UZ39** | Leucine-rich repeat flightless-interacting protein 1 OS=Mus musculus OX=10090 GN=Lrrfip1 PE=1 SV=2 | 1,314 | 1 | 79,2 | 100 |
| **Q9D2V7** | Coronin-7 OS=Mus musculus OX=10090 GN=Coro7 PE=1 SV=2 | 1,296 | 1 | 100,7 | 100 |
| **P58389** | Serine/threonine-protein phosphatase 2A activator OS=Mus musculus OX=10090 GN=Ptpa PE=1 SV=1 | 1,268 | 1 | 36,7 | 100 |
| **Q64152** | Transcription factor BTF3 OS=Mus musculus OX=10090 GN=Btf3 PE=1 SV=3 | 1,26 | 1 | 22 | 100 |
| **A0A0R4J0G6** | Glutaminyl-peptide cyclotransferase OS=Mus musculus OX=10090 GN=Qpct PE=1 SV=1 | 1,258 | 1 | 41,1 | 100 |
| **Q09163** | Protein delta homolog 1 OS=Mus musculus OX=10090 GN=Dlk1 PE=1 SV=1 | 1,24 | 1 | 41,3 | 100 |
| **Q8BT06** | Tetraspanin OS=Mus musculus OX=10090 GN=Cd63 PE=2 SV=1 | 1,219 | 1 | 26,8 | 100 |

| | | | | | |
|---|---|---|---|---|---|
| **P30412** | Peptidyl-prolyl cis-trans isomerase C OS=Mus musculus OX=10090 GN=Ppic PE=1 SV=1 | 1,208 | 1 | 22,8 | 100 |
| **Q9Z1T2** | Thrombospondin-4 OS=Mus musculus OX=10090 GN=Thbs4 PE=1 SV=1 | 1,196 | 1 | 106,3 | 100 |
| **P56959** | RNA-binding protein FUS OS=Mus musculus OX=10090 GN=Fus PE=1 SV=1 | 1,172 | 1 | 52,6 | 100 |
| **G3UY93** | Valine--tRNA ligase (Fragment) OS=Mus musculus OX=10090 GN=Vars PE=1 SV=1 | 1,152 | 1 | 141,3 | 100 |
| **H3BJW3** | Cleavage and polyadenylation-specificity factor subunit 6 OS=Mus musculus OX=10090 GN=Cpsf6 PE=1 SV=1 | 1,145 | 1 | 63,4 | 100 |
| **Q3UNH3** | Uncharacterized protein (Fragment) OS=Mus musculus OX=10090 GN=Txnl1 PE=2 SV=1 | 1,116 | 1 | 33,5 | 100 |
| **P21956** | Lactadherin OS=Mus musculus OX=10090 GN=Mfge8 PE=1 SV=3 | 1,076 | 1 | 51,2 | 100 |
| **A0A1B0GSU0** | Aldehyde dehydrogenase family 16 member A1 OS=Mus musculus OX=10090 GN=Aldh16a1 PE=1 SV=1 | 1,023 | 1 | 84,7 | 100 |
| **Q5I0T8** | Ribosomal protein L19 OS=Mus musculus OX=10090 GN=Rpl19 PE=1 SV=1 | 0,939 | 1 | 23,5 | 100 |
| **O35226-2** | Isoform Rpn10B of 26S proteasome non-ATPase regulatory subunit 4 OS=Mus musculus OX=10090 GN=Psmd4 | 0,89 | 1 | 41 | 100 |
| **Q3TJQ7** | Alpha-1,4 glucan phosphorylase OS=Mus musculus OX=10090 GN=Pygl PE=2 SV=1 | 0,85 | 1 | 97,4 | 100 |
| **O70499** | Small nuclear ribonucleoprotein-associated protein OS=Mus musculus OX=10090 GN=Snrpn PE=3 SV=1 | 0,846 | 1 | 24,6 | 100 |
| **Q544F6** | Cotl1 protein OS=Mus musculus OX=10090 GN=Cotl1 PE=1 SV=1 | 29,64 | 5 | 15,9 | 20,185 |
| **Q3TIQ3** | Uncharacterized protein (Fragment) OS=Mus musculus OX=10090 GN=Pitpna PE=2 SV=1 | 3,695 | 1 | 40,2 | 18,849 |
| **Q62426** | Cystatin-B OS=Mus musculus OX=10090 GN=Cstb PE=1 SV=1 | 8,154 | 3 | 11 | 17,011 |
| **P51670** | C-C motif chemokine 9 OS=Mus musculus OX=10090 GN=Ccl9 PE=1 SV=1 | 9,507 | 2 | 13,9 | 15,692 |
| **P29416** | Beta-hexosaminidase subunit alpha OS=Mus musculus OX=10090 GN=Hexa PE=1 SV=2 | 62,146 | 10 | 60,6 | 15,277 |
| **P68033** | Actin, alpha cardiac muscle 1 OS=Mus musculus OX=10090 GN=Actc1 PE=1 SV=1 | 48,809 | 2 | 42 | 15,211 |
| **P97449** | Aminopeptidase N OS=Mus musculus OX=10090 GN=Anpep PE=1 SV=4 | 10,453 | 2 | 109,6 | 14,835 |
| **Q3V117** | ATP-citrate synthase OS=Mus musculus OX=10090 GN=Acly PE=1 SV=1 | 2,984 | 1 | 120,7 | 13,844 |

| | | | | | |
|---|---|---|---|---|---|
| **Q921I1** | Serotransferrin OS=Mus musculus OX=10090 GN=Tf PE=1 SV=1 | 21,442 | 7 | 76,7 | 13,562 |
| **P63242** | Eukaryotic translation initiation factor 5A-1 OS=Mus musculus OX=10090 GN=Eif5a PE=1 SV=2 | 43,527 | 4 | 16,8 | 12,427 |
| **P00920** | Carbonic anhydrase 2 OS=Mus musculus OX=10090 GN=Ca2 PE=1 SV=4 | 18,688 | 5 | 29 | 11,162 |
| **Q1XID4** | Renin/prorenin receptor OS=Mus musculus OX=10090 GN=Atp6ap2 PE=1 SV=2 | 13,053 | 2 | 39,1 | 11,088 |
| **Q571M4** | MKIAA4014 protein (Fragment) OS=Mus musculus OX=10090 GN=Akr1c13 PE=2 SV=1 | 1,579 | 1 | 37,3 | 10,41 |
| **Q3U561** | Ribosomal protein OS=Mus musculus OX=10090 GN=Rpl10a PE=2 SV=1 | 1,437 | 1 | 24,8 | 10,309 |
| **P63028** | Translationally-controlled tumor protein OS=Mus musculus OX=10090 GN=Tpt1 PE=1 SV=1 | 5,871 | 2 | 19,5 | 9,451 |
| **P11440** | Cyclin-dependent kinase 1 OS=Mus musculus OX=10090 GN=Cdk1 PE=1 SV=3 | 0,825 | 1 | 34,1 | 9,399 |
| **Q04447** | Creatine kinase B-type OS=Mus musculus OX=10090 GN=Ckb PE=1 SV=1 | 203,431 | 15 | 42,7 | 9,396 |
| **Q9Z0P5** | Twinfilin-2 OS=Mus musculus OX=10090 GN=Twf2 PE=1 SV=1 | 26,02 | 5 | 39,4 | 9,33 |
| **A0A338P7K4** | 40S ribosomal protein S10 OS=Mus musculus OX=10090 GN=Rps10 PE=1 SV=1 | 2,676 | 1 | 19,2 | 9,131 |
| **Q6IRU2** | Tropomyosin alpha-4 chain OS=Mus musculus OX=10090 GN=Tpm4 PE=1 SV=3 | 6,18 | 2 | 28,5 | 8,71 |
| **Q4FK54** | Proteaseome (Prosome, macropain) 28 subunit, 3 OS=Mus musculus OX=10090 GN=Psme3 PE=1 SV=1 | 3,581 | 1 | 29,5 | 8,255 |
| **Q561N4** | MCG1032217 OS=Mus musculus OX=10090 GN=Ube2l3 PE=1 SV=1 | 31,771 | 4 | 17,9 | 8,144 |
| **P07901** | Heat shock protein HSP 90-alpha OS=Mus musculus OX=10090 GN=Hsp90aa1 PE=1 SV=4 | 39,651 | 5 | 84,7 | 7,886 |
| **O35286** | Pre-mRNA-splicing factor ATP-dependent RNA helicase DHX15 OS=Mus musculus OX=10090 GN=Dhx15 PE=1 SV=2 | 1,693 | 1 | 90,9 | 7,798 |
| **A1L353** | Transforming growth factor, beta induced OS=Mus musculus OX=10090 GN=Tgfbi PE=1 SV=1 | 33,204 | 7 | 74,6 | 7,433 |
| **A0A0R4J1E2** | Elongation factor 1-delta OS=Mus musculus OX=10090 GN=Eef1d PE=1 SV=1 | 2,957 | 1 | 72,9 | 7,34 |
| **A0A140T8V5** | Proliferating cell nuclear antigen OS=Mus musculus OX=10090 GN=Pcna-ps2 PE=3 SV=1 | 3,722 | 1 | 28,8 | 7,139 |
| **A0A0J9YUL3** | Septin 11, isoform CRA_b OS=Mus musculus OX=10090 GN=Sept11 PE=1 SV=1 | 3,699 | 2 | 49,8 | 7,07 |
| **B2RTB0** | MCG17262 OS=Mus musculus OX=10090 GN=Pdap1 PE=1 SV=1 | 1,311 | 1 | 20,6 | 6,897 |
| **Q3UMM1** | Tubulin beta chain OS=Mus musculus OX=10090 GN=Tubb6 PE=1 SV=1 | 8,29 | 2 | 50,1 | 6,644 |

| | | | | | |
|---|---|---|---|---|---|
| **A0A0R4J0I9** | Low density lipoprotein receptor-related protein 1 OS=Mus musculus OX=10090 GN=Lrp1 PE=1 SV=1 | 47,635 | 15 | 504,4 | 6,576 |
| **P50516** | V-type proton ATPase catalytic subunit A OS=Mus musculus OX=10090 GN=Atp6v1a PE=1 SV=2 | 62,139 | 11 | 68,3 | 6,505 |
| **Q8BU30** | Isoleucine--tRNA ligase, cytoplasmic OS=Mus musculus OX=10090 GN=Iars PE=1 SV=2 | 7,313 | 3 | 144,2 | 6,49 |
| **P16110** | Galectin-3 OS=Mus musculus OX=10090 GN=Lgals3 PE=1 SV=3 | 29,486 | 6 | 27,5 | 6,474 |
| **Q62059** | Versican core protein OS=Mus musculus OX=10090 GN=Vcan PE=1 SV=2 | 3,967 | 1 | 366,6 | 6,422 |
| **Q05117** | Tartrate-resistant acid phosphatase type 5 OS=Mus musculus OX=10090 GN=Acp5 PE=1 SV=2 | 36,835 | 7 | 36,8 | 6,368 |
| **Q76MZ3** | Serine/threonine-protein phosphatase 2A 65 kDa regulatory subunit A alpha isoform OS=Mus musculus OX=10090 GN=Ppp2r1a PE=1 SV=3 | 20,387 | 5 | 65,3 | 6,355 |
| **E0CYM8** | Tyrosine-protein phosphatase non-receptor type substrate 1 OS=Mus musculus OX=10090 GN=Sirpa PE=1 SV=1 | 8,516 | 3 | 56,4 | 6,265 |
| **P62334** | 26S proteasome regulatory subunit 10B OS=Mus musculus OX=10090 GN=Psmc6 PE=1 SV=1 | 4,959 | 1 | 44,1 | 6,252 |
| **P11680** | Properdin OS=Mus musculus OX=10090 GN=Cfp PE=2 SV=2 | 5,893 | 2 | 50,3 | 6,154 |
| **Q6ZWN5** | 40S ribosomal protein S9 OS=Mus musculus OX=10090 GN=Rps9 PE=1 SV=3 | 1,118 | 1 | 22,6 | 6,144 |
| **Q6ZPS9** | MKIAA1375 protein (Fragment) OS=Mus musculus OX=10090 GN=Pdcd6ip PE=2 SV=1 | 14,963 | 4 | 100,1 | 6,132 |
| **P63330** | Serine/threonine-protein phosphatase 2A catalytic subunit alpha isoform OS=Mus musculus OX=10090 GN=Ppp2ca PE=1 SV=1 | 11,324 | 4 | 35,6 | 5,899 |
| **Q3TVV6** | Uncharacterized protein OS=Mus musculus OX=10090 GN=Hnrnpu PE=2 SV=1 | 3,519 | 1 | 87,9 | 5,743 |
| **P55065** | Phospholipid transfer protein OS=Mus musculus OX=10090 GN=Pltp PE=1 SV=1 | 27,499 | 6 | 54,4 | 5,605 |
| **P50543** | Protein S100-A11 OS=Mus musculus OX=10090 GN=S100a11 PE=1 SV=1 | 4,411 | 1 | 11,1 | 5,424 |
| **P11983** | T-complex protein 1 subunit alpha OS=Mus musculus OX=10090 GN=Tcp1 PE=1 SV=3 | 9,765 | 3 | 60,4 | 5,378 |
| **P13020** | Gelsolin OS=Mus musculus OX=10090 GN=Gsn PE=1 SV=3 | 85,826 | 14 | 85,9 | 5,36 |
| **Q3TQR3** | MCG118037 OS=Mus musculus OX=10090 GN=Eif5 PE=1 SV=1 | 2,729 | 1 | 48,9 | 5,324 |
| **Q6RI64** | Proteasome subunit beta OS=Mus musculus OX=10090 GN=Psmb1 PE=1 SV=1 | 4,578 | 3 | 26,4 | 5,322 |

| | | | | |
|---|---|---|---|---|
| **Q3TF87** | Uncharacterized protein OS=Mus musculus OX=10090 GN=Dars PE=2 SV=1 | 1,095 | 1 | 57,1 | 5,266 |
| **Q3UGR5** | Haloacid dehalogenase-like hydrolase domain-containing protein 2 OS=Mus musculus OX=10090 GN=Hdhd2 PE=1 SV=2 | 2,763 | 2 | 28,7 | 5,108 |
| **Q9WVA4** | Transgelin-2 OS=Mus musculus OX=10090 GN=Tagln2 PE=1 SV=4 | 36,722 | 7 | 22,4 | 5,01 |
| **P34960** | Macrophage metalloelastase OS=Mus musculus OX=10090 GN=Mmp12 PE=1 SV=3 | 5,216 | 3 | 54,9 | 5,002 |
| **A0A1Y7VKY1** | MCG116671 OS=Mus musculus OX=10090 GN=Gm11361 PE=3 SV=1 | 4,123 | 2 | 17,7 | 4,973 |
| **A2APM2** | CD44 antigen OS=Mus musculus OX=10090 GN=Cd44 PE=1 SV=1 | 3,718 | 1 | 85,8 | 4,932 |
| **P62897** | Cytochrome c, somatic OS=Mus musculus OX=10090 GN=Cycs PE=1 SV=2 | 5,847 | 2 | 11,6 | 4,861 |
| **P53996** | Cellular nucleic acid-binding protein OS=Mus musculus OX=10090 GN=Cnbp PE=1 SV=2 | 8,575 | 1 | 19,6 | 4,85 |
| **E9QPX1** | Collagen alpha-1(XVIII) chain OS=Mus musculus OX=10090 GN=Col18a1 PE=1 SV=1 | 4,056 | 2 | 182,2 | 4,791 |
| **Q3U1U4** | Predicted gene, 49368 OS=Mus musculus OX=10090 GN=Gm49368 PE=1 SV=1 | 36,065 | 10 | 135,9 | 4,766 |
| **Q3TJG6** | Uncharacterized protein OS=Mus musculus OX=10090 GN=Ptges3 PE=2 SV=1 | 1,866 | 1 | 18,7 | 4,751 |
| **Q8VGW5** | Olfactory receptor OS=Mus musculus OX=10090 GN=Olfr691 PE=2 SV=1 | 1,999 | 1 | 36,2 | 4,746 |
| **P01887** | Beta-2-microglobulin OS=Mus musculus OX=10090 GN=B2m PE=1 SV=2 | 1,268 | 1 | 13,8 | 4,746 |
| **P29351-2** | Isoform 2 of Tyrosine-protein phosphatase non-receptor type 6 OS=Mus musculus OX=10090 GN=Ptpn6 | 2,303 | 1 | 67,7 | 4,663 |
| **P10605** | Cathepsin B OS=Mus musculus OX=10090 GN=Ctsb PE=1 SV=2 | 68,797 | 9 | 37,3 | 4,644 |
| **P61205** | ADP-ribosylation factor 3 OS=Mus musculus OX=10090 GN=Arf3 PE=2 SV=2 | 15,568 | 4 | 20,6 | 4,592 |
| **Q3TG21** | V-type proton ATPase subunit C OS=Mus musculus OX=10090 GN=Atp6v1c1 PE=2 SV=1 | 34,366 | 5 | 43,8 | 4,556 |
| **O35375** | Neuropilin-2 OS=Mus musculus OX=10090 GN=Nrp2 PE=1 SV=2 | 21,937 | 7 | 104,6 | 4,546 |
| **Q8C243** | Uncharacterized protein OS=Mus musculus OX=10090 GN=Ctsd PE=2 SV=1 | 47,628 | 9 | 48,3 | 4,523 |
| **Q6ZQ38** | Cullin-associated NEDD8-dissociated protein 1 OS=Mus musculus OX=10090 GN=Cand1 PE=1 SV=2 | 7,709 | 3 | 136,2 | 4,518 |
| **E9PZF0** | Nucleoside diphosphate kinase OS=Mus musculus OX=10090 GN=Gm20390 PE=3 SV=1 | 41,165 | 10 | 30,2 | 4,517 |
| **Q8BKC5** | Importin-5 OS=Mus musculus OX=10090 GN=Ipo5 PE=1 SV=3 | 16,46 | 7 | 123,5 | 4,485 |

| | | | | | |
|---|---|---|---|---|---|
| **Q3U7Z6** | Phosphoglycerate mutase OS=Mus musculus OX=10090 GN=Pgam1 PE=1 SV=1 | 115,782 | 16 | 28,8 | 4,447 |
| **P45377** | Aldose reductase-related protein 2 OS=Mus musculus OX=10090 GN=Akr1b8 PE=1 SV=2 | 18,629 | 5 | 36,1 | 4,378 |
| **B7ZWC4** | Insulin-like growth factor 2 receptor OS=Mus musculus OX=10090 GN=Igf2r PE=2 SV=1 | 12,09 | 6 | 273,7 | 4,287 |
| **Q8BVK3** | Uncharacterized protein (Fragment) OS=Mus musculus OX=10090 GN=Uap1l1 PE=2 SV=1 | 13,813 | 4 | 56,8 | 4,231 |
| **P11835** | Integrin beta-2 OS=Mus musculus OX=10090 GN=Itgb2 PE=1 SV=2 | 19,12 | 5 | 85 | 4,222 |
| **Q3TT75** | Uncharacterized protein OS=Mus musculus OX=10090 GN=Ctsl PE=2 SV=1 | 38,468 | 7 | 37,5 | 4,221 |
| **Q3THQ5** | Uncharacterized protein OS=Mus musculus OX=10090 GN=Stip1 PE=2 SV=1 | 6,131 | 1 | 62,5 | 4,22 |
| **Q60676** | Serine/threonine-protein phosphatase 5 OS=Mus musculus OX=10090 GN=Ppp5c PE=1 SV=3 | 4,94 | 1 | 56,8 | 4,182 |
| **P25085** | Interleukin-1 receptor antagonist protein OS=Mus musculus OX=10090 GN=Il1rn PE=2 SV=1 | 10,202 | 2 | 20,3 | 4,123 |
| **Q07797** | Galectin-3-binding protein OS=Mus musculus OX=10090 GN=Lgals3bp PE=1 SV=1 | 21,205 | 4 | 64,5 | 4,121 |
| **Q542D9** | Transferrin receptor, isoform CRA_a OS=Mus musculus OX=10090 GN=Tfrc PE=1 SV=1 | 58,143 | 16 | 85,7 | 4,118 |
| **Q3UUH0** | Uncharacterized protein OS=Mus musculus OX=10090 GN=Asl PE=2 SV=1 | 6,443 | 2 | 51,7 | 4,086 |
| **Q8K1X5** | EH-domain containing 1 (Fragment) OS=Mus musculus OX=10090 GN=Ehd1 PE=2 SV=2 | 21,548 | 5 | 61,9 | 3,966 |
| **P17439** | Glucosylceramidase OS=Mus musculus OX=10090 GN=Gba PE=1 SV=1 | 1,549 | 1 | 57,6 | 3,936 |
| **P29341** | Polyadenylate-binding protein 1 OS=Mus musculus OX=10090 GN=Pabpc1 PE=1 SV=2 | 11,999 | 3 | 70,6 | 3,93 |
| **P47753** | F-actin-capping protein subunit alpha-1 OS=Mus musculus OX=10090 GN=Capza1 PE=1 SV=4 | 12,503 | 4 | 32,9 | 3,929 |
| **G5E8T9** | Hydroxyacyl glutathione hydrolase OS=Mus musculus OX=10090 GN=Hagh PE=1 SV=1 | 4,269 | 2 | 34,1 | 3,918 |
| **Q3U6P5** | Uncharacterized protein OS=Mus musculus OX=10090 GN=Hnrnpc PE=2 SV=1 | 8,475 | 3 | 36,9 | 3,911 |
| **F7DBB3** | AHNAK nucleoprotein 2 (Fragment) OS=Mus musculus OX=10090 GN=Ahnak2 PE=1 SV=1 | 1,753 | 1 | 166,4 | 3,899 |

| | | | | | |
|---|---|---|---|---|---|
| **P82343** | N-acylglucosamine 2-epimerase OS=Mus musculus OX=10090 GN=Renbp PE=1 SV=3 | 1,881 | 1 | 49,7 | 3,894 |
| **Q3UM14** | Deoxyribonuclease II alpha OS=Mus musculus OX=10090 GN=Dnase2a PE=1 SV=1 | 2,107 | 1 | 38,8 | 3,892 |
| **P70168** | Importin subunit beta-1 OS=Mus musculus OX=10090 GN=Kpnb1 PE=1 SV=2 | 25,069 | 5 | 97,1 | 3,88 |
| **B1AX58** | Plastin-3 OS=Mus musculus OX=10090 GN=Pls3 PE=1 SV=1 | 35,576 | 4 | 71,7 | 3,869 |
| **H3BKH6** | S-formylglutathione hydrolase OS=Mus musculus OX=10090 GN=Esd PE=1 SV=1 | 39,033 | 7 | 32,8 | 3,868 |
| **V9GWY0** | 40S ribosomal protein S4 OS=Mus musculus OX=10090 GN=Gm15013 PE=3 SV=1 | 2,122 | 1 | 29,9 | 3,868 |
| **A0A1W2P768** | Histone H3.2 OS=Mus musculus OX=10090 GN=Hist2h3c1 PE=1 SV=1 | 2,939 | 2 | 20,2 | 3,801 |
| **Q3UE92** | X-prolyl aminopeptidase (Aminopeptidase P) 1, soluble, isoform CRA_b OS=Mus musculus OX=10090 GN=Xpnpep1 PE=1 SV=1 | 4,077 | 2 | 74,5 | 3,792 |
| **Q8K1B8** | Fermitin family homolog 3 OS=Mus musculus OX=10090 GN=Fermt3 PE=1 SV=1 | 14,199 | 4 | 75,6 | 3,758 |
| **Q3UJ44** | Uncharacterized protein OS=Mus musculus OX=10090 GN=Capg PE=2 SV=1 | 66,99 | 11 | 38,6 | 3,717 |
| **Q3UCL7** | Uncharacterized protein OS=Mus musculus OX=10090 GN=Rps3 PE=2 SV=1 | 4,665 | 1 | 26,7 | 3,714 |
| **P21460** | Cystatin-C OS=Mus musculus OX=10090 GN=Cst3 PE=1 SV=2 | 21,466 | 5 | 15,5 | 3,707 |
| **Q3UA52** | Arp2/3 complex 34 kDa subunit (Fragment) OS=Mus musculus OX=10090 GN=Arpc2 PE=2 SV=1 | 31,253 | 8 | 42,5 | 3,695 |
| **Q99KC8** | von Willebrand factor A domain-containing protein 5A OS=Mus musculus OX=10090 GN=Vwa5a PE=1 SV=2 | 6,286 | 3 | 87,1 | 3,657 |
| **Q3U111** | Uncharacterized protein (Fragment) OS=Mus musculus OX=10090 GN=Rdx PE=2 SV=1 | 16,398 | 1 | 77,4 | 3,636 |
| **P62320** | Small nuclear ribonucleoprotein Sm D3 OS=Mus musculus OX=10090 GN=Snrpd3 PE=1 SV=1 | 2,367 | 1 | 13,9 | 3,578 |
| **Q71LX8** | Heat shock protein 84b OS=Mus musculus OX=10090 GN=Hsp90ab1 PE=1 SV=1 | 92,21 | 17 | 83,2 | 3,567 |
| **P16125** | L-lactate dehydrogenase B chain OS=Mus musculus OX=10090 GN=Ldhb PE=1 SV=2 | 19,185 | 4 | 36,5 | 3,548 |
| **O88958** | Glucosamine-6-phosphate isomerase 1 OS=Mus musculus OX=10090 GN=Gnpda1 PE=1 SV=3 | 11,068 | 3 | 32,5 | 3,541 |
| **Q3U4S9** | Coatomer subunit delta OS=Mus musculus OX=10090 GN=Arcn1 PE=2 SV=1 | 2,923 | 1 | 57,1 | 3,54 |

| | | | | | |
|---|---|---|---|---|---|
| **Q9CWJ9** | Bifunctional purine biosynthesis protein PURH OS=Mus musculus OX=10090 GN=Atic PE=1 SV=2 | 1,198 | 1 | 64,2 | 3,492 |
| **Q3U8F5** | Uncharacterized protein OS=Mus musculus OX=10090 GN=Psmd12 PE=2 SV=1 | 4,044 | 1 | 52,8 | 3,346 |
| **Q60865** | Caprin-1 OS=Mus musculus OX=10090 GN=Caprin1 PE=1 SV=2 | 2,33 | 2 | 78,1 | 3,325 |
| **A0A0A0MQA5** | Tubulin alpha chain (Fragment) OS=Mus musculus OX=10090 GN=Tuba4a PE=1 SV=1 | 68,197 | 2 | 52,9 | 3,306 |
| **Q3UJL7** | Uncharacterized protein OS=Mus musculus OX=10090 GN=Srm PE=2 SV=1 | 6,637 | 3 | 34 | 3,286 |
| **A0A1S6GWG6** | Uncharacterized protein OS=Mus musculus OX=10090 GN=Atp6v1b2 PE=2 SV=1 | 55,344 | 10 | 59,2 | 3,283 |
| **P62242** | 40S ribosomal protein S8 OS=Mus musculus OX=10090 GN=Rps8 PE=1 SV=2 | 10,989 | 4 | 24,2 | 3,279 |
| **Q9D0I9** | Arginine--tRNA ligase, cytoplasmic OS=Mus musculus OX=10090 GN=Rars PE=1 SV=2 | 3,417 | 2 | 75,6 | 3,252 |
| **P10639** | Thioredoxin OS=Mus musculus OX=10090 GN=Txn PE=1 SV=3 | 4,534 | 1 | 11,7 | 3,239 |
| **Q9R0E2** | Procollagen-lysine,2-oxoglutarate 5-dioxygenase 1 OS=Mus musculus OX=10090 GN=Plod1 PE=1 SV=1 | 15,307 | 5 | 83,5 | 3,235 |
| **Q7TPR4** | Alpha-actinin-1 OS=Mus musculus OX=10090 GN=Actn1 PE=1 SV=1 | 147,701 | 19 | 103 | 3,225 |
| **Q8BK67** | Protein RCC2 OS=Mus musculus OX=10090 GN=Rcc2 PE=1 SV=1 | 4,752 | 2 | 55,9 | 3,215 |
| **Q3U8D2** | Uncharacterized protein OS=Mus musculus OX=10090 GN=St13 PE=2 SV=1 | 3,998 | 2 | 41,5 | 3,202 |
| **Q3TWT5** | Uncharacterized protein OS=Mus musculus OX=10090 GN=Asah1 PE=2 SV=1 | 11,285 | 3 | 44,6 | 3,182 |
| **Q80TM2** | MKIAA1027 protein (Fragment) OS=Mus musculus OX=10090 GN=Tln1 PE=2 SV=4 | 140,839 | 24 | 272 | 3,174 |
| **P09581** | Macrophage colony-stimulating factor 1 receptor OS=Mus musculus OX=10090 GN=Csf1r PE=1 SV=3 | 13,054 | 5 | 109,1 | 3,158 |
| **P62806** | Histone H4 OS=Mus musculus OX=10090 GN=Hist1h4a PE=1 SV=2 | 21,345 | 4 | 11,4 | 3,148 |
| **P70670** | Nascent polypeptide-associated complex subunit alpha, muscle-specific form OS=Mus musculus OX=10090 GN=Naca PE=1 SV=2 | 2,668 | 1 | 220,4 | 3,136 |
| **P59999** | Actin-related protein 2/3 complex subunit 4 OS=Mus musculus OX=10090 GN=Arpc4 PE=1 SV=3 | 12,601 | 3 | 19,7 | 3,111 |
| **A0A0R4J092** | Beta-mannosidase OS=Mus musculus OX=10090 GN=Manba PE=1 SV=1 | 15,263 | 3 | 100,8 | 3,098 |
| **Q9WV02** | RNA-binding motif protein, X chromosome OS=Mus musculus OX=10090 GN=Rbmx PE=1 SV=1 | 0,957 | 1 | 42,3 | 3,092 |

| | | | | | |
|---|---|---|---|---|---|
| **Q3TEL5** | Lipase OS=Mus musculus OX=10090 GN=Lipa PE=2 SV=1 | 26,032 | 4 | 45,3 | 3,088 |
| **P08905** | Lysozyme C-2 OS=Mus musculus OX=10090 GN=Lyz2 PE=1 SV=2 | 51,764 | 5 | 16,7 | 3,085 |
| **P11152** | Lipoprotein lipase OS=Mus musculus OX=10090 GN=Lpl PE=1 SV=3 | 11,334 | 2 | 53,1 | 3,063 |
| **Q571E4** | N-acetylgalactosamine-6-sulfatase OS=Mus musculus OX=10090 GN=Galns PE=1 SV=2 | 6,632 | 2 | 57,6 | 3,049 |
| **O88569** | Heterogeneous nuclear ribonucleoproteins A2/B1 OS=Mus musculus OX=10090 GN=Hnrnpa2b1 PE=1 SV=2 | 19,456 | 6 | 37,4 | 3,046 |
| **O08529** | Calpain-2 catalytic subunit OS=Mus musculus OX=10090 GN=Capn2 PE=1 SV=4 | 10,533 | 3 | 79,8 | 3,044 |
| **Q8C845** | EF-hand domain-containing protein D2 OS=Mus musculus OX=10090 GN=Efhd2 PE=1 SV=1 | 4,949 | 3 | 26,8 | 3,02 |
| **P24369** | Peptidyl-prolyl cis-trans isomerase B OS=Mus musculus OX=10090 GN=Ppib PE=1 SV=2 | 13,985 | 3 | 23,7 | 3,016 |
| **Q543J5** | Antithrombin OS=Mus musculus OX=10090 GN=Serpinc1 PE=1 SV=1 | 19,28 | 4 | 52 | 2,992 |
| **P48678** | Prelamin-A/C OS=Mus musculus OX=10090 GN=Lmna PE=1 SV=2 | 48,375 | 16 | 74,2 | 2,983 |
| **Q91Z25** | Actin-related protein 2/3 complex subunit OS=Mus musculus OX=10090 GN=Arpc1b PE=1 SV=1 | 6,739 | 3 | 41,5 | 2,98 |
| **Q3TDF8** | Uncharacterized protein OS=Mus musculus OX=10090 GN=Etf1 PE=2 SV=1 | 3,194 | 1 | 49 | 2,976 |
| **Q4VAI2** | Acid phosphatase 1, soluble OS=Mus musculus OX=10090 GN=Acp1 PE=2 SV=1 | 4,188 | 2 | 18,2 | 2,966 |
| **O70310** | Glycylpeptide N-tetradecanoyltransferase 1 OS=Mus musculus OX=10090 GN=Nmt1 PE=1 SV=1 | 1,574 | 1 | 56,9 | 2,939 |
| **Q9JKB3** | Y-box-binding protein 3 OS=Mus musculus OX=10090 GN=Ybx3 PE=1 SV=2 | 2,607 | 1 | 38,8 | 2,926 |
| **P11276** | Fibronectin OS=Mus musculus OX=10090 GN=Fn1 PE=1 SV=4 | 3,078 | 1 | 272,4 | 2,922 |
| **Q99PT1** | Rho GDP-dissociation inhibitor 1 OS=Mus musculus OX=10090 GN=Arhgdia PE=1 SV=3 | 41,041 | 5 | 23,4 | 2,914 |
| **P58252** | Elongation factor 2 OS=Mus musculus OX=10090 GN=Eef2 PE=1 SV=2 | 161,033 | 24 | 95,3 | 2,901 |
| **Q64426** | Histone H2A (Fragment) OS=Mus musculus domesticus OX=10092 GN=H2A PE=2 SV=1 | 20,865 | 3 | 14,7 | 2,9 |
| **Q3TW77** | Uncharacterized protein OS=Mus musculus OX=10090 GN=Grn PE=2 SV=1 | 2,553 | 1 | 65 | 2,894 |
| **P08228** | Superoxide dismutase [Cu-Zn] OS=Mus musculus OX=10090 GN=Sod1 PE=1 SV=2 | 7,9 | 2 | 15,9 | 2,876 |

| | | | | | |
|---|---|---|---|---|---|
| **Q3TJ94** | Prothrombin OS=Mus musculus OX=10090 GN=F2 PE=1 SV=1 | 16,167 | 4 | 70,2 | 2,873 |
| **O89020-3** | Isoform 3 of Afamin OS=Mus musculus OX=10090 GN=Afm | 1,019 | 1 | 69,6 | 2,844 |
| **P55144-2** | Isoform 2 of Tyrosine-protein kinase receptor TYRO3 OS=Mus musculus OX=10090 GN=Tyro3 | 1,046 | 1 | 97,3 | 2,843 |
| **Q3U850** | Uncharacterized protein OS=Mus musculus OX=10090 GN=Rpl5 PE=2 SV=1 | 2,123 | 1 | 34,3 | 2,842 |
| **Q8BVQ9** | 26S proteasome regulatory subunit 7 OS=Mus musculus OX=10090 GN=Psmc2 PE=1 SV=1 | 8,489 | 2 | 52,8 | 2,837 |
| **E9PYH2** | Cytosolic acyl coenzyme A thioester hydrolase OS=Mus musculus OX=10090 GN=Acot7 PE=1 SV=1 | 3,134 | 1 | 42,8 | 2,837 |
| **Q3UD67** | Uncharacterized protein OS=Mus musculus OX=10090 GN=Aars PE=2 SV=1 | 16,168 | 4 | 106,8 | 2,83 |
| **P14152** | Malate dehydrogenase, cytoplasmic OS=Mus musculus OX=10090 GN=Mdh1 PE=1 SV=3 | 39,59 | 8 | 36,5 | 2,805 |
| **B2RWX2** | Complement component 4B (Childo blood group) OS=Mus musculus OX=10090 GN=C4b PE=2 SV=1 | 2,69 | 2 | 192,8 | 2,786 |
| **O09131** | Glutathione S-transferase omega-1 OS=Mus musculus OX=10090 GN=Gsto1 PE=1 SV=2 | 16,042 | 4 | 27,5 | 2,752 |
| **Q3U890** | Uncharacterized protein OS=Mus musculus OX=10090 GN=Hars PE=2 SV=1 | 6,476 | 2 | 57,4 | 2,746 |
| **Q545T0** | Cathepsin K OS=Mus musculus OX=10090 GN=Ctsk PE=2 SV=1 | 25,668 | 7 | 36,9 | 2,739 |
| **P05201** | Aspartate aminotransferase, cytoplasmic OS=Mus musculus OX=10090 GN=Got1 PE=1 SV=3 | 40,887 | 11 | 46,2 | 2,721 |
| **E9PY39** | Predicted gene 20431 OS=Mus musculus OX=10090 GN=Gm20431 PE=4 SV=1 | 3,647 | 2 | 42,1 | 2,712 |
| **Q3ULT2** | Actinin alpha 4 OS=Mus musculus OX=10090 GN=Actn4 PE=1 SV=1 | 118,401 | 18 | 104,9 | 2,71 |
| **P40124** | Adenylyl cyclase-associated protein 1 OS=Mus musculus OX=10090 GN=Cap1 PE=1 SV=4 | 40,888 | 8 | 51,5 | 2,699 |
| **B2RUJ7** | Xanthine dehydrogenase OS=Mus musculus OX=10090 GN=Xdh PE=2 SV=1 | 5,633 | 2 | 146,5 | 2,693 |
| **Q8C2Q7** | Heterogeneous nuclear ribonucleoprotein H OS=Mus musculus OX=10090 GN=Hnrnph1 PE=1 SV=1 | 2,417 | 1 | 51,2 | 2,69 |
| **P63101** | 14-3-3 protein zeta/delta OS=Mus musculus OX=10090 GN=Ywhaz PE=1 SV=1 | 49,217 | 8 | 27,8 | 2,665 |
| **P60335** | Poly(rC)-binding protein 1 OS=Mus musculus OX=10090 GN=Pcbp1 PE=1 SV=1 | 1,2 | 1 | 37,5 | 2,663 |
| **Q8CGP2-2** | Isoform 2 of Histone H2B type 1-P OS=Mus musculus OX=10090 GN=Hist1h2bp | 12,1 | 3 | 15,6 | 2,655 |

| | | | | | |
|---|---|---|---|---|---|
| **Q61599** | Rho GDP-dissociation inhibitor 2 OS=Mus musculus OX=10090 GN=Arhgdib PE=1 SV=3 | 23,073 | 3 | 22,8 | 2,649 |
| **Q3TE63** | Peptidyl-prolyl cis-trans isomerase OS=Mus musculus OX=10090 GN=Ppia PE=2 SV=1 | 40,03 | 8 | 17,9 | 2,644 |
| **Q6A0F1** | MKIAA0002 protein (Fragment) OS=Mus musculus OX=10090 GN=Cct8 PE=2 SV=1 | 17,142 | 5 | 60,2 | 2,628 |
| **P19096** | Fatty acid synthase OS=Mus musculus OX=10090 GN=Fasn PE=1 SV=2 | 10,815 | 4 | 272,3 | 2,624 |
| **Q61233** | Plastin-2 OS=Mus musculus OX=10090 GN=Lcp1 PE=1 SV=4 | 190,42 | 26 | 70,1 | 2,61 |
| **Q9ERK4** | Exportin-2 OS=Mus musculus OX=10090 GN=Cse1l PE=1 SV=1 | 4,792 | 1 | 110,4 | 2,607 |
| **Q99LX0** | Protein/nucleic acid deglycase DJ-1 OS=Mus musculus OX=10090 GN=Park7 PE=1 SV=1 | 7,555 | 2 | 20 | 2,594 |
| **Q91V99** | 40S ribosomal protein S30 (Fragment) OS=Mus musculus OX=10090 GN=fau PE=3 SV=1 | 2,386 | 1 | 14,8 | 2,584 |
| **Q8BVE3** | V-type proton ATPase subunit H OS=Mus musculus OX=10090 GN=Atp6v1h PE=1 SV=1 | 10,933 | 4 | 55,8 | 2,578 |
| **P01027** | Complement C3 OS=Mus musculus OX=10090 GN=C3 PE=1 SV=3 | 29,472 | 11 | 186,4 | 2,566 |
| **Q3TIZ0** | Tubulin alpha chain OS=Mus musculus OX=10090 GN=Tuba1c PE=2 SV=1 | 77,962 | 5 | 49,9 | 2,554 |
| **Q5FWB6** | 60S acidic ribosomal protein P0 OS=Mus musculus OX=10090 GN=Rplp0 PE=2 SV=1 | 9,724 | 3 | 34,2 | 2,55 |
| **Q922Q8** | Leucine-rich repeat-containing protein 59 OS=Mus musculus OX=10090 GN=Lrrc59 PE=1 SV=1 | 8,244 | 3 | 34,9 | 2,544 |
| **A0A0R4J093** | UMP-CMP kinase OS=Mus musculus OX=10090 GN=Cmpk1 PE=1 SV=1 | 5,023 | 3 | 25,7 | 2,541 |
| **Q3UJP8** | Trifunctional purine biosynthetic protein adenosine-3 OS=Mus musculus OX=10090 GN=Gart PE=2 SV=1 | 2,456 | 1 | 107,4 | 2,528 |
| **B9EIU1** | Glutamyl-prolyl-tRNA synthetase OS=Mus musculus OX=10090 GN=Eprs PE=2 SV=1 | 7,072 | 3 | 169,9 | 2,525 |
| **P00493** | Hypoxanthine-guanine phosphoribosyltransferase OS=Mus musculus OX=10090 GN=Hprt1 PE=1 SV=3 | 17,107 | 3 | 24,6 | 2,491 |
| **A0A1S6GWH2** | Uncharacterized protein OS=Mus musculus OX=10090 GN=Ddx39b PE=2 SV=1 | 13,916 | 4 | 54,6 | 2,49 |
| **Q3UW40** | Uncharacterized protein OS=Mus musculus OX=10090 GN=Rpl24 PE=2 SV=1 | 1,712 | 1 | 18,2 | 2,486 |

| | | | | | |
|---|---|---|---|---|---|
| **Q01853** | Transitional endoplasmic reticulum ATPase OS=Mus musculus OX=10090 GN=Vcp PE=1 SV=4 | 68,396 | 16 | 89,3 | 2,467 |
| **Q01730** | Ras suppressor protein 1 OS=Mus musculus OX=10090 GN=Rsu1 PE=1 SV=3 | 1,805 | 2 | 31,5 | 2,463 |
| **B2MWM9** | Calreticulin OS=Mus musculus OX=10090 GN=Calr PE=1 SV=1 | 8,557 | 2 | 48 | 2,461 |
| **Q3TRW3** | Uncharacterized protein OS=Mus musculus OX=10090 GN=Snd1 PE=2 SV=1 | 2,925 | 2 | 102 | 2,434 |
| **P56399** | Ubiquitin carboxyl-terminal hydrolase 5 OS=Mus musculus OX=10090 GN=Usp5 PE=1 SV=1 | 3,719 | 1 | 95,8 | 2,431 |
| **Q9JM76** | Actin-related protein 2/3 complex subunit 3 OS=Mus musculus OX=10090 GN=Arpc3 PE=1 SV=3 | 13,378 | 3 | 20,5 | 2,426 |
| **P80315** | T-complex protein 1 subunit delta OS=Mus musculus OX=10090 GN=Cct4 PE=1 SV=3 | 13,097 | 2 | 58 | 2,42 |
| **Q102J0** | Chitobiase, di-N-acetyl-, isoform CRA_a OS=Mus musculus OX=10090 GN=Ctbs PE=2 SV=1 | 7,358 | 1 | 41,3 | 2,42 |
| **D0ESZ4** | Elastin microfibril interfacer 2 OS=Mus musculus OX=10090 GN=Emilin2 PE=2 SV=1 | 5,616 | 3 | 117,2 | 2,403 |
| **P50580** | Proliferation-associated protein 2G4 OS=Mus musculus OX=10090 GN=Pa2g4 PE=1 SV=3 | 15,591 | 3 | 43,7 | 2,398 |
| **B2RRX1** | Actin, beta OS=Mus musculus OX=10090 GN=Actb PE=2 SV=1 | 176,64 | 8 | 41,7 | 2,393 |
| **Q11136** | Xaa-Pro dipeptidase OS=Mus musculus OX=10090 GN=Pepd PE=1 SV=3 | 19,536 | 5 | 55 | 2,383 |
| **P62137** | Serine/threonine-protein phosphatase PP1-alpha catalytic subunit OS=Mus musculus OX=10090 GN=Ppp1ca PE=1 SV=1 | 9,278 | 2 | 37,5 | 2,372 |
| **P62259** | 14-3-3 protein epsilon OS=Mus musculus OX=10090 GN=Ywhae PE=1 SV=1 | 32,437 | 4 | 29,2 | 2,354 |
| **Q3TGW0** | Uncharacterized protein OS=Mus musculus OX=10090 GN=Actr3 PE=2 SV=1 | 50,92 | 10 | 47,3 | 2,348 |
| **Q8K183** | Pyridoxal kinase OS=Mus musculus OX=10090 GN=Pdxk PE=1 SV=1 | 13,652 | 3 | 35 | 2,348 |
| **Q3TJ43** | Vacuolar protein sorting-associated protein 35 OS=Mus musculus OX=10090 GN=Vps35 PE=2 SV=1 | 10,577 | 4 | 91,6 | 2,343 |
| **P17751** | Triosephosphate isomerase OS=Mus musculus OX=10090 GN=Tpi1 PE=1 SV=4 | 82,906 | 9 | 32,2 | 2,342 |
| **Q06138** | Calcium-binding protein 39 OS=Mus musculus OX=10090 GN=Cab39 PE=1 SV=2 | 5,41 | 2 | 39,8 | 2,328 |
| **A0A1W2P7A1** | 40S ribosomal protein S12 OS=Mus musculus OX=10090 GN=Rps12 PE=1 SV=1 | 1,159 | 1 | 16 | 2,316 |
| **P62852** | 40S ribosomal protein S25 OS=Mus musculus OX=10090 GN=Rps25 PE=1 SV=1 | 6,809 | 3 | 13,7 | 2,311 |

| | | | | | |
|---|---|---|---|---|---|
| **P09405** | Nucleolin OS=Mus musculus OX=10090 GN=Ncl PE=1 SV=2 | 27,168 | 7 | 76,7 | 2,302 |
| **P50396** | Rab GDP dissociation inhibitor alpha OS=Mus musculus OX=10090 GN=Gdi1 PE=1 SV=3 | 19,642 | 3 | 50,5 | 2,273 |
| **P14685** | 26S proteasome non-ATPase regulatory subunit 3 OS=Mus musculus OX=10090 GN=Psmd3 PE=1 SV=3 | 4,131 | 2 | 60,7 | 2,273 |
| **Q8BTM8** | Filamin-A OS=Mus musculus OX=10090 GN=Flna PE=1 SV=5 | 81,546 | 22 | 281 | 2,268 |
| **O54752** | Naglu OS=Mus musculus OX=10090 GN=Naglu PE=2 SV=1 | 21,866 | 7 | 82,6 | 2,268 |
| **P06745** | Glucose-6-phosphate isomerase OS=Mus musculus OX=10090 GN=Gpi PE=1 SV=4 | 105,28 | 19 | 62,7 | 2,265 |
| **Q61171** | Peroxiredoxin-2 OS=Mus musculus OX=10090 GN=Prdx2 PE=1 SV=3 | 7,958 | 3 | 21,8 | 2,241 |
| **Q58DZ3** | MCG20799 OS=Mus musculus OX=10090 GN=Rpl30 PE=1 SV=1 | 2,885 | 1 | 12,8 | 2,233 |
| **P47757-4** | Isoform 3 of F-actin-capping protein subunit beta OS=Mus musculus OX=10090 GN=Capzb | 7,395 | 4 | 33,7 | 2,226 |
| **P28656** | Nucleosome assembly protein 1-like 1 OS=Mus musculus OX=10090 GN=Nap1l1 PE=1 SV=2 | 4,36 | 2 | 45,3 | 2,223 |
| **Q3TCP5** | Uncharacterized protein OS=Mus musculus OX=10090 GN=Ezr PE=2 SV=1 | 11,035 | 1 | 69,4 | 2,221 |
| **Q3THN2** | Uncharacterized protein OS=Mus musculus OX=10090 GN=Dera PE=2 SV=1 | 2,539 | 2 | 35 | 2,217 |
| **Q9JHU4** | Cytoplasmic dynein 1 heavy chain 1 OS=Mus musculus OX=10090 GN=Dync1h1 PE=1 SV=2 | 13,781 | 3 | 531,7 | 2,214 |
| **P26041** | Moesin OS=Mus musculus OX=10090 GN=Msn PE=1 SV=3 | 63,215 | 8 | 67,7 | 2,197 |
| **Q9QXS1** | Plectin OS=Mus musculus OX=10090 GN=Plec PE=1 SV=3 | 115,783 | 33 | 533,9 | 2,193 |
| **Q8C847** | Beta-galactosidase OS=Mus musculus OX=10090 PE=2 SV=1 | 26,841 | 8 | 84,9 | 2,184 |
| **Q923D2** | Flavin reductase (NADPH) OS=Mus musculus OX=10090 GN=Blvrb PE=1 SV=3 | 19,437 | 5 | 22,2 | 2,184 |
| **Q5CZY9** | Rps16 protein OS=Mus musculus OX=10090 GN=Rps16 PE=2 SV=1 | 2,13 | 1 | 19,3 | 2,181 |
| **Q3UWT6** | Proteasome subunit alpha type OS=Mus musculus OX=10090 GN=Psma2 PE=2 SV=1 | 8,944 | 4 | 27,5 | 2,18 |
| **Q542X7** | Chaperonin subunit 2 (Beta), isoform CRA_a OS=Mus musculus OX=10090 GN=Cct2 PE=1 SV=1 | 30,746 | 6 | 57,4 | 2,175 |
| **P70195** | Proteasome subunit beta type-7 OS=Mus musculus OX=10090 GN=Psmb7 PE=1 SV=1 | 1,201 | 1 | 29,9 | 2,166 |

| | | | | | |
|---|---|---|---|---|---|
| **P47791** | Glutathione reductase, mitochondrial OS=Mus musculus OX=10090 GN=Gsr PE=1 SV=3 | 4,996 | 2 | 53,6 | 2,156 |
| **Q6ZQK2** | MKIAA0051 protein (Fragment) OS=Mus musculus OX=10090 GN=Iqgap1 PE=2 SV=1 | 63,351 | 14 | 191,2 | 2,148 |
| **P29699** | Alpha-2-HS-glycoprotein OS=Mus musculus OX=10090 GN=Ahsg PE=1 SV=1 | 1,428 | 1 | 37,3 | 2,144 |
| **P57722** | Poly(rC)-binding protein 3 OS=Mus musculus OX=10090 GN=Pcbp3 PE=1 SV=3 | 1,04 | 1 | 39,3 | 2,139 |
| **O08800** | Serpin B8 OS=Mus musculus OX=10090 GN=Serpinb8 PE=1 SV=2 | 13,732 | 4 | 42,1 | 2,132 |
| **F8WIV2** | Serine (or cysteine) peptidase inhibitor, clade B, member 6a OS=Mus musculus OX=10090 GN=Serpinb6a PE=1 SV=1 | 27,138 | 6 | 44,7 | 2,126 |
| **P05063** | Fructose-bisphosphate aldolase C OS=Mus musculus OX=10090 GN=Aldoc PE=1 SV=4 | 9,729 | 2 | 39,4 | 2,125 |
| **Q497I3** | Fatty acid binding protein 5, epidermal OS=Mus musculus OX=10090 GN=Fabp5 PE=1 SV=1 | 8,179 | 4 | 15,1 | 2,1 |
| **P12265** | Beta-glucuronidase OS=Mus musculus OX=10090 GN=Gusb PE=1 SV=2 | 75,432 | 13 | 74,1 | 2,095 |
| **P35979** | 60S ribosomal protein L12 OS=Mus musculus OX=10090 GN=Rpl12 PE=1 SV=2 | 5,568 | 2 | 17,8 | 2,087 |
| **D3Z722** | 40S ribosomal protein S19 OS=Mus musculus OX=10090 GN=Rps19 PE=1 SV=1 | 7,394 | 4 | 23,1 | 2,082 |
| **G3X8U3** | Queuosine salvage protein OS=Mus musculus OX=10090 GN=2210016F16Rik PE=1 SV=1 | 2,353 | 1 | 38,6 | 2,059 |
| **Q6ZWQ6** | Huntingtin interacting protein 2, isoform CRA_e OS=Mus musculus OX=10090 GN=Ube2k PE=1 SV=1 | 4,338 | 1 | 22,4 | 2,048 |
| **Q11011** | Puromycin-sensitive aminopeptidase OS=Mus musculus OX=10090 GN=Npepps PE=1 SV=2 | 3,69 | 1 | 103,3 | 2,045 |
| **Q64727** | Vinculin OS=Mus musculus OX=10090 GN=Vcl PE=1 SV=4 | 32,118 | 7 | 116,6 | 2,023 |
| **Q3U6S1** | Uncharacterized protein OS=Mus musculus OX=10090 GN=Vim PE=2 SV=1 | 28,163 | 10 | 53,6 | 2,021 |
| **Q544Y7** | Cofilin 1, non-muscle OS=Mus musculus OX=10090 GN=Cfl1 PE=2 SV=1 | 49,973 | 9 | 18,5 | 2,012 |
| **Q5SX50** | Profilin OS=Mus musculus OX=10090 GN=Pfn1 PE=1 SV=1 | 45,55 | 7 | 14,9 | 2,004 |
| **P70372** | ELAV-like protein 1 OS=Mus musculus OX=10090 GN=Elavl1 PE=1 SV=2 | 1,98 | 1 | 36,1 | 2,004 |
| **F6VW30** | 14-3-3 protein theta (Fragment) OS=Mus musculus OX=10090 GN=Ywhaq PE=1 SV=1 | 11,693 | 1 | 34,3 | 2,003 |

| | | | | | |
|---|---|---|---|---|---|
| **Q9JMA1** | Ubiquitin carboxyl-terminal hydrolase 14 OS=Mus musculus OX=10090 GN=Usp14 PE=1 SV=3 | 15,408 | 5 | 56 | 1,987 |
| **A0A0R4J138** | Arylsulfatase B OS=Mus musculus OX=10090 GN=Arsb PE=1 SV=1 | 7,726 | 4 | 59,7 | 1,96 |
| **A8IP69** | 14-3-3 protein gamma subtype OS=Mus musculus OX=10090 GN=Ywhag PE=1 SV=1 | 56,684 | 6 | 28,3 | 1,959 |
| **A0A2K6EDK8** | Dipeptidase (Fragment) OS=Mus musculus OX=10090 GN=Dpep2 PE=1 SV=1 | 2,413 | 1 | 64,7 | 1,957 |
| **Q9D8N0** | Elongation factor 1-gamma OS=Mus musculus OX=10090 GN=Eef1g PE=1 SV=3 | 18,636 | 4 | 50 | 1,951 |
| **A0A1B0GSX0** | L-lactate dehydrogenase OS=Mus musculus OX=10090 GN=Ldha PE=1 SV=1 | 111,399 | 2 | 39,7 | 1,927 |
| **Q3TI05** | Chaperonin containing Tcp1, subunit 6a (Zeta) OS=Mus musculus OX=10090 GN=Cct6a PE=2 SV=1 | 14,515 | 4 | 58 | 1,922 |
| **Q3UZG4** | Aminoacyl tRNA synthase complex-interacting multifunctional protein 1 OS=Mus musculus OX=10090 GN=Aimp1 PE=1 SV=1 | 3,81 | 1 | 35,1 | 1,91 |
| **O88342** | WD repeat-containing protein 1 OS=Mus musculus OX=10090 GN=Wdr1 PE=1 SV=3 | 49,976 | 11 | 66,4 | 1,907 |
| **Q6GQT1** | Alpha-2-macroglobulin-P OS=Mus musculus OX=10090 GN=A2m PE=2 SV=2 | 24,082 | 2 | 164,2 | 1,891 |
| **Q4FJQ0** | MCG130610 OS=Mus musculus OX=10090 GN=Rab7 PE=1 SV=1 | 11,27 | 3 | 23,5 | 1,882 |
| **Q3UJW9** | Uncharacterized protein OS=Mus musculus OX=10090 GN=Akr1a1 PE=2 SV=1 | 25,361 | 8 | 36,6 | 1,879 |
| **P61089** | Ubiquitin-conjugating enzyme E2 N OS=Mus musculus OX=10090 GN=Ube2n PE=1 SV=1 | 8,223 | 4 | 17,1 | 1,877 |
| **A0A1S6GWH5** | Uncharacterized protein OS=Mus musculus OX=10090 GN=Uba1 PE=2 SV=1 | 36,496 | 8 | 124,2 | 1,868 |
| **Q3UYQ4** | Uncharacterized protein OS=Mus musculus OX=10090 GN=Api5 PE=2 SV=1 | 1,141 | 1 | 59 | 1,865 |
| **P10126** | Elongation factor 1-alpha 1 OS=Mus musculus OX=10090 GN=Eef1a1 PE=1 SV=3 | 62,76 | 10 | 50,1 | 1,859 |
| **Q543N3** | LIM and SH3 protein 1, isoform CRA_b OS=Mus musculus OX=10090 GN=Lasp1 PE=1 SV=1 | 2,807 | 1 | 30 | 1,849 |
| **A1L3T3** | N-sulfoglucosamine sulfohydrolase (Sulfamidase) OS=Mus musculus OX=10090 GN=Sgsh PE=2 SV=1 | 7,807 | 3 | 56,7 | 1,843 |
| **P99024** | Tubulin beta-5 chain OS=Mus musculus OX=10090 GN=Tubb5 PE=1 SV=1 | 39,643 | 2 | 49,6 | 1,842 |
| **Q58E70** | Tpm3 protein OS=Mus musculus OX=10090 GN=Tpm3 PE=2 SV=1 | 13,296 | 6 | 29 | 1,841 |
| **P35700** | Peroxiredoxin-1 OS=Mus musculus OX=10090 GN=Prdx1 PE=1 SV=1 | 31,942 | 10 | 22,2 | 1,832 |
| **Q3TAV1** | Uncharacterized protein OS=Mus musculus OX=10090 GN=Gpnmb PE=2 SV=1 | 30,419 | 3 | 63,6 | 1,83 |

| | | | | | |
|---|---|---|---|---|---|
| **P80313** | T-complex protein 1 subunit eta OS=Mus musculus OX=10090 GN=Cct7 PE=1 SV=1 | 14,499 | 2 | 59,6 | 1,821 |
| **Q80YQ1** | Thrombospondin-1 OS=Mus musculus OX=10090 GN=Thbs1 PE=1 SV=1 | 37,868 | 9 | 129,6 | 1,82 |
| **P51855** | Glutathione synthetase OS=Mus musculus OX=10090 GN=Gss PE=1 SV=1 | 6,142 | 2 | 52,2 | 1,82 |
| **Q9ES94** | Cathepsin Z OS=Mus musculus OX=10090 GN=Ctsz PE=2 SV=1 | 17,137 | 3 | 34,2 | 1,815 |
| **P34884** | Macrophage migration inhibitory factor OS=Mus musculus OX=10090 GN=Mif PE=1 SV=2 | 1,51 | 1 | 12,5 | 1,811 |
| **O35405** | Phospholipase D3 OS=Mus musculus OX=10090 GN=Pld3 PE=1 SV=1 | 4,765 | 3 | 54,4 | 1,81 |
| **Q99PL5** | Ribosome-binding protein 1 OS=Mus musculus OX=10090 GN=Rrbp1 PE=1 SV=2 | 1,994 | 2 | 172,8 | 1,803 |
| **P40142** | Transketolase OS=Mus musculus OX=10090 GN=Tkt PE=1 SV=1 | 71,043 | 12 | 67,6 | 1,8 |
| **Q61768** | Kinesin-1 heavy chain OS=Mus musculus OX=10090 GN=Kif5b PE=1 SV=3 | 2,962 | 2 | 109,5 | 1,797 |
| **Q66JR7** | Pgm2 protein (Fragment) OS=Mus musculus OX=10090 GN=Pgm1 PE=2 SV=1 | 3,538 | 3 | 64,1 | 1,783 |
| **Q9R1P0** | Proteasome subunit alpha type-4 OS=Mus musculus OX=10090 GN=Psma4 PE=1 SV=1 | 31,948 | 5 | 29,5 | 1,776 |
| **P41245** | Matrix metalloproteinase-9 OS=Mus musculus OX=10090 GN=Mmp9 PE=1 SV=2 | 161,402 | 24 | 80,5 | 1,768 |
| **Q5SW83** | ARP2 actin-related protein 2 homolog (Yeast) OS=Mus musculus OX=10090 GN=Actr2 PE=1 SV=1 | 7,964 | 3 | 44,7 | 1,767 |
| **P62858** | 40S ribosomal protein S28 OS=Mus musculus OX=10090 GN=Rps28 PE=1 SV=1 | 5,27 | 1 | 7,8 | 1,764 |
| **P62830** | 60S ribosomal protein L23 OS=Mus musculus OX=10090 GN=Rpl23 PE=1 SV=1 | 2,845 | 1 | 14,9 | 1,753 |
| **Q9QYX7** | Protein piccolo OS=Mus musculus OX=10090 GN=Pclo PE=1 SV=4 | 1,262 | 1 | 550,5 | 1,745 |
| **Q3UBZ3** | Uncharacterized protein OS=Mus musculus OX=10090 GN=Capza2 PE=2 SV=1 | 10,321 | 3 | 33 | 1,727 |
| **Q571M2** | MKIAA4025 protein (Fragment) OS=Mus musculus OX=10090 GN=Hspa4 PE=2 SV=1 | 41,19 | 12 | 103,2 | 1,721 |
| **Q62009** | Periostin OS=Mus musculus OX=10090 GN=Postn PE=1 SV=2 | 2,704 | 1 | 93,1 | 1,717 |
| **Q9CQV8** | 14-3-3 protein beta/alpha OS=Mus musculus OX=10090 GN=Ywhab PE=1 SV=3 | 44,609 | 4 | 28,1 | 1,715 |
| **A6ZI44** | Fructose-bisphosphate aldolase OS=Mus musculus OX=10090 GN=Aldoa PE=1 SV=1 | 91,391 | 13 | 45,1 | 1,71 |
| **P52480-2** | Isoform M1 of Pyruvate kinase PKM OS=Mus musculus OX=10090 GN=Pkm | 140,205 | 1 | 57,9 | 1,707 |
| **Q3TF14** | Adenosylhomocysteinase OS=Mus musculus OX=10090 GN=Ahcy PE=1 SV=1 | 17,905 | 6 | 47,7 | 1,7 |
| **P40240** | CD9 antigen OS=Mus musculus OX=10090 GN=Cd9 PE=1 SV=2 | 7,684 | 1 | 25,2 | 1,7 |

| | | | | | |
|---|---|---|---|---|---|
| **P52480** | Pyruvate kinase PKM OS=Mus musculus OX=10090 GN=Pkm PE=1 SV=4 | 165,673 | 4 | 57,8 | 1,694 |
| **P23492** | Purine nucleoside phosphorylase OS=Mus musculus OX=10090 GN=Pnp PE=1 SV=2 | 13,733 | 4 | 32,3 | 1,691 |
| **Q6A028** | Switch-associated protein 70 OS=Mus musculus OX=10090 GN=Swap70 PE=1 SV=2 | 8,235 | 2 | 69 | 1,691 |
| **G3X977** | Inter-alpha trypsin inhibitor, heavy chain 2 OS=Mus musculus OX=10090 GN=Itih2 PE=1 SV=1 | 30,772 | 6 | 106,3 | 1,687 |
| **P63017** | Heat shock cognate 71 kDa protein OS=Mus musculus OX=10090 GN=Hspa8 PE=1 SV=1 | 145,885 | 21 | 70,8 | 1,664 |
| **P20060** | Beta-hexosaminidase subunit beta OS=Mus musculus OX=10090 GN=Hexb PE=1 SV=2 | 33,321 | 11 | 61,1 | 1,65 |
| **Q3TLP8** | RAS-related C3 botulinum substrate 1, isoform CRA_a OS=Mus musculus OX=10090 GN=Rac1 PE=1 SV=1 | 0,995 | 1 | 23,4 | 1,648 |
| **Q9CQ65** | S-methyl-5'-thioadenosine phosphorylase OS=Mus musculus OX=10090 GN=Mtap PE=1 SV=1 | 5,008 | 1 | 31 | 1,645 |
| **Q8VDD5** | Myosin-9 OS=Mus musculus OX=10090 GN=Myh9 PE=1 SV=4 | 75,365 | 16 | 226,2 | 1,636 |
| **Q3V0Z8** | Uncharacterized protein (Fragment) OS=Mus musculus OX=10090 GN=Ddx5 PE=2 SV=1 | 4,931 | 2 | 76,7 | 1,625 |
| **Q3TS44** | Proteasome subunit alpha type OS=Mus musculus OX=10090 GN=Psma1 PE=1 SV=1 | 18,021 | 6 | 29,5 | 1,573 |
| **Q20BD0** | Heterogeneous nuclear ribonucleoprotein A/B OS=Mus musculus OX=10090 GN=Hnrnpab PE=1 SV=1 | 11,11 | 4 | 36,2 | 1,564 |
| **Q921M7** | Protein FAM49B OS=Mus musculus OX=10090 GN=Fam49b PE=1 SV=1 | 7,214 | 2 | 36,8 | 1,564 |
| **Q3UPA3** | Rab GDP dissociation inhibitor (Fragment) OS=Mus musculus OX=10090 GN=Gdi2 PE=2 SV=1 | 76,78 | 11 | 57,7 | 1,563 |
| **P97807** | Fumarate hydratase, mitochondrial OS=Mus musculus OX=10090 GN=Fh PE=1 SV=3 | 3,304 | 1 | 54,3 | 1,554 |
| **Q3U5L3** | Uncharacterized protein OS=Mus musculus OX=10090 GN=Snx2 PE=2 SV=1 | 1,854 | 1 | 58,4 | 1,531 |
| **Q8BFR4** | N-acetylglucosamine-6-sulfatase OS=Mus musculus OX=10090 GN=Gns PE=1 SV=1 | 22,581 | 9 | 61,1 | 1,514 |

| | | | | | |
|---|---|---|---|---|---|
| **Q60963** | Platelet-activating factor acetylhydrolase OS=Mus musculus OX=10090 GN=Pla2g7 PE=2 SV=2 | 25,324 | 6 | 49,2 | 1,503 |
| **E9QNP0** | KxDL motif-containing protein 1 OS=Mus musculus OX=10090 GN=Kxd1 PE=1 SV=1 | 6,923 | 2 | 26,8 | 1,498 |
| **J3QPG5** | Prosaposin OS=Mus musculus OX=10090 GN=Psap PE=1 SV=1 | 6,748 | 4 | 61,3 | 1,498 |
| **P68040** | Receptor of activated protein C kinase 1 OS=Mus musculus OX=10090 GN=Rack1 PE=1 SV=3 | 17,993 | 7 | 35,1 | 1,487 |
| **A0A0G2JFP4** | Ferric-chelate reductase 1 OS=Mus musculus OX=10090 GN=Frrs1 PE=1 SV=1 | 3,003 | 2 | 66 | 1,473 |
| **Q5FW97** | Enolase 1, alpha non-neuron OS=Mus musculus OX=10090 GN=EG433182 PE=1 SV=1 | 142,771 | 14 | 47,1 | 1,467 |
| **D3YUM8** | Proteasome subunit beta OS=Mus musculus OX=10090 GN=Gm4950 PE=1 SV=1 | 6,764 | 2 | 23 | 1,451 |
| **A7M7S6** | Hemoglobin X, alpha-like embryonic chain in Hba complex (Fragment) OS=Mus musculus OX=10090 GN=Hba-x PE=1 SV=1 | 0,936 | 1 | 18 | 1,447 |
| **Q3TFG3** | Uncharacterized protein OS=Mus musculus OX=10090 GN=Eif4a1 PE=2 SV=1 | 38,53 | 11 | 46,2 | 1,445 |
| **P09411** | Phosphoglycerate kinase 1 OS=Mus musculus OX=10090 GN=Pgk1 PE=1 SV=4 | 88,794 | 13 | 44,5 | 1,444 |
| **O08553** | Dihydropyrimidinase-related protein 2 OS=Mus musculus OX=10090 GN=Dpysl2 PE=1 SV=2 | 14,153 | 4 | 62,2 | 1,444 |
| **P62827** | GTP-binding nuclear protein Ran OS=Mus musculus OX=10090 GN=Ran PE=1 SV=3 | 13,434 | 4 | 24,4 | 1,435 |
| **Q3TN31** | Proteasome subunit alpha type OS=Mus musculus OX=10090 GN=Psma7 PE=2 SV=1 | 27,523 | 7 | 27,9 | 1,415 |
| **Q80U89** | MKIAA0034 protein (Fragment) OS=Mus musculus OX=10090 GN=mKIAA0034 PE=4 SV=2 | 55,767 | 15 | 192,4 | 1,394 |
| **Q542F1** | Chloride intracellular channel protein OS=Mus musculus OX=10090 GN=Clic1 PE=1 SV=1 | 17,028 | 5 | 27 | 1,382 |
| **Q8CHP8** | Glycerol-3-phosphate phosphatase OS=Mus musculus OX=10090 GN=Pgp PE=1 SV=1 | 9,735 | 2 | 34,5 | 1,376 |
| **Q8VCT3** | Aminopeptidase B OS=Mus musculus OX=10090 GN=Rnpep PE=1 SV=2 | 20,421 | 7 | 72,4 | 1,369 |
| **Q3T9S3** | SET translocation OS=Mus musculus OX=10090 GN=Set PE=2 SV=1 | 1,158 | 1 | 33,4 | 1,336 |
| **E9Q616** | AHNAK nucleoprotein (desmoyokin) OS=Mus musculus OX=10090 GN=Ahnak PE=1 SV=1 | 8,338 | 4 | 603,9 | 1,327 |

| | | | | | |
|---|---|---|---|---|---|
| **Q8R1F1** | Niban-like protein 1 OS=Mus musculus OX=10090 GN=Fam129b PE=1 SV=2 | 4,115 | 2 | 84,8 | 1,326 |
| **Q8R016** | Bleomycin hydrolase OS=Mus musculus OX=10090 GN=Blmh PE=1 SV=1 | 6,749 | 2 | 52,5 | 1,314 |
| **Q3UD32** | Uncharacterized protein OS=Mus musculus OX=10090 GN=Ctss PE=2 SV=1 | 51,117 | 9 | 38,8 | 1,302 |
| **Q9CY64** | Biliverdin reductase A OS=Mus musculus OX=10090 GN=Blvra PE=1 SV=1 | 5,367 | 3 | 33,5 | 1,302 |
| **Q3T9L1** | Coronin OS=Mus musculus OX=10090 GN=Coro1a PE=2 SV=1 | 7,255 | 2 | 51 | 1,298 |
| **A8DUN2** | Beta-globin OS=Mus musculus OX=10090 GN=Hbbt1 PE=3 SV=1 | 12,761 | 2 | 15,7 | 1,293 |
| **A0A0A0MQF6** | Glyceraldehyde-3-phosphate dehydrogenase OS=Mus musculus OX=10090 GN=Gapdh PE=1 SV=1 | 23,193 | 7 | 38,6 | 1,292 |
| **Q9ET22** | Dipeptidyl peptidase 2 OS=Mus musculus OX=10090 GN=Dpp7 PE=1 SV=2 | 11,853 | 2 | 56,2 | 1,283 |
| **P68510** | 14-3-3 protein eta OS=Mus musculus OX=10090 GN=Ywhah PE=1 SV=2 | 23,354 | 4 | 28,2 | 1,262 |
| **D3YXG0** | Hemicentin-1 OS=Mus musculus OX=10090 GN=Hmcn1 PE=1 SV=1 | 0,903 | 1 | 611,2 | 1,261 |
| **Q0PD66** | RAB1B, member RAS oncogene family, isoform CRA_c OS=Mus musculus OX=10090 GN=Rab1b PE=1 SV=1 | 4,52 | 2 | 22,2 | 1,26 |
| **A6MDD3** | CD109 antigen OS=Mus musculus OX=10090 GN=Cd109 PE=1 SV=1 | 20,475 | 6 | 161,6 | 1,241 |
| **A0A068BIT8** | Proteasome subunit beta OS=Mus musculus OX=10090 GN=Psmb8 PE=2 SV=1 | 2,236 | 1 | 30,3 | 1,24 |
| **Q4FJY5** | Ltb4dh protein OS=Mus musculus OX=10090 GN=Ptgr1 PE=2 SV=1 | 5,505 | 1 | 35,6 | 1,224 |
| **Q3U4U6** | T-complex protein 1 subunit gamma OS=Mus musculus OX=10090 GN=Cct3 PE=1 SV=1 | 11,768 | 5 | 60,6 | 1,223 |
| **Q8C6A3** | Uncharacterized protein OS=Mus musculus OX=10090 GN=Prep PE=2 SV=1 | 7,249 | 4 | 82,7 | 1,219 |
| **P20029** | Endoplasmic reticulum chaperone BiP OS=Mus musculus OX=10090 GN=Hspa5 PE=1 SV=3 | 28,172 | 5 | 72,4 | 1,215 |
| **P40336-2** | Isoform 2 of Vacuolar protein sorting-associated protein 26A OS=Mus musculus OX=10090 GN=Vps26a | 4,253 | 2 | 41,5 | 1,214 |
| **P61759** | Prefoldin subunit 3 OS=Mus musculus OX=10090 GN=Vbp1 PE=1 SV=2 | 1,85 | 1 | 22,4 | 1,213 |
| **Q9QVP9** | Protein-tyrosine kinase 2-beta OS=Mus musculus OX=10090 GN=Ptk2b PE=1 SV=2 | 1,021 | 1 | 115,7 | 1,202 |
| **Q9CRB5** | Prolactin-7C1 OS=Mus musculus OX=10090 GN=Prl7c1 PE=2 SV=1 | 0,846 | 1 | 29,2 | 1,201 |
| **Q3TLX1** | Uncharacterized protein OS=Mus musculus OX=10090 GN=Nampt PE=2 SV=1 | 7,333 | 3 | 55,4 | 1,183 |
| **E9PYL9** | Predicted gene 10036 OS=Mus musculus OX=10090 GN=Gm10036 PE=3 SV=1 | 1,939 | 1 | 20,3 | 1,172 |
| **Q8BVI4** | Dihydropteridine reductase OS=Mus musculus OX=10090 GN=Qdpr PE=1 SV=2 | 13,872 | 5 | 25,6 | 1,154 |

| | | | | | |
|---|---|---|---|---|---|
| **P43277** | Histone H1.3 OS=Mus musculus OX=10090 GN=Hist1h1d PE=1 SV=2 | 1,353 | 1 | 22,1 | 1,144 |
| **Q9JMH6** | Thioredoxin reductase 1, cytoplasmic OS=Mus musculus OX=10090 GN=Txnrd1 PE=1 SV=3 | 31,741 | 5 | 67 | 1,135 |
| **Q32NZ6** | Transmembrane channel-like protein 5 OS=Mus musculus OX=10090 GN=Tmc5 PE=2 SV=1 | 1,052 | 1 | 111 | 1,12 |
| **Q3UDY1** | MCG6067, isoform CRA_b OS=Mus musculus OX=10090 GN=Akr1b3 PE=1 SV=1 | 6,888 | 2 | 35,7 | 1,113 |
| **Q6A0D0** | MKIAA0106 protein (Fragment) OS=Mus musculus OX=10090 GN=Prdx6 PE=2 SV=1 | 12,75 | 6 | 25,1 | 1,093 |
| **Q99KI0** | Aconitate hydratase, mitochondrial OS=Mus musculus OX=10090 GN=Aco2 PE=1 SV=1 | 25,83 | 7 | 85,4 | 1,083 |
| **Q3UDD5** | Dipeptidyl peptidase 3 OS=Mus musculus OX=10090 GN=Dpp3 PE=2 SV=1 | 37,577 | 9 | 82,9 | 1,07 |
| **P09103** | Protein disulfide-isomerase OS=Mus musculus OX=10090 GN=P4hb PE=1 SV=2 | 14,571 | 5 | 57 | 1,069 |
| **O35235** | Tumor necrosis factor ligand superfamily member 11 OS=Mus musculus OX=10090 GN=Tnfsf11 PE=1 SV=2 | 21,717 | 4 | 35 | 1,046 |
| **P24527** | Leukotriene A-4 hydrolase OS=Mus musculus OX=10090 GN=Lta4h PE=1 SV=4 | 4,15 | 2 | 69 | 1,03 |
| **Q91V28** | 6-phosphogluconate dehydrogenase, decarboxylating OS=Mus musculus OX=10090 GN=Pgd PE=2 SV=1 | 83,847 | 17 | 53,2 | 1,026 |
| **Q61838** | Pregnancy zone protein OS=Mus musculus OX=10090 GN=Pzp PE=1 SV=3 | 0,873 | 1 | 165,7 | 1,02 |
| **Q9D8E6** | 60S ribosomal protein L4 OS=Mus musculus OX=10090 GN=Rpl4 PE=1 SV=3 | 3,757 | 2 | 47,1 | 1,014 |
| **P97298** | Pigment epithelium-derived factor OS=Mus musculus OX=10090 GN=Serpinf1 PE=1 SV=2 | 25,16 | 3 | 46,2 | 0,993 |
| **A0A1B0GR11** | Transaldolase OS=Mus musculus OX=10090 GN=Taldo1 PE=1 SV=1 | 16,84 | 7 | 42,1 | 0,987 |
| **Q99PU7** | Ubiquitin carboxyl-terminal hydrolase BAP1 OS=Mus musculus OX=10090 GN=Bap1 PE=1 SV=1 | 0,878 | 1 | 80,4 | 0,967 |
| **Q3TLY5** | Alpha-galactosidase OS=Mus musculus OX=10090 GN=Gla PE=2 SV=1 | 3,726 | 1 | 47,8 | 0,927 |
| **Q3TUI9** | Proteasome subunit alpha type OS=Mus musculus OX=10090 GN=Psma5 PE=2 SV=1 | 7,254 | 2 | 26,4 | 0,916 |
| **Q3U3C2** | Uncharacterized protein OS=Mus musculus OX=10090 GN=Npc2 PE=2 SV=1 | 6,994 | 2 | 16,5 | 0,876 |
| **P27773** | Protein disulfide-isomerase A3 OS=Mus musculus OX=10090 GN=Pdia3 PE=1 SV=2 | 34,6 | 8 | 56,6 | 0,871 |

| | | | | | |
|---|---|---|---|---|---|
| **Q00623** | Apolipoprotein A-I OS=Mus musculus OX=10090 GN=Apoa1 PE=1 SV=2 | 2,775 | 1 | 30,6 | 0,857 |
| **V9GX81** | Maestro heat-like repeat family member 6 OS=Mus musculus OX=10090 GN=Mroh6 PE=1 SV=1 | 1,662 | 1 | 78,4 | 0,847 |
| **Q9D1A2** | Cytosolic non-specific dipeptidase OS=Mus musculus OX=10090 GN=Cndp2 PE=1 SV=1 | 35,087 | 7 | 52,7 | 0,831 |
| **P08249** | Malate dehydrogenase, mitochondrial OS=Mus musculus OX=10090 GN=Mdh2 PE=1 SV=3 | 50,712 | 12 | 35,6 | 0,811 |
| **E9QNY8** | Sacsin OS=Mus musculus OX=10090 GN=Sacs PE=1 SV=1 | 1,111 | 1 | 520,4 | 0,808 |
| **Q61176** | Arginase-1 OS=Mus musculus OX=10090 GN=Arg1 PE=1 SV=1 | 3,345 | 2 | 34,8 | 0,807 |
| **B8JJN0** | Predicted gene 20547 OS=Mus musculus OX=10090 GN=Gm20547 PE=3 SV=1 | 2,437 | 1 | 142,2 | 0,806 |
| **Q3U9N8** | Inosine-5'-monophosphate dehydrogenase OS=Mus musculus OX=10090 GN=Impdh2 PE=2 SV=1 | 3,31 | 1 | 55,7 | 0,799 |
| **P05202** | Aspartate aminotransferase, mitochondrial OS=Mus musculus OX=10090 GN=Got2 PE=1 SV=1 | 15,913 | 4 | 47,4 | 0,775 |
| **O09159** | Lysosomal alpha-mannosidase OS=Mus musculus OX=10090 GN=Man2b1 PE=1 SV=4 | 5,949 | 2 | 114,6 | 0,751 |
| **Q3TJL8** | Uncharacterized protein OS=Mus musculus OX=10090 GN=Pdia6 PE=2 SV=1 | 4,458 | 2 | 48,6 | 0,742 |
| **A0A1B0GSG5** | Ribonuclease inhibitor OS=Mus musculus OX=10090 GN=Rnh1 PE=1 SV=1 | 10,39 | 2 | 53,9 | 0,736 |
| **Q546G4** | Albumin 1 OS=Mus musculus OX=10090 GN=Alb PE=1 SV=1 | 26,164 | 2 | 68,6 | 0,73 |
| **P56480** | ATP synthase subunit beta, mitochondrial OS=Mus musculus OX=10090 GN=Atp5f1b PE=1 SV=2 | 4,643 | 2 | 56,3 | 0,714 |
| **Q792Z1** | MCG140784 OS=Mus musculus OX=10090 GN=Try10 PE=1 SV=1 | 6,346 | 2 | 26,2 | 0,686 |
| **Q9CZU6** | Citrate synthase, mitochondrial OS=Mus musculus OX=10090 GN=Cs PE=1 SV=1 | 8,968 | 3 | 51,7 | 0,581 |
| **Q3US43** | Annexin OS=Mus musculus OX=10090 GN=Anxa1 PE=2 SV=1 | 21,064 | 3 | 40,3 | 0,56 |
| **Q3UCD3** | Annexin OS=Mus musculus OX=10090 GN=Anxa2 PE=2 SV=1 | 28,602 | 7 | 38,6 | 0,556 |
| **Q545N7** | Creatine kinase, mitochondrial 1, ubiquitous, isoform CRA_a OS=Mus musculus OX=10090 GN=Ckmt1 PE=1 SV=1 | 7,955 | 2 | 47 | 0,544 |
| **F7D5X7** | snRNA-activating protein complex subunit 4 (Fragment) OS=Mus musculus OX=10090 GN=Snapc4 PE=1 SV=1 | 1,099 | 1 | 29,5 | 0,527 |

| | | | | | |
|---|---|---|---|---|---|
| **P07141** | Macrophage colony-stimulating factor 1 OS=Mus musculus OX=10090 GN=Csf1 PE=1 SV=2 | 22,455 | 4 | 60,6 | 0,497 |
| **Q7TSG5-2** | Isoform 2 of SH3 domain-containing protein 21 OS=Mus musculus OX=10090 GN=Sh3d21 | 2,204 | 1 | 73,8 | 0,496 |
| **A2A9M4** | Dedicator of cytokinesis protein 7 OS=Mus musculus OX=10090 GN=Dock7 PE=1 SV=2 | 2,853 | 1 | 241,3 | 0,474 |
| **E9Q557** | Desmoplakin OS=Mus musculus OX=10090 GN=Dsp PE=1 SV=1 | 8,063 | 2 | 332,7 | 0,424 |
| **Q3V2E0** | Uncharacterized protein OS=Mus musculus OX=10090 GN=Try5 PE=2 SV=1 | 5,759 | 1 | 27,1 | 0,392 |
| **Q3U9P7** | Succinyl-CoA:3-ketoacid-coenzyme A transferase OS=Mus musculus OX=10090 GN=Oxct1 PE=2 SV=1 | 6,785 | 1 | 56 | 0,368 |
| **Q60710** | Deoxynucleoside triphosphate triphosphohydrolase SAMHD1 OS=Mus musculus OX=10090 GN=Samhd1 PE=1 SV=3 | 1,961 | 2 | 75,8 | 0,35 |
| **Q9Z1R9** | MCG124046 OS=Mus musculus OX=10090 GN=Prss1 PE=1 SV=1 | 8,703 | 1 | 26,1 | 0,321 |
| **F6T4M4** | Serine/arginine repetitive matrix protein 1 (Fragment) OS=Mus musculus OX=10090 GN=Srrm1 PE=1 SV=1 | 0,954 | 1 | 16,2 | 0,272 |
| **E9Q0B5** | Fc fragment of IgG-binding protein OS=Mus musculus OX=10090 GN=Fcgbp PE=1 SV=1 | 1,529 | 1 | 275 | 0,258 |
| **Q9CPN9** | RIKEN cDNA 2210010C04 gene OS=Mus musculus OX=10090 GN=2210010C04Rik PE=1 SV=1 | 3,736 | 1 | 26,4 | 0,257 |
| **B9EKT8** | Uncharacterized protein OS=Mus musculus OX=10090 GN=Cep162 PE=2 SV=1 | 2,863 | 1 | 160,8 | 0,252 |
| **E9Q0F0** | Keratin 78 OS=Mus musculus OX=10090 GN=Krt78 PE=1 SV=1 | 5,617 | 1 | 112,2 | 0,168 |
| **E9Q1Z0** | Keratin 90 OS=Mus musculus OX=10090 GN=Krt90 PE=1 SV=1 | 10,569 | 1 | 58,2 | 0,134 |
| **Q6NXH9** | Keratin, type II cytoskeletal 73 OS=Mus musculus OX=10090 GN=Krt73 PE=1 SV=1 | 19,272 | 1 | 58,9 | 0,129 |
| **A0A2R8VHP3** | Predicted pseudogene 5478 OS=Mus musculus OX=10090 GN=Gm5478 PE=1 SV=1 | 16,203 | 1 | 57,9 | 0,128 |
| **Q62422** | Osteoclast-stimulating factor 1 OS=Mus musculus OX=10090 GN=Ostf1 PE=1 SV=2 | 4,562 | 1 | 23,8 | 0,115 |
| **Q6IFX2** | Keratin, type I cytoskeletal 42 OS=Mus musculus OX=10090 GN=Krt42 PE=1 SV=1 | 13,436 | 1 | 50,1 | 0,104 |
| **P04104** | Keratin, type II cytoskeletal 1 OS=Mus musculus OX=10090 GN=Krt1 PE=1 SV=4 | 12,545 | 1 | 65,6 | 0,098 |

| | | | | | |
|---|---|---|---|---|---|
| **Q3UV17** | Keratin, type II cytoskeletal 2 oral OS=Mus musculus OX=10090 GN=Krt76 PE=1 SV=1 | 10,065 | 1 | 62,8 | 0,078 |
| **E9PX52** | Arf-GAP with SH3 domain, ANK repeat and PH domain-containing protein 2 OS=Mus musculus OX=10090 GN=Asap2 PE=1 SV=1 | 1,474 | 1 | 111,1 | 0,072 |
| **B2RTP7** | Krt2 protein OS=Mus musculus OX=10090 GN=Krt2 PE=2 SV=1 | 10,093 | 2 | 70,9 | 0,06 |
| **Q8BGZ7** | Keratin, type II cytoskeletal 75 OS=Mus musculus OX=10090 GN=Krt75 PE=1 SV=1 | 26,198 | 1 | 59,7 | 0,054 |
| **P02535** | Keratin, type I cytoskeletal 10 OS=Mus musculus OX=10090 GN=Krt10 PE=1 SV=3 | 26,365 | 5 | 57,7 | 0,043 |
| **Q9QWL7** | Keratin, type I cytoskeletal 17 OS=Mus musculus OX=10090 GN=Krt17 PE=1 SV=3 | 23,94 | 5 | 48,1 | 0,039 |
| **Q32P04** | Keratin 5 OS=Mus musculus OX=10090 GN=Krt5 PE=1 SV=2 | 28 | 3 | 61,7 | 0,025 |
| **Q8VED5** | Keratin, type II cytoskeletal 79 OS=Mus musculus OX=10090 GN=Krt79 PE=1 SV=2 | 8,325 | 1 | 57,5 | 0,019 |
| **P50446** | Keratin, type II cytoskeletal 6A OS=Mus musculus OX=10090 GN=Krt6a PE=1 SV=3 | 34,647 | 3 | 59,3 | 0,016 |
| **Q6IME9** | Keratin, type II cytoskeletal 72 OS=Mus musculus OX=10090 GN=Krt72 PE=3 SV=1 | 4,56 | 1 | 56,7 | 0,01 |
| **Q0VBK2** | Keratin, type II cytoskeletal 80 OS=Mus musculus OX=10090 GN=Krt80 PE=1 SV=1 | 0,84 | 1 | 50,6 | 0,01 |
