## Supplemental table S3 for "Sensory neurons sprouting is dependent on osteoclast-derived extracellular vesicles involving the activation of epidermal growth factor receptors"

Table S3: LC-MS/MS analysis of protein content from extracellular vesicles enriched samples from osteoclasts secretome.

| ACCESSION | DESCRIPTION | # PEPTIDES | # UNIQUE PEPTIDES |
|---|---|---|---|
| **P26039** | Talin-1 OS=Mus musculus OX=10090 GN=Tln1 PE=1 SV=2 | 47 | 47 |
| **B7FAV1** | Filamin, alpha (Fragment) OS=Mus musculus OX=10090 GN=Flna PE=1 SV=1 | 31 | 31 |
| **Q8VDD5** | Myosin-9 OS=Mus musculus OX=10090 GN=Myh9 PE=1 SV=4 | 25 | 23 |
| **Q9EQK5** | Major vault protein OS=Mus musculus OX=10090 GN=Mvp PE=1 SV=4 | 20 | 20 |
| **Q68FD5** | Clathrin heavy chain 1 OS=Mus musculus OX=10090 GN=Cltc PE=1 SV=3 | 19 | 19 |
| **P52480** | Pyruvate kinase PKM OS=Mus musculus OX=10090 GN=Pkm PE=1 SV=4 | 16 | 16 |
| **P48036** | Annexin A5 OS=Mus musculus OX=10090 GN=Anxa5 PE=1 SV=1 | 14 | 14 |
| **P17182** | Alpha-enolase OS=Mus musculus OX=10090 GN=Eno1 PE=1 SV=3 | 14 | 14 |
| **P97429** | Annexin A4 OS=Mus musculus OX=10090 GN=Anxa4 PE=1 SV=4 | 12 | 12 |
| **A0A0A6YWC8** | Vimentin OS=Mus musculus OX=10090 GN=Vim PE=1 SV=1 | 12 | 12 |
| **P13020** | Gelsolin OS=Mus musculus OX=10090 GN=Gsn PE=1 SV=3 | 11 | 11 |
| **P07901** | Heat shock protein HSP 90-alpha OS=Mus musculus OX=10090 GN=Hsp90aa1 PE=1 SV=4 | 16 | 11 |
| **P07356** | Annexin A2 OS=Mus musculus OX=10090 GN=Anxa2 PE=1 SV=2 | 11 | 11 |
| **A6ZI44** | Fructose-bisphosphate aldolase OS=Mus musculus OX=10090 GN=Aldoa PE=1 SV=1 | 12 | 11 |
| **Q8K1B8** | Fermitin family homolog 3 OS=Mus musculus OX=10090 GN=Fermt3 PE=1 SV=1 | 11 | 11 |
| **Q9D8E6** | 60S ribosomal protein L4 OS=Mus musculus OX=10090 GN=Rpl4 PE=1 SV=3 | 10 | 10 |
| **P41245** | Matrix metalloproteinase-9 OS=Mus musculus OX=10090 GN=Mmp9 PE=1 SV=2 | 10 | 10 |
| **Q9JKF1** | Ras GTPase-activating-like protein IQGAP1 OS=Mus musculus OX=10090 GN=Iqgap1 PE=1 SV=2 | 10 | 10 |
| **P62242** | 40S ribosomal protein S8 OS=Mus musculus OX=10090 GN=Rps8 PE=1 SV=2 | 9 | 9 |
| **P10107** | Annexin A1 OS=Mus musculus OX=10090 GN=Anxa1 PE=1 SV=2 | 9 | 9 |
| **P09405** | Nucleolin OS=Mus musculus OX=10090 GN=Ncl PE=1 SV=2 | 9 | 9 |
| **P63017** | Heat shock cognate 71 kDa protein OS=Mus musculus OX=10090 GN=Hspa8 PE=1 SV=1 | 12 | 9 |
| **Q61233** | Plastin-2 OS=Mus musculus OX=10090 GN=Lcp1 PE=1 SV=4 | 12 | 9 |
| **Q8VDN2** | Sodium/potassium-transporting ATPase subunit alpha-1 OS=Mus musculus OX=10090 GN=Atp1a1 PE=1 SV=1 | 9 | 9 |
| **P62814** | V-type proton ATPase subunit B, brain isoform OS=Mus musculus OX=10090 GN=Atp6v1b2 PE=1 SV=1 | 9 | 9 |

| | | | |
|---|---|---|---|
| **P58252** | Elongation factor 2 OS=Mus musculus OX=10090 GN=Eef2 PE=1 SV=2 | 9 | 9 |
| **P01027** | Complement C3 OS=Mus musculus OX=10090 GN=C3 PE=1 SV=3 | 8 | 8 |
| **P12970** | 60S ribosomal protein L7a OS=Mus musculus OX=10090 GN=Rpl7a PE=1 SV=2 | 8 | 8 |
| **P35441** | Thrombospondin-1 OS=Mus musculus OX=10090 GN=Thbs1 PE=1 SV=1 | 8 | 8 |
| **P50247** | Adenosylhomocysteinase OS=Mus musculus OX=10090 GN=Ahcy PE=1 SV=3 | 8 | 8 |
| **P62908** | 40S ribosomal protein S3 OS=Mus musculus OX=10090 GN=Rps3 PE=1 SV=1 | 8 | 8 |
| **P14869** | 60S acidic ribosomal protein P0 OS=Mus musculus OX=10090 GN=Rplp0 PE=1 SV=3 | 8 | 8 |
| **P97449** | Aminopeptidase N OS=Mus musculus OX=10090 GN=Anpep PE=1 SV=4 | 8 | 8 |
| **E9Q3W4** | Plectin OS=Mus musculus OX=10090 GN=Plec PE=1 SV=1 | 8 | 8 |
| **P60710** | Actin, cytoplasmic 1 OS=Mus musculus OX=10090 GN=Actb PE=1 SV=1 | 14 | 7 |
| **P32261** | Antithrombin-III OS=Mus musculus OX=10090 GN=Serpinc1 PE=1 SV=1 | 7 | 7 |
| **P14148** | 60S ribosomal protein L7 OS=Mus musculus OX=10090 GN=Rpl7 PE=1 SV=2 | 7 | 7 |
| **P47911** | 60S ribosomal protein L6 OS=Mus musculus OX=10090 GN=Rpl6 PE=1 SV=3 | 7 | 7 |
| **P11499** | Heat shock protein HSP 90-beta OS=Mus musculus OX=10090 GN=Hsp90ab1 PE=1 SV=3 | 12 | 7 |
| **A0A1D5RLW5** | 60S ribosomal protein L18a OS=Mus musculus OX=10090 GN=Rpl18a PE=1 SV=1 | 7 | 7 |
| **Q9JHU4** | Cytoplasmic dynein 1 heavy chain 1 OS=Mus musculus OX=10090 GN=Dync1h1 PE=1 SV=2 | 7 | 7 |
| **Q61703** | Inter-alpha-trypsin inhibitor heavy chain H2 OS=Mus musculus OX=10090 GN=Itih2 PE=1 SV=1 | 6 | 6 |
| **A0A0A0MQA5** | Tubulin alpha chain (Fragment) OS=Mus musculus OX=10090 GN=Tuba4a PE=1 SV=1 | 15 | 6 |
| **P99024** | Tubulin beta-5 chain OS=Mus musculus OX=10090 GN=Tubb5 PE=1 SV=1 | 17 | 6 |
| **P68369** | Tubulin alpha-1A chain OS=Mus musculus OX=10090 GN=Tuba1a PE=1 SV=1 | 15 | 6 |
| **P09411** | Phosphoglycerate kinase 1 OS=Mus musculus OX=10090 GN=Pgk1 PE=1 SV=4 | 6 | 6 |
| **Q04447** | Creatine kinase B-type OS=Mus musculus OX=10090 GN=Ckb PE=1 SV=1 | 6 | 6 |
| **P63101** | 14-3-3 protein zeta/delta OS=Mus musculus OX=10090 GN=Ywhaz PE=1 SV=1 | 8 | 6 |
| **P27659** | 60S ribosomal protein L3 OS=Mus musculus OX=10090 GN=Rpl3 PE=1 SV=3 | 6 | 6 |
| **A0A1B0GSR9** | L-lactate dehydrogenase OS=Mus musculus OX=10090 GN=Ldha PE=1 SV=1 | 7 | 6 |
| **P10126** | Elongation factor 1-alpha 1 OS=Mus musculus OX=10090 GN=Eef1a1 PE=1 SV=3 | 6 | 6 |
| **Q9DBJ1** | Phosphoglycerate mutase 1 OS=Mus musculus OX=10090 GN=Pgam1 PE=1 SV=3 | 6 | 6 |
| **H3BL49** | T-complex protein 1 subunit theta OS=Mus musculus OX=10090 GN=Cct8 PE=1 SV=1 | 6 | 6 |
| **E9PWQ3** | Collagen, type VI, alpha 3 OS=Mus musculus OX=10090 GN=Col6a3 PE=1 SV=2 | 6 | 6 |

| | | | |
|---|---|---|---|
| **Q64727** | Vinculin OS=Mus musculus OX=10090 GN=Vcl PE=1 SV=4 | 6 | 6 |
| **A0A087WS56** | Fibronectin OS=Mus musculus OX=10090 GN=Fn1 PE=1 SV=1 | 6 | 6 |
| **Q8R1B4** | Eukaryotic translation initiation factor 3 subunit C OS=Mus musculus OX=10090 GN=Eif3c PE=1 SV=1 | 6 | 6 |
| **Q7TPV4** | Myb-binding protein 1A OS=Mus musculus OX=10090 GN=Mybbp1a PE=1 SV=2 | 6 | 6 |
| **P62806** | Histone H4 OS=Mus musculus OX=10090 GN=Hist1h4a PE=1 SV=2 | 5 | 5 |
| **H7BX99** | Prothrombin OS=Mus musculus OX=10090 GN=F2 PE=1 SV=1 | 5 | 5 |
| **A0A2I3BRQ3** | Inter-alpha-trypsin inhibitor heavy chain H3 OS=Mus musculus OX=10090 GN=Itih3 PE=1 SV=1 | 5 | 5 |
| **A0A0A0MQF6** | Glyceraldehyde-3-phosphate dehydrogenase OS=Mus musculus OX=10090 GN=Gapdh PE=1 SV=1 | 5 | 5 |
| **A0A1B0GQU8** | 60S ribosomal protein L18 OS=Mus musculus OX=10090 GN=Rpl18 PE=1 SV=1 | 5 | 5 |
| **P01029** | Complement C4-B OS=Mus musculus OX=10090 GN=C4b PE=1 SV=3 | 5 | 5 |
| **F7CJS8** | 40S ribosomal protein S9 (Fragment) OS=Mus musculus OX=10090 GN=Rps9 PE=1 SV=1 | 5 | 5 |
| **Q61696** | Heat shock 70 kDa protein 1A OS=Mus musculus OX=10090 GN=Hspa1a PE=1 SV=2 | 9 | 5 |
| **Q9CPR4** | 60S ribosomal protein L17 OS=Mus musculus OX=10090 GN=Rpl17 PE=1 SV=3 | 5 | 5 |
| **Q99JI6** | Ras-related protein Rap-1b OS=Mus musculus OX=10090 GN=Rap1b PE=1 SV=2 | 5 | 5 |
| **P62259** | 14-3-3 protein epsilon OS=Mus musculus OX=10090 GN=Ywhae PE=1 SV=1 | 7 | 5 |
| **Q61598** | Rab GDP dissociation inhibitor beta OS=Mus musculus OX=10090 GN=Gdi2 PE=1 SV=1 | 6 | 5 |
| **Q8VEK3** | Heterogeneous nuclear ribonucleoprotein U OS=Mus musculus OX=10090 GN=Hnrnpu PE=1 SV=1 | 5 | 5 |
| **Q07113** | Cation-independent mannose-6-phosphate receptor OS=Mus musculus OX=10090 GN=Igf2r PE=1 SV=1 | 5 | 5 |
| **P97351** | 40S ribosomal protein S3a OS=Mus musculus OX=10090 GN=Rps3a PE=1 SV=3 | 5 | 5 |
| **Q9WU78** | Programmed cell death 6-interacting protein OS=Mus musculus OX=10090 GN=Pdcd6ip PE=1 SV=3 | 5 | 5 |
| **P20029** | Endoplasmic reticulum chaperone BiP OS=Mus musculus OX=10090 GN=Hspa5 PE=1 SV=3 | 7 | 5 |
| **P06745** | Glucose-6-phosphate isomerase OS=Mus musculus OX=10090 GN=Gpi PE=1 SV=4 | 5 | 5 |
| **P28271** | Cytoplasmic aconitate hydratase OS=Mus musculus OX=10090 GN=Aco1 PE=1 SV=3 | 5 | 5 |
| **Q05117** | Tartrate-resistant acid phosphatase type 5 OS=Mus musculus OX=10090 GN=Acp5 PE=1 SV=2 | 5 | 5 |
| **P50516** | V-type proton ATPase catalytic subunit A OS=Mus musculus OX=10090 GN=Atp6v1a PE=1 SV=2 | 5 | 5 |
| **Q9R0P3** | S-formylglutathione hydrolase OS=Mus musculus OX=10090 GN=Esd PE=1 SV=1 | 5 | 5 |
| **P97321** | Prolyl endopeptidase FAP OS=Mus musculus OX=10090 GN=Fap PE=1 SV=1 | 5 | 5 |
| **P48678** | Prelamin-A/C OS=Mus musculus OX=10090 GN=Lmna PE=1 SV=2 | 5 | 5 |
| **P11983** | T-complex protein 1 subunit alpha OS=Mus musculus OX=10090 GN=Tcp1 PE=1 SV=3 | 5 | 5 |

| | | | |
|---|---|---|---|
| **A2AKI5** | Integrin alpha-V OS=Mus musculus OX=10090 GN=Itgav PE=1 SV=1 | 5 | 5 |
| **Q01853** | Transitional endoplasmic reticulum ATPase OS=Mus musculus OX=10090 GN=Vcp PE=1 SV=4 | 5 | 5 |
| **Q91VB8** | Alpha globin 1 OS=Mus musculus OX=10090 GN=Hba-a1 PE=1 SV=1 | 4 | 4 |
| **P97298** | Pigment epithelium-derived factor OS=Mus musculus OX=10090 GN=Serpinf1 PE=1 SV=2 | 4 | 4 |
| **P51885** | Lumican OS=Mus musculus OX=10090 GN=Lum PE=1 SV=2 | 4 | 4 |
| **A2AQ07** | Tubulin beta-1 chain OS=Mus musculus OX=10090 GN=Tubb1 PE=1 SV=1 | 7 | 4 |
| **P14115** | 60S ribosomal protein L27a OS=Mus musculus OX=10090 GN=Rpl27a PE=1 SV=5 | 4 | 4 |
| **P14131** | 40S ribosomal protein S16 OS=Mus musculus OX=10090 GN=Rps16 PE=1 SV=4 | 4 | 4 |
| **Q9R0G6** | Cartilage oligomeric matrix protein OS=Mus musculus OX=10090 GN=Comp PE=1 SV=2 | 6 | 4 |
| **P17742** | Peptidyl-prolyl cis-trans isomerase A OS=Mus musculus OX=10090 GN=Ppia PE=1 SV=2 | 4 | 4 |
| **Q5SX49** | Profilin OS=Mus musculus OX=10090 GN=Pfn1 PE=1 SV=1 | 4 | 4 |
| **P11087** | Collagen alpha-1(I) chain OS=Mus musculus OX=10090 GN=Col1a1 PE=1 SV=4 | 4 | 4 |
| **E9Q1Y3** | Apolipoprotein B-100 (Fragment) OS=Mus musculus OX=10090 GN=Apob PE=1 SV=1 | 4 | 4 |
| **P61358** | 60S ribosomal protein L27 OS=Mus musculus OX=10090 GN=Rpl27 PE=1 SV=2 | 4 | 4 |
| **Q9ES46** | Beta-parvin OS=Mus musculus OX=10090 GN=Parvb PE=1 SV=1 | 4 | 4 |
| **I7HLV2** | 60S ribosomal protein L10 (Fragment) OS=Mus musculus OX=10090 GN=Rpl10 PE=1 SV=1 | 4 | 4 |
| **Q9Z204** | Heterogeneous nuclear ribonucleoproteins C1/C2 OS=Mus musculus OX=10090 GN=Hnrnpc PE=1 SV=1 | 4 | 4 |
| **P18760** | Cofilin-1 OS=Mus musculus OX=10090 GN=Cfl1 PE=1 SV=3 | 4 | 4 |
| **P40124** | Adenylyl cyclase-associated protein 1 OS=Mus musculus OX=10090 GN=Cap1 PE=1 SV=4 | 4 | 4 |
| **Q9Z1Q5** | Chloride intracellular channel protein 1 OS=Mus musculus OX=10090 GN=Clic1 PE=1 SV=3 | 4 | 4 |
| **P63158** | High mobility group protein B1 OS=Mus musculus OX=10090 GN=Hmgb1 PE=1 SV=2 | 5 | 4 |
| **Q62009** | Periostin OS=Mus musculus OX=10090 GN=Postn PE=1 SV=2 | 4 | 4 |
| **P16110** | Galectin-3 OS=Mus musculus OX=10090 GN=Lgals3 PE=1 SV=3 | 4 | 4 |
| **Q9WU81** | Glucose-6-phosphate exchanger SLC37A2 OS=Mus musculus OX=10090 GN=Slc37a2 PE=1 SV=1 | 4 | 4 |
| **P97370** | Sodium/potassium-transporting ATPase subunit beta-3 OS=Mus musculus OX=10090 GN=Atp1b3 PE=1 SV=1 | 4 | 4 |
| **P80314** | T-complex protein 1 subunit beta OS=Mus musculus OX=10090 GN=Cct2 PE=1 SV=4 | 4 | 4 |
| **A0A0R4J0I9** | Low density lipoprotein receptor-related protein 1 OS=Mus musculus OX=10090 GN=Lrp1 PE=1 SV=1 | 4 | 4 |
| **P97384** | Annexin A11 OS=Mus musculus OX=10090 GN=Anxa11 PE=1 SV=2 | 4 | 4 |

| | | | |
|---|---|---|---|
| **P60843** | Eukaryotic initiation factor 4A-I OS=Mus musculus OX=10090 GN=Eif4a1 PE=1 SV=1 | 4 | 4 |
| **Q9QZQ8** | Core histone macro-H2A.1 OS=Mus musculus OX=10090 GN=H2afy PE=1 SV=3 | 4 | 4 |
| **P80317** | T-complex protein 1 subunit zeta OS=Mus musculus OX=10090 GN=Cct6a PE=1 SV=3 | 4 | 4 |
| **Q99PV0** | Pre-mRNA-processing-splicing factor 8 OS=Mus musculus OX=10090 GN=Prpf8 PE=1 SV=2 | 4 | 4 |
| **Q99P72** | Reticulon-4 OS=Mus musculus OX=10090 GN=Rtn4 PE=1 SV=2 | 4 | 4 |
| **Q542I8** | Integrin beta OS=Mus musculus OX=10090 GN=Itgb2 PE=1 SV=1 | 4 | 4 |
| **P80316** | T-complex protein 1 subunit epsilon OS=Mus musculus OX=10090 GN=Cct5 PE=1 SV=1 | 4 | 4 |
| **E9Q634** | Unconventional myosin-Ie OS=Mus musculus OX=10090 GN=Myo1e PE=1 SV=1 | 4 | 4 |
| **Q02053** | Ubiquitin-like modifier-activating enzyme 1 OS=Mus musculus OX=10090 GN=Uba1 PE=1 SV=1 | 4 | 4 |
| **Q8BVE3** | V-type proton ATPase subunit H OS=Mus musculus OX=10090 GN=Atp6v1h PE=1 SV=1 | 4 | 4 |
| **P09103** | Protein disulfide-isomerase OS=Mus musculus OX=10090 GN=P4hb PE=1 SV=2 | 4 | 4 |
| **E9Q390** | Myoferlin OS=Mus musculus OX=10090 GN=Myof PE=1 SV=2 | 4 | 4 |
| **F6QYF8** | Aminopeptidase (Fragment) OS=Mus musculus OX=10090 GN=Npepps PE=1 SV=1 | 4 | 4 |
| **P07724** | Serum albumin OS=Mus musculus OX=10090 GN=Alb PE=1 SV=3 | 3 | 3 |
| **Q64475** | Histone H2B type 1-B OS=Mus musculus OX=10090 GN=Hist1h2bb PE=1 SV=3 | 3 | 3 |
| **Q08879** | Fibulin-1 OS=Mus musculus OX=10090 GN=Fbln1 PE=1 SV=2 | 3 | 3 |
| **P25444** | 40S ribosomal protein S2 OS=Mus musculus OX=10090 GN=Rps2 PE=1 SV=3 | 3 | 3 |
| **A0A1B0GS68** | Predicted gene 45713 OS=Mus musculus OX=10090 GN=Gm45713 PE=2 SV=1 | 3 | 3 |
| **Q9CR57** | 60S ribosomal protein L14 OS=Mus musculus OX=10090 GN=Rpl14 PE=1 SV=3 | 3 | 3 |
| **A0A1L1SV25** | Alpha-actinin-4 OS=Mus musculus OX=10090 GN=Actn4 PE=1 SV=1 | 9 | 3 |
| **P43276** | Histone H1.5 OS=Mus musculus OX=10090 GN=Hist1h1b PE=1 SV=2 | 3 | 3 |
| **Q8BP67** | 60S ribosomal protein L24 OS=Mus musculus OX=10090 GN=Rpl24 PE=1 SV=2 | 3 | 3 |
| **O88783** | Coagulation factor V OS=Mus musculus OX=10090 GN=F5 PE=1 SV=1 | 3 | 3 |
| **P62889** | 60S ribosomal protein L30 OS=Mus musculus OX=10090 GN=Rpl30 PE=1 SV=2 | 3 | 3 |
| **Q04857** | Collagen alpha-1(VI) chain OS=Mus musculus OX=10090 GN=Col6a1 PE=1 SV=1 | 3 | 3 |
| **P55097** | Cathepsin K OS=Mus musculus OX=10090 GN=Ctsk PE=1 SV=2 | 3 | 3 |
| **P14206** | 40S ribosomal protein SA OS=Mus musculus OX=10090 GN=Rpsa PE=1 SV=4 | 3 | 3 |
| **P84099** | 60S ribosomal protein L19 OS=Mus musculus OX=10090 GN=Rpl19 PE=1 SV=1 | 3 | 3 |
| **H7BXC3** | Triosephosphate isomerase OS=Mus musculus OX=10090 GN=Tpi1 PE=1 SV=1 | 3 | 3 |

| | | | |
|---|---|---|---|
| **P62918** | 60S ribosomal protein L8 OS=Mus musculus OX=10090 GN=Rpl8 PE=1 SV=2 | 3 | 3 |
| **O88342** | WD repeat-containing protein 1 OS=Mus musculus OX=10090 GN=Wdr1 PE=1 SV=3 | 3 | 3 |
| **P26041** | Moesin OS=Mus musculus OX=10090 GN=Msn PE=1 SV=3 | 4 | 3 |
| **Q9WVK4** | EH domain-containing protein 1 OS=Mus musculus OX=10090 GN=Ehd1 PE=1 SV=1 | 4 | 3 |
| **P70670** | Nascent polypeptide-associated complex subunit alpha, muscle-specific form OS=Mus musculus OX=10090 GN=Naca PE=1 SV=2 | 3 | 3 |
| **P35700** | Peroxiredoxin-1 OS=Mus musculus OX=10090 GN=Prdx1 PE=1 SV=1 | 3 | 3 |
| **P61982** | 14-3-3 protein gamma OS=Mus musculus OX=10090 GN=Ywhag PE=1 SV=2 | 5 | 3 |
| **P62855** | 40S ribosomal protein S26 OS=Mus musculus OX=10090 GN=Rps26 PE=1 SV=3 | 3 | 3 |
| **H7BX95** | Serine/arginine-rich-splicing factor 1 OS=Mus musculus OX=10090 GN=Srsf1 PE=1 SV=1 | 3 | 3 |
| **P62827** | GTP-binding nuclear protein Ran OS=Mus musculus OX=10090 GN=Ran PE=1 SV=3 | 3 | 3 |
| **Q60737** | Casein kinase II subunit alpha OS=Mus musculus OX=10090 GN=Csnk2a1 PE=1 SV=2 | 3 | 3 |
| **P80315** | T-complex protein 1 subunit delta OS=Mus musculus OX=10090 GN=Cct4 PE=1 SV=3 | 3 | 3 |
| **Q01730** | Ras suppressor protein 1 OS=Mus musculus OX=10090 GN=Rsu1 PE=1 SV=3 | 3 | 3 |
| **Q9JHF5** | V-type proton ATPase subunit a OS=Mus musculus OX=10090 GN=Tcirg1 PE=1 SV=1 | 3 | 3 |
| **O08992** | Syntenin-1 OS=Mus musculus OX=10090 GN=Sdcbp PE=1 SV=1 | 3 | 3 |
| **P14211** | Calreticulin OS=Mus musculus OX=10090 GN=Calr PE=1 SV=1 | 3 | 3 |
| **P67984** | 60S ribosomal protein L22 OS=Mus musculus OX=10090 GN=Rpl22 PE=1 SV=2 | 3 | 3 |
| **P62874** | Guanine nucleotide-binding protein G(I)/G(S)/G(T) subunit beta-1 OS=Mus musculus OX=10090 GN=Gnb1 PE=1 SV=3 | 4 | 3 |
| **P29341** | Polyadenylate-binding protein 1 OS=Mus musculus OX=10090 GN=Pabpc1 PE=1 SV=2 | 3 | 3 |
| **Q9Z1Z0** | General vesicular transport factor p115 OS=Mus musculus OX=10090 GN=Uso1 PE=1 SV=2 | 3 | 3 |
| **P23116** | Eukaryotic translation initiation factor 3 subunit A OS=Mus musculus OX=10090 GN=Eif3a PE=1 SV=5 | 3 | 3 |
| **A1BN54** | Alpha actinin 1a OS=Mus musculus OX=10090 GN=Actn1 PE=1 SV=1 | 9 | 3 |
| **P60766** | Cell division control protein 42 homolog OS=Mus musculus OX=10090 GN=Cdc42 PE=1 SV=2 | 3 | 3 |
| **Q8CIE6** | Coatomer subunit alpha OS=Mus musculus OX=10090 GN=Copa PE=1 SV=2 | 3 | 3 |
| **E9Q740** | Signal recognition particle subunit SRP72 OS=Mus musculus OX=10090 GN=Srp72 PE=1 SV=1 | 3 | 3 |
| **Q93092** | Transaldolase OS=Mus musculus OX=10090 GN=Taldo1 PE=1 SV=2 | 3 | 3 |
| **P27773** | Protein disulfide-isomerase A3 OS=Mus musculus OX=10090 GN=Pdia3 PE=1 SV=2 | 3 | 3 |

| | | | |
|---|---|---|---|
| **Q9Z1G3** | V-type proton ATPase subunit C 1 OS=Mus musculus OX=10090 GN=Atp6v1c1 PE=1 SV=4 | 3 | 3 |
| **Q3V117** | ATP-citrate synthase OS=Mus musculus OX=10090 GN=Acly PE=1 SV=1 | 3 | 3 |
| **O88685** | 26S proteasome regulatory subunit 6A OS=Mus musculus OX=10090 GN=Psmc3 PE=1 SV=2 | 3 | 3 |
| **P68510** | 14-3-3 protein eta OS=Mus musculus OX=10090 GN=Ywhah PE=1 SV=2 | 5 | 3 |
| **P08752** | Guanine nucleotide-binding protein G(i) subunit alpha-2 OS=Mus musculus OX=10090 GN=Gnai2 PE=1 SV=5 | 3 | 3 |
| **Q6GQT1** | Alpha-2-macroglobulin-P OS=Mus musculus OX=10090 GN=A2m PE=2 SV=2 | 2 | 2 |
| **P04186** | Complement factor B OS=Mus musculus OX=10090 GN=Cfb PE=1 SV=2 | 2 | 2 |
| **Q61838** | Pregnancy zone protein OS=Mus musculus OX=10090 GN=Pzp PE=1 SV=3 | 2 | 2 |
| **E9Q5F6** | Polyubiquitin-C (Fragment) OS=Mus musculus OX=10090 GN=Ubc PE=1 SV=1 | 2 | 2 |
| **P62754** | 40S ribosomal protein S6 OS=Mus musculus OX=10090 GN=Rps6 PE=1 SV=1 | 2 | 2 |
| **A0A1B0GR60** | Ferritin OS=Mus musculus OX=10090 GN=Ftl1 PE=1 SV=1 | 2 | 2 |
| **Q9JJN5** | Carboxypeptidase N catalytic chain OS=Mus musculus OX=10090 GN=Cpn1 PE=1 SV=1 | 2 | 2 |
| **P61205** | ADP-ribosylation factor 3 OS=Mus musculus OX=10090 GN=Arf3 PE=2 SV=2 | 2 | 2 |
| **Q9Z1R3** | Apolipoprotein M OS=Mus musculus OX=10090 GN=Apom PE=1 SV=1 | 2 | 2 |
| **Q9D7B2** | LIM and senescent cell antigen-like-containing domain protein OS=Mus musculus OX=10090 GN=Lims1 PE=1 SV=2 | 2 | 2 |
| **P00687** | Alpha-amylase 1 OS=Mus musculus OX=10090 GN=Amy1 PE=1 SV=2 | 2 | 2 |
| **P99027** | 60S acidic ribosomal protein P2 OS=Mus musculus OX=10090 GN=Rplp2 PE=1 SV=3 | 2 | 2 |
| **P62267** | 40S ribosomal protein S23 OS=Mus musculus OX=10090 GN=Rps23 PE=1 SV=3 | 2 | 2 |
| **Q9CZM2** | 60S ribosomal protein L15 OS=Mus musculus OX=10090 GN=Rpl15 PE=1 SV=4 | 2 | 2 |
| **O89053** | Coronin-1A OS=Mus musculus OX=10090 GN=Coro1a PE=1 SV=5 | 2 | 2 |
| **Q6GT24** | Peroxiredoxin-6 OS=Mus musculus OX=10090 GN=Prdx6 PE=1 SV=1 | 2 | 2 |
| **P50543** | Protein S100-A11 OS=Mus musculus OX=10090 GN=S100a11 PE=1 SV=1 | 2 | 2 |
| **D3Z7F0** | L-lactate dehydrogenase (Fragment) OS=Mus musculus OX=10090 GN=Ldhb PE=1 SV=1 | 3 | 2 |
| **Q61937** | Nucleophosmin OS=Mus musculus OX=10090 GN=Npm1 PE=1 SV=1 | 2 | 2 |
| **Q9WVA4** | Transgelin-2 OS=Mus musculus OX=10090 GN=Tagln2 PE=1 SV=4 | 2 | 2 |
| **O35474** | EGF-like repeat and discoidin I-like domain-containing protein 3 OS=Mus musculus OX=10090 GN=Edil3 PE=1 SV=2 | 2 | 2 |

| | | | |
|---|---|---|---|
| **Q8VCM7** | Fibrinogen gamma chain OS=Mus musculus OX=10090 GN=Fgg PE=1 SV=1 | 2 | 2 |
| **F6UFG6** | Acidic leucine-rich nuclear phosphoprotein 32 family member A (Fragment) OS=Mus musculus OX=10090 GN=Anp32a PE=1 SV=1 | 2 | 2 |
| **O35326** | Serine/arginine-rich splicing factor 5 OS=Mus musculus OX=10090 GN=Srsf5 PE=1 SV=2 | 2 | 2 |
| **G3UZX4** | Casein kinase II subunit beta OS=Mus musculus OX=10090 GN=Csnk2b PE=1 SV=1 | 2 | 2 |
| **P68372** | Tubulin beta-4B chain OS=Mus musculus OX=10090 GN=Tubb4b PE=1 SV=1 | 14 | 2 |
| **P47753** | F-actin-capping protein subunit alpha-1 OS=Mus musculus OX=10090 GN=Capza1 PE=1 SV=4 | 3 | 2 |
| **Q9ESU7** | Amino acid transporter OS=Mus musculus OX=10090 GN=Slc1a5 PE=1 SV=1 | 2 | 2 |
| **O88569** | Heterogeneous nuclear ribonucleoproteins A2/B1 OS=Mus musculus OX=10090 GN=Hnrnpa2b1 PE=1 SV=2 | 2 | 2 |
| **Q07076** | Annexin A7 OS=Mus musculus OX=10090 GN=Anxa7 PE=1 SV=2 | 2 | 2 |
| **P16460** | Argininosuccinate synthase OS=Mus musculus OX=10090 GN=Ass1 PE=1 SV=1 | 2 | 2 |
| **P51150** | Ras-related protein Rab-7a OS=Mus musculus OX=10090 GN=Rab7a PE=1 SV=2 | 2 | 2 |
| **Q9CQV8** | 14-3-3 protein beta/alpha OS=Mus musculus OX=10090 GN=Ywhab PE=1 SV=3 | 4 | 2 |
| **Q62000** | Mimecan OS=Mus musculus OX=10090 GN=Ogn PE=1 SV=1 | 2 | 2 |
| **A0A1B0GQX9** | High mobility group protein B2 (Fragment) OS=Mus musculus OX=10090 GN=Hmgb2 PE=1 SV=1 | 3 | 2 |
| **P06800** | Receptor-type tyrosine-protein phosphatase C OS=Mus musculus OX=10090 GN=Ptprc PE=1 SV=4 | 2 | 2 |
| **P35278** | Ras-related protein Rab-5C OS=Mus musculus OX=10090 GN=Rab5c PE=1 SV=2 | 2 | 2 |
| **Q9WUM4** | Coronin-1C OS=Mus musculus OX=10090 GN=Coro1c PE=1 SV=2 | 2 | 2 |
| **P08113** | Endoplasmin OS=Mus musculus OX=10090 GN=Hsp90b1 PE=1 SV=2 | 2 | 2 |
| **Q3U367** | 4-trimethylaminobutyraldehyde dehydrogenase OS=Mus musculus OX=10090 GN=Aldh9a1 PE=1 SV=1 | 2 | 2 |
| **A0A087WPJ5** | Collagen alpha-1(III) chain (Fragment) OS=Mus musculus OX=10090 GN=Col3a1 PE=1 SV=1 | 2 | 2 |
| **B1B0C7** | Basement membrane-specific heparan sulfate proteoglycan core protein OS=Mus musculus OX=10090 GN=Hspg2 PE=1 SV=1 | 2 | 2 |
| **D3Z6I8** | Tropomyosin alpha-3 chain OS=Mus musculus OX=10090 GN=Tpm3 PE=1 SV=1 | 2 | 2 |
| **Q9CWH6** | Proteasome subunit alpha-type 7-like OS=Mus musculus OX=10090 GN=Psma8 PE=1 SV=1 | 2 | 2 |
| **P55264** | Adenosine kinase OS=Mus musculus OX=10090 GN=Adk PE=1 SV=2 | 2 | 2 |
| **P15535** | Beta-1,4-galactosyltransferase 1 OS=Mus musculus OX=10090 GN=B4galt1 PE=1 SV=1 | 2 | 2 |
| **P35979** | 60S ribosomal protein L12 OS=Mus musculus OX=10090 GN=Rpl12 PE=1 SV=2 | 2 | 2 |
| **Q8BSH9** | Nucleosome assembly protein 1-like 1 OS=Mus musculus OX=10090 GN=Nap1l1 PE=1 SV=1 | 2 | 2 |

| | | | |
|---|---|---|---|
| **Q9DCH4** | Eukaryotic translation initiation factor 3 subunit F OS=Mus musculus OX=10090 GN=Eif3f PE=1 SV=2 | 2 | 2 |
| **Q01149** | Collagen alpha-2(I) chain OS=Mus musculus OX=10090 GN=Col1a2 PE=1 SV=2 | 2 | 2 |
| **E9Q133** | T-complex protein 1 subunit gamma OS=Mus musculus OX=10090 GN=Cct3 PE=1 SV=1 | 2 | 2 |
| **Q9Z2X1** | Heterogeneous nuclear ribonucleoprotein F OS=Mus musculus OX=10090 GN=Hnrnpf PE=1 SV=3 | 2 | 2 |
| **A0A087WQ14** | Actin-related protein 3 (Fragment) OS=Mus musculus OX=10090 GN=Actr3 PE=1 SV=1 | 2 | 2 |
| **P63323** | 40S ribosomal protein S12 OS=Mus musculus OX=10090 GN=Rps12 PE=1 SV=2 | 2 | 2 |
| **Q9D8N0** | Elongation factor 1-gamma OS=Mus musculus OX=10090 GN=Eef1g PE=1 SV=3 | 2 | 2 |
| **P12382** | ATP-dependent 6-phosphofructokinase, liver type OS=Mus musculus OX=10090 GN=Pfkl PE=1 SV=4 | 2 | 2 |
| **P61161** | Actin-related protein 2 OS=Mus musculus OX=10090 GN=Actr2 PE=1 SV=1 | 2 | 2 |
| **Q8CI94** | Glycogen phosphorylase, brain form OS=Mus musculus OX=10090 GN=Pygb PE=1 SV=3 | 3 | 2 |
| **E0CXN0** | Hepatocyte growth factor-like protein OS=Mus musculus OX=10090 GN=Mst1 PE=1 SV=1 | 2 | 2 |
| **Q91VK2** | Eef1d protein OS=Mus musculus OX=10090 GN=Eef1d PE=1 SV=1 | 2 | 2 |
| **Q99LB4** | Capping protein (Actin filament), gelsolin-like OS=Mus musculus OX=10090 GN=Capg PE=1 SV=1 | 2 | 2 |
| **A2AP78** | High mobility group protein B3 (Fragment) OS=Mus musculus OX=10090 GN=Hmgb3 PE=1 SV=1 | 2 | 2 |
| **P62849** | 40S ribosomal protein S24 OS=Mus musculus OX=10090 GN=Rps24 PE=1 SV=1 | 2 | 2 |
| **P60867** | 40S ribosomal protein S20 OS=Mus musculus OX=10090 GN=Rps20 PE=1 SV=1 | 2 | 2 |
| **A2CEK3** | Phosphoglucomutase-1 OS=Mus musculus OX=10090 GN=Pgm1 PE=1 SV=1 | 2 | 2 |
| **Q9QUI0** | Transforming protein RhoA OS=Mus musculus OX=10090 GN=Rhoa PE=1 SV=1 | 2 | 2 |
| **Q9ET01** | Glycogen phosphorylase, liver form OS=Mus musculus OX=10090 GN=Pygl PE=1 SV=4 | 3 | 2 |
| **P08030** | Adenine phosphoribosyltransferase OS=Mus musculus OX=10090 GN=Aprt PE=1 SV=2 | 2 | 2 |
| **P62702** | 40S ribosomal protein S4, X isoform OS=Mus musculus OX=10090 GN=Rps4x PE=1 SV=2 | 2 | 2 |
| **Q922F4** | Tubulin beta-6 chain OS=Mus musculus OX=10090 GN=Tubb6 PE=1 SV=1 | 7 | 2 |
| **P60335** | Poly(rC)-binding protein 1 OS=Mus musculus OX=10090 GN=Pcbp1 PE=1 SV=1 | 2 | 2 |
| **D3Z1M1** | Guanine nucleotide-binding protein G(I)/G(S)/G(T) subunit beta-2 (Fragment) OS=Mus musculus OX=10090 GN=Gnb2 PE=1 SV=1 | 3 | 2 |
| **P63242** | Eukaryotic translation initiation factor 5A-1 OS=Mus musculus OX=10090 GN=Eif5a PE=1 SV=2 | 2 | 2 |
| **P12815** | Programmed cell death protein 6 OS=Mus musculus OX=10090 GN=Pdcd6 PE=1 SV=2 | 2 | 2 |
| **Q9CS84** | Neurexin-1 OS=Mus musculus OX=10090 GN=Nrxn1 PE=1 SV=3 | 2 | 2 |
| **P45376** | Aldo-keto reductase family 1 member B1 OS=Mus musculus OX=10090 GN=Akr1b1 PE=1 SV=3 | 2 | 2 |

| | | | |
|---|---|---|---|
| **Q8CF98** | Collectin-10 OS=Mus musculus OX=10090 GN=Colec10 PE=2 SV=1 | 2 | 2 |
| **P48758** | Carbonyl reductase [NADPH] 1 OS=Mus musculus OX=10090 GN=Cbr1 PE=1 SV=3 | 2 | 2 |
| **P62281** | 40S ribosomal protein S11 OS=Mus musculus OX=10090 GN=Rps11 PE=1 SV=3 | 2 | 2 |
| **Q80YX1** | Tenascin OS=Mus musculus OX=10090 GN=Tnc PE=1 SV=1 | 2 | 2 |
| **O09061** | Proteasome subunit beta type-1 OS=Mus musculus OX=10090 GN=Psmb1 PE=1 SV=1 | 2 | 2 |
| **H3BKU1** | Protein phosphatase 2 (Formerly 2A), regulatory subunit A (PR 65), beta isoform, isoform CRA_b OS=Mus musculus OX=10090 GN=Ppp2r1b PE=1 SV=1 | 2 | 2 |
| **Q5BKR2** | Sodium/hydrogen exchanger 9B2 OS=Mus musculus OX=10090 GN=Slc9b2 PE=1 SV=2 | 2 | 2 |
| **P10493** | Nidogen-1 OS=Mus musculus OX=10090 GN=Nid1 PE=1 SV=2 | 2 | 2 |
| **O08553** | Dihydropyrimidinase-related protein 2 OS=Mus musculus OX=10090 GN=Dpysl2 PE=1 SV=2 | 2 | 2 |
| **P08228** | Superoxide dismutase [Cu-Zn] OS=Mus musculus OX=10090 GN=Sod1 PE=1 SV=2 | 2 | 2 |
| **P09470** | Angiotensin-converting enzyme OS=Mus musculus OX=10090 GN=Ace PE=1 SV=3 | 2 | 2 |
| **A0A1B0GS70** | Proteasome endopeptidase complex OS=Mus musculus OX=10090 GN=Psma1 PE=1 SV=1 | 2 | 2 |
| **Q8JZQ9** | Eukaryotic translation initiation factor 3 subunit B OS=Mus musculus OX=10090 GN=Eif3b PE=1 SV=1 | 2 | 2 |
| **Q11136** | Xaa-Pro dipeptidase OS=Mus musculus OX=10090 GN=Pepd PE=1 SV=3 | 2 | 2 |
| **Q61171** | Peroxiredoxin-2 OS=Mus musculus OX=10090 GN=Prdx2 PE=1 SV=3 | 2 | 2 |
| **G3UZ34** | 116 kDa U5 small nuclear ribonucleoprotein component OS=Mus musculus OX=10090 GN=Eftud2 PE=1 SV=1 | 2 | 2 |
| **A0A087WNY6** | TAR DNA-binding protein 43 (Fragment) OS=Mus musculus OX=10090 GN=Tardbp PE=1 SV=1 | 2 | 2 |
| **E9Q1G8** | Septin-7 OS=Mus musculus OX=10090 GN=Sept7 PE=1 SV=2 | 2 | 2 |
| **P62315** | Small nuclear ribonucleoprotein Sm D1 OS=Mus musculus OX=10090 GN=Snrpd1 PE=1 SV=1 | 2 | 2 |
| **Q6P4T2** | U5 small nuclear ribonucleoprotein 200 kDa helicase OS=Mus musculus OX=10090 GN=Snrnp200 PE=1 SV=1 | 2 | 2 |
| **Q5SUF2** | Luc7-like protein 3 OS=Mus musculus OX=10090 GN=Luc7l3 PE=1 SV=1 | 2 | 2 |
| **P97326** | Cadherin-6 OS=Mus musculus OX=10090 GN=Cdh6 PE=1 SV=2 | 2 | 2 |
| **Q9JII6** | Aldo-keto reductase family 1 member A1 OS=Mus musculus OX=10090 GN=Akr1a1 PE=1 SV=3 | 2 | 2 |
| **P49312** | Heterogeneous nuclear ribonucleoprotein A1 OS=Mus musculus OX=10090 GN=Hnrnpa1 PE=1 SV=2 | 2 | 2 |
| **P42208** | Septin-2 OS=Mus musculus OX=10090 GN=Sept2 PE=1 SV=2 | 2 | 2 |
| **P01901** | H-2 class I histocompatibility antigen, K-B alpha chain OS=Mus musculus OX=10090 GN=H2-K1 PE=1 SV=1 | 2 | 2 |

| | | | |
|---|---|---|---|
| **P53994** | Ras-related protein Rab-2A OS=Mus musculus OX=10090 GN=Rab2a PE=1 SV=1 | 2 | 2 |
| **Q9DCD0** | 6-phosphogluconate dehydrogenase, decarboxylating OS=Mus musculus OX=10090 GN=Pgd PE=1 SV=3 | 2 | 2 |
| **Q9CZD3** | Glycine--tRNA ligase OS=Mus musculus OX=10090 GN=Gars PE=1 SV=1 | 2 | 2 |
| **P50396** | Rab GDP dissociation inhibitor alpha OS=Mus musculus OX=10090 GN=Gdi1 PE=1 SV=3 | 3 | 2 |
| **A0A0R4J0P5** | Proline-serine-threonine phosphatase-interacting protein 1 OS=Mus musculus OX=10090 GN=Pstpip1 PE=1 SV=1 | 2 | 2 |
| **P62082** | 40S ribosomal protein S7 OS=Mus musculus OX=10090 GN=Rps7 PE=2 SV=1 | 2 | 2 |
| **D3YWF6** | Ubiquitin thioesterase OTUB1 OS=Mus musculus OX=10090 GN=Otub1 PE=1 SV=1 | 2 | 2 |
| **A0A0N4SV00** | T-complex protein 1 subunit eta OS=Mus musculus OX=10090 GN=Cct7 PE=1 SV=1 | 2 | 2 |
| **O54890** | Integrin beta-3 OS=Mus musculus OX=10090 GN=Itgb3 PE=1 SV=2 | 2 | 2 |
| **Q8BFU2** | Histone H2A type 3 OS=Mus musculus OX=10090 GN=Hist3h2a PE=1 SV=3 | 4 | 1 |
| **P02088** | Hemoglobin subunit beta-1 OS=Mus musculus OX=10090 GN=Hbb-b1 PE=1 SV=2 | 2 | 1 |
| **A0A338P7H5** | Alpha-2-HS-glycoprotein OS=Mus musculus OX=10090 GN=Ahsg PE=1 SV=1 | 1 | 1 |
| **Q00623** | Apolipoprotein A-I OS=Mus musculus OX=10090 GN=Apoa1 PE=1 SV=2 | 1 | 1 |
| **F7CJN9** | Transferrin (Fragment) OS=Mus musculus OX=10090 GN=Trf PE=4 SV=1 | 1 | 1 |
| **P68433** | Histone H3.1 OS=Mus musculus OX=10090 GN=Hist1h3a PE=1 SV=2 | 3 | 1 |
| **Q9Z1R9** | MCG124046 OS=Mus musculus OX=10090 GN=Prss1 PE=1 SV=1 | 1 | 1 |
| **Q8BQM7** | Single-pass membrane and coiled-coil domain-containing protein 3 OS=Mus musculus OX=10090 GN=Smco3 PE=2 SV=1 | 1 | 1 |
| **P43274** | Histone H1.4 OS=Mus musculus OX=10090 GN=Hist1h1e PE=1 SV=2 | 3 | 1 |
| **P68134** | Actin, alpha skeletal muscle OS=Mus musculus OX=10090 GN=Acta1 PE=1 SV=1 | 8 | 1 |
| **V9GX81** | Maestro heat-like repeat family member 6 OS=Mus musculus OX=10090 GN=Mroh6 PE=1 SV=1 | 1 | 1 |
| **P84228** | Histone H3.2 OS=Mus musculus OX=10090 GN=Hist1h3b PE=1 SV=2 | 3 | 1 |
| **P15864** | Histone H1.2 OS=Mus musculus OX=10090 GN=Hist1h1c PE=1 SV=2 | 3 | 1 |
| **Q9CQM8** | 60S ribosomal protein L21 OS=Mus musculus OX=10090 GN=Rpl21 PE=1 SV=1 | 1 | 1 |
| **Q9DC71** | 28S ribosomal protein S15, mitochondrial OS=Mus musculus OX=10090 GN=Mrps15 PE=1 SV=2 | 1 | 1 |
| **D3YXF5** | Complement component 7 OS=Mus musculus OX=10090 GN=C7 PE=1 SV=2 | 1 | 1 |
| **A0A1L1ST95** | Down syndrome cell adhesion molecule-like protein 1 homolog OS=Mus musculus OX=10090 GN=Dscaml1 PE=4 SV=1 | 1 | 1 |

| | | | |
|---|---|---|---|
| **P84244** | Histone H3.3 OS=Mus musculus OX=10090 GN=H3f3a PE=1 SV=2 | 3 | 1 |
| **Q8K1I3** | Secreted phosphoprotein 24 OS=Mus musculus OX=10090 GN=Spp2 PE=1 SV=2 | 1 | 1 |
| **O89020** | Afamin OS=Mus musculus OX=10090 GN=Afm PE=1 SV=2 | 1 | 1 |
| **E9Q5L2** | Inter alpha-trypsin inhibitor, heavy chain 4 OS=Mus musculus OX=10090 GN=Itih4 PE=1 SV=1 | 1 | 1 |
| **Q923D2** | Flavin reductase (NADPH) OS=Mus musculus OX=10090 GN=Blvrb PE=1 SV=3 | 1 | 1 |
| **Q8VGW5** | Olfactory receptor OS=Mus musculus OX=10090 GN=Olfr691 PE=2 SV=1 | 1 | 1 |
| **O08677** | Kininogen-1 OS=Mus musculus OX=10090 GN=Kng1 PE=1 SV=1 | 1 | 1 |
| **Q9QZH3** | Peptidyl-prolyl cis-trans isomerase E OS=Mus musculus OX=10090 GN=Ppie PE=1 SV=2 | 1 | 1 |
| **Q8R1K1** | Ubiquitin-associated domain-containing protein 2 OS=Mus musculus OX=10090 GN=Ubac2 PE=1 SV=1 | 1 | 1 |
| **P13745** | Glutathione S-transferase A1 OS=Mus musculus OX=10090 GN=Gsta1 PE=1 SV=2 | 1 | 1 |
| **Q05144** | Ras-related C3 botulinum toxin substrate 2 OS=Mus musculus OX=10090 GN=Rac2 PE=1 SV=1 | 1 | 1 |
| **D3Z7C6** | Prostaglandin E synthase 3 OS=Mus musculus OX=10090 GN=Ptges3 PE=1 SV=1 | 1 | 1 |
| **S4R1W7** | Ras-related protein Rab-35 OS=Mus musculus OX=10090 GN=Rab35 PE=1 SV=1 | 1 | 1 |
| **Q9JHK5** | Pleckstrin OS=Mus musculus OX=10090 GN=Plek PE=1 SV=1 | 1 | 1 |
| **P02104** | Hemoglobin subunit epsilon-Y2 OS=Mus musculus OX=10090 GN=Hbb-y PE=1 SV=2 | 2 | 1 |
| **P49182** | Heparin cofactor 2 OS=Mus musculus OX=10090 GN=Serpind1 PE=1 SV=1 | 1 | 1 |
| **Q07968** | Coagulation factor XIII B chain OS=Mus musculus OX=10090 GN=F13b PE=1 SV=2 | 1 | 1 |
| **P24668** | Cation-dependent mannose-6-phosphate receptor OS=Mus musculus OX=10090 GN=M6pr PE=1 SV=1 | 1 | 1 |
| **Q08761** | Vitamin K-dependent protein S OS=Mus musculus OX=10090 GN=Pros1 PE=2 SV=1 | 1 | 1 |
| **Q62426** | Cystatin-B OS=Mus musculus OX=10090 GN=Cstb PE=1 SV=1 | 1 | 1 |
| **Q9D0J8** | Parathymosin OS=Mus musculus OX=10090 GN=Ptms PE=1 SV=3 | 1 | 1 |
| **P10639** | Thioredoxin OS=Mus musculus OX=10090 GN=Txn PE=1 SV=3 | 1 | 1 |
| **P47963** | 60S ribosomal protein L13 OS=Mus musculus OX=10090 GN=Rpl13 PE=1 SV=3 | 1 | 1 |
| **Q8CEB6** | Aldo-keto reductase family 1, member E1 OS=Mus musculus OX=10090 GN=Akr1e1 PE=1 SV=1 | 1 | 1 |
| **E9PV24** | Fibrinogen alpha chain OS=Mus musculus OX=10090 GN=Fga PE=1 SV=1 | 1 | 1 |
| **Q9QYX7** | Protein piccolo OS=Mus musculus OX=10090 GN=Pclo PE=1 SV=4 | 1 | 1 |
| **D3Z1S8** | 40S ribosomal protein S5 (Fragment) OS=Mus musculus OX=10090 GN=Rps5 PE=1 SV=1 | 1 | 1 |
| **A0A0G2JG40** | Guanine nucleotide-binding protein subunit alpha-12 (Fragment) OS=Mus musculus OX=10090 GN=Gna12 PE=1 SV=1 | 1 | 1 |

| | | | |
|---|---|---|---|
| **Q9R0N0** | Galactokinase OS=Mus musculus OX=10090 GN=Galk1 PE=1 SV=2 | 1 | 1 |
| **P28665** | Murinoglobulin-1 OS=Mus musculus OX=10090 GN=Mug1 PE=1 SV=3 | 1 | 1 |
| **Q64522** | Histone H2A type 2-B OS=Mus musculus OX=10090 GN=Hist2h2ab PE=1 SV=3 | 3 | 1 |
| **Q6URW6** | Myosin-14 OS=Mus musculus OX=10090 GN=Myh14 PE=1 SV=1 | 3 | 1 |
| **P49429** | 4-hydroxyphenylpyruvate dioxygenase OS=Mus musculus OX=10090 GN=Hpd PE=1 SV=3 | 1 | 1 |
| **P55284** | Cadherin-5 OS=Mus musculus OX=10090 GN=Cdh5 PE=1 SV=2 | 1 | 1 |
| **P30275** | Creatine kinase U-type, mitochondrial OS=Mus musculus OX=10090 GN=Ckmt1 PE=1 SV=1 | 1 | 1 |
| **A0A0R4J0S2** | Insulin-like growth factor-binding protein complex acid labile subunit OS=Mus musculus OX=10090 GN=Igfals PE=1 SV=1 | 1 | 1 |
| **P08207** | Protein S100-A10 OS=Mus musculus OX=10090 GN=S100a10 PE=1 SV=2 | 1 | 1 |
| **P20918** | Plasminogen OS=Mus musculus OX=10090 GN=Plg PE=1 SV=3 | 1 | 1 |
| **A2BH06** | 60S ribosomal protein L11 (Fragment) OS=Mus musculus OX=10090 GN=Rpl11 PE=1 SV=1 | 1 | 1 |
| **E9QNL5** | Sulfotransferase OS=Mus musculus OX=10090 GN=Sult1a1 PE=1 SV=2 | 1 | 1 |
| **Q5DU37** | Zinc finger FYVE domain-containing protein 26 OS=Mus musculus OX=10090 GN=Zfyve26 PE=1 SV=2 | 1 | 1 |
| **D3Z4N4** | Insulin-like growth factor II (Fragment) OS=Mus musculus OX=10090 GN=Igf2 PE=1 SV=2 | 1 | 1 |
| **G3UY13** | Interleukin-1 receptor accessory protein OS=Mus musculus OX=10090 GN=Il1rap PE=1 SV=1 | 1 | 1 |
| **D3YYK8** | Microtubule-associated protein RP/EB family member 2 (Fragment) OS=Mus musculus OX=10090 GN=Mapre2 PE=1 SV=1 | 1 | 1 |
| **Q9D1A2** | Cytosolic non-specific dipeptidase OS=Mus musculus OX=10090 GN=Cndp2 PE=1 SV=1 | 1 | 1 |
| **Q9QYB1** | Chloride intracellular channel protein 4 OS=Mus musculus OX=10090 GN=Clic4 PE=1 SV=3 | 1 | 1 |
| **Q6ZWV7** | 60S ribosomal protein L35 OS=Mus musculus OX=10090 GN=Rpl35 PE=1 SV=1 | 1 | 1 |
| **D3Z2C5** | Dimethylaniline monooxygenase [N-oxide-forming] 4 (Fragment) OS=Mus musculus OX=10090 GN=Fmo4 PE=3 SV=8 | 1 | 1 |
| **Q6ZWZ4** | 60S ribosomal protein L36 OS=Mus musculus OX=10090 GN=Rpl36 PE=1 SV=1 | 1 | 1 |
| **Q8BL97** | Serine/arginine-rich splicing factor 7 OS=Mus musculus OX=10090 GN=Srsf7 PE=1 SV=1 | 1 | 1 |
| **P35564** | Calnexin OS=Mus musculus OX=10090 GN=Canx PE=1 SV=1 | 1 | 1 |
| **Q91YI0** | Argininosuccinate lyase OS=Mus musculus OX=10090 GN=Asl PE=1 SV=1 | 1 | 1 |
| **P82198** | Transforming growth factor-beta-induced protein ig-h3 OS=Mus musculus OX=10090 GN=Tgfbi PE=1 SV=1 | 1 | 1 |
| **A2A4X6** | MCG21910 OS=Mus musculus OX=10090 GN=Gm12355 PE=4 SV=1 | 1 | 1 |

| | | | |
|---|---|---|---|
| **Q8BWP8** | Beta-1,4-glucuronyltransferase 1 OS=Mus musculus OX=10090 GN=B4gat1 PE=1 SV=1 | 1 | 1 |
| **A0A494BAP3** | Ferritin heavy chain (Fragment) OS=Mus musculus OX=10090 GN=Fth1 PE=4 SV=1 | 1 | 1 |
| **Q8C872** | Transferrin receptor protein 1 OS=Mus musculus OX=10090 GN=Tfrc PE=1 SV=1 | 1 | 1 |
| **O88200** | C-type lectin domain family 11 member A OS=Mus musculus OX=10090 GN=Clec11a PE=1 SV=1 | 1 | 1 |
| **Q9Z1T2** | Thrombospondin-4 OS=Mus musculus OX=10090 GN=Thbs4 PE=1 SV=1 | 3 | 1 |
| **Q9CVB6** | Actin-related protein 2/3 complex subunit 2 OS=Mus musculus OX=10090 GN=Arpc2 PE=1 SV=3 | 1 | 1 |
| **Q9CPW4** | Actin-related protein 2/3 complex subunit 5 OS=Mus musculus OX=10090 GN=Arpc5 PE=1 SV=3 | 1 | 1 |
| **P16294** | Coagulation factor IX OS=Mus musculus OX=10090 GN=F9 PE=2 SV=3 | 1 | 1 |
| **P21956** | Lactadherin OS=Mus musculus OX=10090 GN=Mfge8 PE=1 SV=3 | 1 | 1 |
| **Q9ES97** | Reticulon-3 OS=Mus musculus OX=10090 GN=Rtn3 PE=1 SV=2 | 1 | 1 |
| **P68254** | 14-3-3 protein theta OS=Mus musculus OX=10090 GN=Ywhaq PE=1 SV=1 | 3 | 1 |
| **P62715** | Serine/threonine-protein phosphatase 2A catalytic subunit beta isoform OS=Mus musculus OX=10090 GN=Ppp2cb PE=1 SV=1 | 1 | 1 |
| **F8WJ05** | Inter-alpha-trypsin inhibitor heavy chain H1 OS=Mus musculus OX=10090 GN=Itih1 PE=1 SV=1 | 1 | 1 |
| **P14152** | Malate dehydrogenase, cytoplasmic OS=Mus musculus OX=10090 GN=Mdh1 PE=1 SV=3 | 1 | 1 |
| **Q9EST5** | Acidic leucine-rich nuclear phosphoprotein 32 family member B OS=Mus musculus OX=10090 GN=Anp32b PE=1 SV=1 | 1 | 1 |
| **Q8BZF8** | Phosphoglucomutase-like protein 5 OS=Mus musculus OX=10090 GN=Pgm5 PE=1 SV=2 | 1 | 1 |
| **P16015** | Carbonic anhydrase 3 OS=Mus musculus OX=10090 GN=Ca3 PE=1 SV=3 | 1 | 1 |
| **P05132** | cAMP-dependent protein kinase catalytic subunit alpha OS=Mus musculus OX=10090 GN=Prkaca PE=1 SV=3 | 1 | 1 |
| **D6RHS6** | Phosphatidylethanolamine-binding protein 1 OS=Mus musculus OX=10090 GN=Pebp1 PE=1 SV=1 | 1 | 1 |
| **Q99KF1** | Transmembrane emp24 domain-containing protein 9 OS=Mus musculus OX=10090 GN=Tmed9 PE=1 SV=2 | 1 | 1 |
| **P62830** | 60S ribosomal protein L23 OS=Mus musculus OX=10090 GN=Rpl23 PE=1 SV=1 | 1 | 1 |
| **P57746** | V-type proton ATPase subunit D OS=Mus musculus OX=10090 GN=Atp6v1d PE=1 SV=1 | 1 | 1 |
| **P59999** | Actin-related protein 2/3 complex subunit 4 OS=Mus musculus OX=10090 GN=Arpc4 PE=1 SV=3 | 1 | 1 |
| **P51675** | C-C chemokine receptor type 1 OS=Mus musculus OX=10090 GN=Ccr1 PE=2 SV=2 | 1 | 1 |
| **Q9JKB3** | Y-box-binding protein 3 OS=Mus musculus OX=10090 GN=Ybx3 PE=1 SV=2 | 1 | 1 |
| **P61164** | Alpha-centractin OS=Mus musculus OX=10090 GN=Actr1a PE=1 SV=1 | 1 | 1 |

| | | | |
|---|---|---|---|
| **Q8BH61** | Coagulation factor XIII A chain OS=Mus musculus OX=10090 GN=F13a1 PE=1 SV=3 | 1 | 1 |
| **Q62093** | Serine/arginine-rich splicing factor 2 OS=Mus musculus OX=10090 GN=Srsf2 PE=1 SV=4 | 1 | 1 |
| **Q91X83** | S-adenosylmethionine synthase isoform type-1 OS=Mus musculus OX=10090 GN=Mat1a PE=1 SV=1 | 1 | 1 |
| **F8WJG3** | Transformer-2 protein homolog beta OS=Mus musculus OX=10090 GN=Tra2b PE=1 SV=1 | 1 | 1 |
| **P43275** | Histone H1.1 OS=Mus musculus OX=10090 GN=Hist1h1a PE=1 SV=2 | 2 | 1 |
| **Q9QZF2** | Glypican-1 OS=Mus musculus OX=10090 GN=Gpc1 PE=1 SV=1 | 1 | 1 |
| **Q91V41** | Ras-related protein Rab-14 OS=Mus musculus OX=10090 GN=Rab14 PE=1 SV=3 | 1 | 1 |
| **Q9QY76** | Vesicle-associated membrane protein-associated protein B OS=Mus musculus OX=10090 GN=Vapb PE=1 SV=3 | 1 | 1 |
| **F6Y6L6** | Predicted gene, 49369 (Fragment) OS=Mus musculus OX=10090 GN=Gm49369 PE=1 SV=1 | 1 | 1 |
| **Q8BH35** | Complement component C8 beta chain OS=Mus musculus OX=10090 GN=C8b PE=1 SV=1 | 1 | 1 |
| **O88844** | Isocitrate dehydrogenase [NADP] cytoplasmic OS=Mus musculus OX=10090 GN=Idh1 PE=1 SV=2 | 1 | 1 |
| **P61027** | Ras-related protein Rab-10 OS=Mus musculus OX=10090 GN=Rab10 PE=1 SV=1 | 1 | 1 |
| **A0A1L1SSH9** | SPARC OS=Mus musculus OX=10090 GN=Sparc PE=1 SV=1 | 1 | 1 |
| **P41105** | 60S ribosomal protein L28 OS=Mus musculus OX=10090 GN=Rpl28 PE=1 SV=2 | 1 | 1 |
| **Q3THE2** | Myosin regulatory light chain 12B OS=Mus musculus OX=10090 GN=Myl12b PE=1 SV=2 | 1 | 1 |
| **A2AD84** | Serine/threonine-protein kinase 26 OS=Mus musculus OX=10090 GN=Stk26 PE=1 SV=1 | 1 | 1 |
| **A0A1L1SUN1** | 60S ribosomal protein L29 (Fragment) OS=Mus musculus OX=10090 GN=Rpl29 PE=4 SV=1 | 1 | 1 |
| **O08710** | Thyroglobulin OS=Mus musculus OX=10090 GN=Tg PE=1 SV=3 | 1 | 1 |
| **Q9CR51** | V-type proton ATPase subunit G 1 OS=Mus musculus OX=10090 GN=Atp6v1g1 PE=1 SV=3 | 1 | 1 |
| **Q9Z0L8** | Gamma-glutamyl hydrolase OS=Mus musculus OX=10090 GN=Ggh PE=1 SV=2 | 1 | 1 |
| **P26040** | Ezrin OS=Mus musculus OX=10090 GN=Ezr PE=1 SV=3 | 2 | 1 |
| **P14602** | Heat shock protein beta-1 OS=Mus musculus OX=10090 GN=Hspb1 PE=1 SV=3 | 1 | 1 |
| **Q9QUM9** | Proteasome subunit alpha type-6 OS=Mus musculus OX=10090 GN=Psma6 PE=1 SV=1 | 1 | 1 |
| **Q3THW5** | Histone H2A.V OS=Mus musculus OX=10090 GN=H2afv PE=1 SV=3 | 3 | 1 |
| **P16627** | Heat shock 70 kDa protein 1-like OS=Mus musculus OX=10090 GN=Hspa1l PE=1 SV=4 | 5 | 1 |
| **D3YYV8** | 60S ribosomal protein L5 (Fragment) OS=Mus musculus OX=10090 GN=Rpl5 PE=1 SV=1 | 1 | 1 |
| **Q8R0Y6** | Cytosolic 10-formyltetrahydrofolate dehydrogenase OS=Mus musculus OX=10090 GN=Aldh1l1 PE=1 SV=1 | 1 | 1 |

| | | | |
|---|---|---|---|
| **E9PUB0** | Arf-GAP with Rho-GAP domain, ANK repeat and PH domain-containing protein 1 OS=Mus musculus OX=10090 GN=Arap1 PE=1 SV=1 | 1 | 1 |
| **P62320** | Small nuclear ribonucleoprotein Sm D3 OS=Mus musculus OX=10090 GN=Snrpd3 PE=1 SV=1 | 1 | 1 |
| **Q543K9** | Purine nucleoside phosphorylase OS=Mus musculus OX=10090 GN=Pnp PE=1 SV=1 | 1 | 1 |
| **A0A286YE28** | Transketolase (Fragment) OS=Mus musculus OX=10090 GN=Tkt PE=1 SV=1 | 1 | 1 |
| **P09055** | Integrin beta-1 OS=Mus musculus OX=10090 GN=Itgb1 PE=1 SV=1 | 1 | 1 |
| **Q61753** | D-3-phosphoglycerate dehydrogenase OS=Mus musculus OX=10090 GN=Phgdh PE=1 SV=3 | 1 | 1 |
| **Q9JK48** | Endophilin-B1 OS=Mus musculus OX=10090 GN=Sh3glb1 PE=1 SV=1 | 1 | 1 |
| **Q04519** | Sphingomyelin phosphodiesterase OS=Mus musculus OX=10090 GN=Smpd1 PE=1 SV=2 | 1 | 1 |
| **A0A0N4SW34** | V-type proton ATPase subunit E 1 (Fragment) OS=Mus musculus OX=10090 GN=Atp6v1e1 PE=1 SV=1 | 1 | 1 |
| **A2ALB3** | A disintegrin and metalloproteinase with thrombospondin motifs 13 OS=Mus musculus OX=10090 GN=Adamts13 PE=1 SV=1 | 1 | 1 |
| **Q8C845** | EF-hand domain-containing protein D2 OS=Mus musculus OX=10090 GN=Efhd2 PE=1 SV=1 | 1 | 1 |
| **Q80SY3** | V-type proton ATPase subunit d 2 OS=Mus musculus OX=10090 GN=Atp6v0d2 PE=2 SV=2 | 1 | 1 |
| **Q99PT1** | Rho GDP-dissociation inhibitor 1 OS=Mus musculus OX=10090 GN=Arhgdia PE=1 SV=3 | 1 | 1 |
| **P63087** | Serine/threonine-protein phosphatase PP1-gamma catalytic subunit OS=Mus musculus OX=10090 GN=Ppp1cc PE=1 SV=1 | 1 | 1 |
| **A0A1W2P7L5** | C-1-tetrahydrofolate synthase, cytoplasmic (Fragment) OS=Mus musculus OX=10090 GN=Mthfd1 PE=1 SV=1 | 1 | 1 |
| **Q9DBC7** | cAMP-dependent protein kinase type I-alpha regulatory subunit OS=Mus musculus OX=10090 GN=Prkar1a PE=1 SV=3 | 1 | 1 |
| **E9Q1V0** | Hsc70-interacting protein (Fragment) OS=Mus musculus OX=10090 GN=St13 PE=1 SV=1 | 1 | 1 |
| **O55234** | Proteasome subunit beta type-5 OS=Mus musculus OX=10090 GN=Psmb5 PE=1 SV=3 | 1 | 1 |
| **P46460** | Vesicle-fusing ATPase OS=Mus musculus OX=10090 GN=Nsf PE=1 SV=2 | 1 | 1 |
| **D3YUT3** | 40S ribosomal protein S19 (Fragment) OS=Mus musculus OX=10090 GN=Rps19 PE=1 SV=2 | 1 | 1 |
| **P20060** | Beta-hexosaminidase subunit beta OS=Mus musculus OX=10090 GN=Hexb PE=1 SV=2 | 1 | 1 |
| **Q62095** | ATP-dependent RNA helicase DDX3Y OS=Mus musculus OX=10090 GN=Ddx3y PE=1 SV=2 | 1 | 1 |
| **D3Z712** | 40S ribosomal protein S15a (Fragment) OS=Mus musculus OX=10090 GN=Rps15a PE=1 SV=2 | 1 | 1 |
| **Q9D0R2** | Threonine--tRNA ligase, cytoplasmic OS=Mus musculus OX=10090 GN=Tars PE=1 SV=2 | 1 | 1 |
| **P47754** | F-actin-capping protein subunit alpha-2 OS=Mus musculus OX=10090 GN=Capza2 PE=1 SV=3 | 2 | 1 |

| | | | |
|---|---|---|---|
| **P00920** | Carbonic anhydrase 2 OS=Mus musculus OX=10090 GN=Ca2 PE=1 SV=4 | 1 | 1 |
| **Q00915** | Retinol-binding protein 1 OS=Mus musculus OX=10090 GN=Rbp1 PE=1 SV=2 | 1 | 1 |
| **O55134** | Protocadherin-12 OS=Mus musculus OX=10090 GN=Pcdh12 PE=1 SV=2 | 1 | 1 |
| **P49722** | Proteasome subunit alpha type-2 OS=Mus musculus OX=10090 GN=Psma2 PE=1 SV=3 | 1 | 1 |
| **Q99J77** | Sialic acid synthase OS=Mus musculus OX=10090 GN=Nans PE=1 SV=1 | 1 | 1 |
| **A0A0N4SUQ1** | Plasminogen activator inhibitor 1 RNA-binding protein OS=Mus musculus OX=10090 GN=Serbp1 PE=1 SV=1 | 1 | 1 |
| **Q8R4V5** | Oncoprotein-induced transcript 3 protein OS=Mus musculus OX=10090 GN=Oit3 PE=2 SV=2 | 1 | 1 |
| **A0A0U1RNT6** | S-adenosylmethionine synthase OS=Mus musculus OX=10090 GN=Mat2a PE=1 SV=1 | 1 | 1 |
| **A0A2R8W6U6** | Poly(rC)-binding protein 2 (Fragment) OS=Mus musculus OX=10090 GN=Pcbp2 PE=1 SV=1 | 1 | 1 |
| **P10852** | 4F2 cell-surface antigen heavy chain OS=Mus musculus OX=10090 GN=Slc3a2 PE=1 SV=1 | 1 | 1 |
| **Q921R2** | 40S ribosomal protein S13 OS=Mus musculus OX=10090 GN=Rps13 PE=1 SV=1 | 1 | 1 |
| **E9PY39** | Predicted gene 20431 OS=Mus musculus OX=10090 GN=Gm20431 PE=4 SV=1 | 1 | 1 |
| **P60229** | Eukaryotic translation initiation factor 3 subunit E OS=Mus musculus OX=10090 GN=Eif3e PE=1 SV=1 | 1 | 1 |
| **P21995** | Embigin OS=Mus musculus OX=10090 GN=Emb PE=1 SV=2 | 1 | 1 |
| **P08905** | Lysozyme C-2 OS=Mus musculus OX=10090 GN=Lyz2 PE=1 SV=2 | 1 | 1 |
| **Q9CQI6** | Coactosin-like protein OS=Mus musculus OX=10090 GN=Cotl1 PE=1 SV=3 | 1 | 1 |
| **P68040** | Receptor of activated protein C kinase 1 OS=Mus musculus OX=10090 GN=Rack1 PE=1 SV=3 | 1 | 1 |
| **Q05722** | Collagen alpha-1(IX) chain OS=Mus musculus OX=10090 GN=Col9a1 PE=2 SV=2 | 1 | 1 |
| **B1AY86** | Plexin domain-containing protein 2 OS=Mus musculus OX=10090 GN=Plxdc2 PE=1 SV=1 | 1 | 1 |
| **A0A2R8VHW2** | Ribosomal L1 domain-containing protein 1 (Fragment) OS=Mus musculus OX=10090 GN=Rsl1d1 PE=1 SV=1 | 1 | 1 |
| **P59470** | DNA-directed RNA polymerase III subunit RPC2 OS=Mus musculus OX=10090 GN=Polr3b PE=1 SV=2 | 1 | 1 |
| **P70460** | Vasodilator-stimulated phosphoprotein OS=Mus musculus OX=10090 GN=Vasp PE=1 SV=4 | 1 | 1 |
| **Q91WP0** | Mannan-binding lectin serine protease 2 OS=Mus musculus OX=10090 GN=Masp2 PE=1 SV=1 | 1 | 1 |
| **P21981** | Protein-glutamine gamma-glutamyltransferase 2 OS=Mus musculus OX=10090 GN=Tgm2 PE=1 SV=4 | 1 | 1 |
| **O09131** | Glutathione S-transferase omega-1 OS=Mus musculus OX=10090 GN=Gsto1 PE=1 SV=2 | 1 | 1 |
| **P97464** | Exostosin-1 OS=Mus musculus OX=10090 GN=Ext1 PE=1 SV=1 | 1 | 1 |
| **E9PUD2** | Dynamin-1-like protein OS=Mus musculus OX=10090 GN=Dnm1l PE=1 SV=1 | 1 | 1 |
| **A0A0R4J119** | Cytoplasmic FMR1-interacting protein OS=Mus musculus OX=10090 GN=Cyfip1 PE=1 SV=1 | 1 | 1 |
| **B9EJ86** | Oxysterol-binding protein-related protein 8 OS=Mus musculus OX=10090 GN=Osbpl8 PE=1 SV=1 | 1 | 1 |

| | | | |
|---|---|---|---|
| **P97798** | Neogenin OS=Mus musculus OX=10090 GN=Neo1 PE=1 SV=1 | 1 | 1 |
| **H3BJV3** | Serpin A11 (Fragment) OS=Mus musculus OX=10090 GN=Serpina11 PE=1 SV=1 | 1 | 1 |
| **A0A0G2JF67** | Phosphodiesterase OS=Mus musculus OX=10090 GN=Pde5a PE=1 SV=1 | 1 | 1 |
| **Q8R3X6** | Glypican-6 OS=Mus musculus OX=10090 GN=Gpc6 PE=1 SV=1 | 1 | 1 |
| **P26369** | Splicing factor U2AF 65 kDa subunit OS=Mus musculus OX=10090 GN=U2af2 PE=1 SV=3 | 1 | 1 |
| **Q91Y97** | Fructose-bisphosphate aldolase B OS=Mus musculus OX=10090 GN=Aldob PE=1 SV=3 | 2 | 1 |
| **Q62165** | Dystroglycan OS=Mus musculus OX=10090 GN=Dag1 PE=1 SV=4 | 1 | 1 |
| **Q8BK67** | Protein RCC2 OS=Mus musculus OX=10090 GN=Rcc2 PE=1 SV=1 | 1 | 1 |
| **P46467** | Vacuolar protein sorting-associated protein 4B OS=Mus musculus OX=10090 GN=Vps4b PE=1 SV=2 | 1 | 1 |
| **Q8BPU7** | Engulfment and cell motility protein 1 OS=Mus musculus OX=10090 GN=Elmo1 PE=1 SV=2 | 1 | 1 |
| **P07742** | Ribonucleoside-diphosphate reductase large subunit OS=Mus musculus OX=10090 GN=Rrm1 PE=1 SV=2 | 1 | 1 |
| **Q9D0I9** | Arginine--tRNA ligase, cytoplasmic OS=Mus musculus OX=10090 GN=Rars PE=1 SV=2 | 1 | 1 |
| **O35295** | Transcriptional activator protein Pur-beta OS=Mus musculus OX=10090 GN=Purb PE=1 SV=3 | 1 | 1 |
| **Q8CJ40** | Rootletin OS=Mus musculus OX=10090 GN=Crocc PE=1 SV=2 | 1 | 1 |
| **A2A841** | Protein 4.1 OS=Mus musculus OX=10090 GN=Epb41 PE=1 SV=1 | 1 | 1 |
| **P05063** | Fructose-bisphosphate aldolase C OS=Mus musculus OX=10090 GN=Aldoc PE=1 SV=4 | 2 | 1 |
| **P14824** | Annexin A6 OS=Mus musculus OX=10090 GN=Anxa6 PE=1 SV=3 | 1 | 1 |
| **Q3TCR7** | Dynamin-2 OS=Mus musculus OX=10090 GN=Dnm2 PE=1 SV=1 | 1 | 1 |
| **E0CYW7** | Hepatoma-derived growth factor OS=Mus musculus OX=10090 GN=Hdgf PE=1 SV=1 | 1 | 1 |
| **D3Z4B0** | Serine/arginine-rich-splicing factor 11 (Fragment) OS=Mus musculus OX=10090 GN=Srsf11 PE=1 SV=1 | 1 | 1 |
| **A0A1C7CYV0** | Plastin-3 (Fragment) OS=Mus musculus OX=10090 GN=Pls3 PE=1 SV=1 | 4 | 1 |
| **P63005** | Platelet-activating factor acetylhydrolase IB subunit alpha OS=Mus musculus OX=10090 GN=Pafah1b1 PE=1 SV=2 | 1 | 1 |
| **Q8QZR4** | Out at first protein homolog OS=Mus musculus OX=10090 GN=Oaf PE=2 SV=1 | 1 | 1 |
| **Q62465** | Synaptic vesicle membrane protein VAT-1 homolog OS=Mus musculus OX=10090 GN=Vat1 PE=1 SV=3 | 1 | 1 |
| **Q9QXY6** | EH domain-containing protein 3 OS=Mus musculus OX=10090 GN=Ehd3 PE=1 SV=2 | 2 | 1 |
| **P16045** | Galectin-1 OS=Mus musculus OX=10090 GN=Lgals1 PE=1 SV=3 | 1 | 1 |
| **O55222** | Integrin-linked protein kinase OS=Mus musculus OX=10090 GN=Ilk PE=1 SV=2 | 1 | 1 |
| **Q5SYD0** | Unconventional myosin-Id OS=Mus musculus OX=10090 GN=Myo1d PE=1 SV=1 | 1 | 1 |

| | | | |
|---|---|---|---|
| **A0A0N4SUN5** | Nuclear receptor-interacting protein 2 (Fragment) OS=Mus musculus OX=10090 GN=Nrip2 PE=4 SV=1 | 1 | 1 |
| **O35658** | Complement component 1 Q subcomponent-binding protein, mitochondrial OS=Mus musculus OX=10090 GN=C1qbp PE=1 SV=1 | 1 | 1 |
| **Q9WTI7** | Unconventional myosin-Ic OS=Mus musculus OX=10090 GN=Myo1c PE=1 SV=2 | 1 | 1 |
| **Q6A0A9** | Constitutive coactivator of PPAR-gamma-like protein 1 OS=Mus musculus OX=10090 GN=FAM120A PE=1 SV=2 | 1 | 1 |
| **P63028** | Translationally-controlled tumor protein OS=Mus musculus OX=10090 GN=Tpt1 PE=1 SV=1 | 1 | 1 |
| **O54833** | Casein kinase II subunit alpha' OS=Mus musculus OX=10090 GN=Csnk2a2 PE=1 SV=1 | 1 | 1 |
| **G3UZ26** | Serine hydroxymethyltransferase (Fragment) OS=Mus musculus OX=10090 GN=Shmt1 PE=1 SV=1 | 1 | 1 |
| **Q8BVQ9** | 26S proteasome regulatory subunit 7 OS=Mus musculus OX=10090 GN=Psmc2 PE=1 SV=1 | 1 | 1 |
| **Q9JJI8** | 60S ribosomal protein L38 OS=Mus musculus OX=10090 GN=Rpl38 PE=1 SV=3 | 1 | 1 |
| **A7E1W8** | SIGLEC-I OS=Mus musculus OX=10090 GN=Siglec15 PE=2 SV=1 | 1 | 1 |
| **Q9EP69** | Phosphatidylinositide phosphatase SAC1 OS=Mus musculus OX=10090 GN=Sacm1l PE=1 SV=1 | 1 | 1 |
| **Q9DBG6** | Dolichyl-diphosphooligosaccharide--protein glycosyltransferase subunit 2 OS=Mus musculus OX=10090 GN=Rpn2 PE=1 SV=1 | 1 | 1 |
| **Q9Z1T1** | AP-3 complex subunit beta-1 OS=Mus musculus OX=10090 GN=Ap3b1 PE=1 SV=2 | 1 | 1 |
| **P62317** | Small nuclear ribonucleoprotein Sm D2 OS=Mus musculus OX=10090 GN=Snrpd2 PE=1 SV=1 | 1 | 1 |
| **P35822** | Receptor-type tyrosine-protein phosphatase kappa OS=Mus musculus OX=10090 GN=Ptprk PE=1 SV=1 | 1 | 1 |
| **Q9CR16** | Peptidyl-prolyl cis-trans isomerase D OS=Mus musculus OX=10090 GN=Ppid PE=1 SV=3 | 1 | 1 |
| **Q00PI9** | Heterogeneous nuclear ribonucleoprotein U-like protein 2 OS=Mus musculus OX=10090 GN=Hnrnpul2 PE=1 SV=2 | 1 | 1 |
| **Q6P9Q4** | FH1/FH2 domain-containing protein 1 OS=Mus musculus OX=10090 GN=Fhod1 PE=1 SV=3 | 1 | 1 |
| **Q9Z0K8** | Pantetheinase OS=Mus musculus OX=10090 GN=Vnn1 PE=1 SV=3 | 1 | 1 |
| **Q9D0M5** | Dynein light chain 2, cytoplasmic OS=Mus musculus OX=10090 GN=Dynll2 PE=1 SV=1 | 1 | 1 |
| **P28301** | Protein-lysine 6-oxidase OS=Mus musculus OX=10090 GN=Lox PE=1 SV=1 | 1 | 1 |
| **Q9D1H9** | Microfibril-associated glycoprotein 4 OS=Mus musculus OX=10090 GN=Mfap4 PE=1 SV=1 | 1 | 1 |
| **Q6Q477** | Plasma membrane calcium-transporting ATPase 4 OS=Mus musculus OX=10090 GN=Atp2b4 PE=1 SV=1 | 1 | 1 |
| **Q9Z1N5** | Spliceosome RNA helicase Ddx39b OS=Mus musculus OX=10090 GN=Ddx39b PE=1 SV=1 | 1 | 1 |
| **P62334** | 26S proteasome regulatory subunit 10B OS=Mus musculus OX=10090 GN=Psmc6 PE=1 SV=1 | 1 | 1 |
| **Q9R1P1** | Proteasome subunit beta type-3 OS=Mus musculus OX=10090 GN=Psmb3 PE=1 SV=1 | 1 | 1 |

| | | | |
|---|---|---|---|
| **P46735** | Unconventional myosin-Ib OS=Mus musculus OX=10090 GN=Myo1b PE=1 SV=3 | 1 | 1 |
| **Q9QZE5** | Coatomer subunit gamma-1 OS=Mus musculus OX=10090 GN=Copg1 PE=1 SV=1 | 1 | 1 |
| **Q9R1P0** | Proteasome subunit alpha type-4 OS=Mus musculus OX=10090 GN=Psma4 PE=1 SV=1 | 1 | 1 |
| **Q08943** | FACT complex subunit SSRP1 OS=Mus musculus OX=10090 GN=Ssrp1 PE=1 SV=2 | 1 | 1 |
| **E9PYK3** | Protein mono-ADP-ribosyltransferase PARP4 OS=Mus musculus OX=10090 GN=Parp4 PE=1 SV=1 | 1 | 1 |
| **P52194** | Calmegin OS=Mus musculus OX=10090 GN=Clgn PE=1 SV=2 | 1 | 1 |
| **P46638** | Ras-related protein Rab-11B OS=Mus musculus OX=10090 GN=Rab11b PE=1 SV=3 | 1 | 1 |
| **A0A0G2JE29** | Procollagen C-endopeptidase enhancer 1 (Fragment) OS=Mus musculus OX=10090 GN=Pcolce PE=1 SV=1 | 1 | 1 |
| **A0A1B0GSU0** | Aldehyde dehydrogenase family 16 member A1 OS=Mus musculus OX=10090 GN=Aldh16a1 PE=1 SV=1 | 1 | 1 |
| **Q80VQ0** | Aldehyde dehydrogenase family 3 member B1 OS=Mus musculus OX=10090 GN=Aldh3b1 PE=1 SV=1 | 1 | 1 |
| **P70168** | Importin subunit beta-1 OS=Mus musculus OX=10090 GN=Kpnb1 PE=1 SV=2 | 1 | 1 |
| **A0A338P6M5** | 26S proteasome non-ATPase regulatory subunit 2 (Fragment) OS=Mus musculus OX=10090 GN=Psmd2 PE=1 SV=1 | 1 | 1 |
| **O89023** | Tripeptidyl-peptidase 1 OS=Mus musculus OX=10090 GN=Tpp1 PE=1 SV=2 | 1 | 1 |
| **Q91YQ5** | Dolichyl-diphosphooligosaccharide--protein glycosyltransferase subunit 1 OS=Mus musculus OX=10090 GN=Rpn1 PE=1 SV=1 | 1 | 1 |
| **Q9ES52** | Phosphatidylinositol 3,4,5-trisphosphate 5-phosphatase 1 OS=Mus musculus OX=10090 GN=Inpp5d PE=1 SV=2 | 1 | 1 |
| **O55029** | Coatomer subunit beta' OS=Mus musculus OX=10090 GN=Copb2 PE=1 SV=2 | 1 | 1 |
| **Q8VHY0** | Chondroitin sulfate proteoglycan 4 OS=Mus musculus OX=10090 GN=Cspg4 PE=1 SV=3 | 1 | 1 |
| **B1AWD6** | Golgi-associated plant pathogenesis-related protein 1 OS=Mus musculus OX=10090 GN=Glipr2 PE=1 SV=1 | 1 | 1 |
| **A0A494BA97** | Osteoclast-stimulating factor 1 OS=Mus musculus OX=10090 GN=Ostf1 PE=4 SV=1 | 1 | 1 |
| **P14069** | Protein S100-A6 OS=Mus musculus OX=10090 GN=S100a6 PE=1 SV=3 | 1 | 1 |
| **E9PZG9** | Small nuclear ribonucleoprotein E (Fragment) OS=Mus musculus OX=10090 GN=Snrpe PE=1 SV=1 | 1 | 1 |
| **Q00724** | Retinol-binding protein 4 OS=Mus musculus OX=10090 GN=Rbp4 PE=1 SV=2 | 1 | 1 |
| **Q5SQX6** | Cytoplasmic FMR1-interacting protein 2 OS=Mus musculus OX=10090 GN=Cyfip2 PE=1 SV=2 | 1 | 1 |
| **A2AC16** | Dicarbonyl L-xylulose reductase, isoform CRA_a OS=Mus musculus OX=10090 GN=Dcxr PE=1 SV=1 | 1 | 1 |
| **Q8K183** | Pyridoxal kinase OS=Mus musculus OX=10090 GN=Pdxk PE=1 SV=1 | 1 | 1 |
| **Q9D6F9** | Tubulin beta-4A chain OS=Mus musculus OX=10090 GN=Tubb4a PE=1 SV=3 | 12 | 1 |

| | | | |
|---|---|---|---|
| **D6RGM3** | Echinoderm microtubule-associated protein-like 2 OS=Mus musculus OX=10090 GN=Eml2 PE=1 SV=1 | 1 | 1 |
| **Q60864** | Stress-induced-phosphoprotein 1 OS=Mus musculus OX=10090 GN=Stip1 PE=1 SV=1 | 1 | 1 |
| **Q60790** | Ras GTPase-activating protein 3 OS=Mus musculus OX=10090 GN=Rasa3 PE=1 SV=2 | 1 | 1 |
| **P29351** | Tyrosine-protein phosphatase non-receptor type 6 OS=Mus musculus OX=10090 GN=Ptpn6 PE=1 SV=2 | 1 | 1 |
| **P33434** | 72 kDa type IV collagenase OS=Mus musculus OX=10090 GN=Mmp2 PE=1 SV=1 | 1 | 1 |
| **O70503** | Very-long-chain 3-oxoacyl-CoA reductase OS=Mus musculus OX=10090 GN=Hsd17b12 PE=1 SV=1 | 1 | 1 |
| **P62192** | 26S proteasome regulatory subunit 4 OS=Mus musculus OX=10090 GN=Psmc1 PE=1 SV=1 | 1 | 1 |
| **Q8R1F1** | Niban-like protein 1 OS=Mus musculus OX=10090 GN=Fam129b PE=1 SV=2 | 1 | 1 |
| **Q8K1K2** | 26S proteasome regulatory subunit 8 OS=Mus musculus OX=10090 GN=Psmc5 PE=1 SV=1 | 1 | 1 |
| **P97315** | Cysteine and glycine-rich protein 1 OS=Mus musculus OX=10090 GN=Csrp1 PE=1 SV=3 | 1 | 1 |
| **Q8BH43** | Wiskott-Aldrich syndrome protein family member 2 OS=Mus musculus OX=10090 GN=Wasf2 PE=1 SV=1 | 1 | 1 |
| **Q8JZK9** | Hydroxymethylglutaryl-CoA synthase, cytoplasmic OS=Mus musculus OX=10090 GN=Hmgcs1 PE=1 SV=1 | 1 | 1 |
| **P97807** | Fumarate hydratase, mitochondrial OS=Mus musculus OX=10090 GN=Fh PE=1 SV=3 | 1 | 1 |
| **P28481** | Collagen alpha-1(II) chain OS=Mus musculus OX=10090 GN=Col2a1 PE=1 SV=2 | 1 | 1 |
| **P11152** | Lipoprotein lipase OS=Mus musculus OX=10090 GN=Lpl PE=1 SV=3 | 1 | 1 |
| **Q9JJU8** | SH3 domain-binding glutamic acid-rich-like protein OS=Mus musculus OX=10090 GN=Sh3bgrl PE=1 SV=1 | 1 | 1 |
| **Q9R182** | Angiopoietin-related protein 3 OS=Mus musculus OX=10090 GN=Angptl3 PE=1 SV=1 | 1 | 1 |
| **A2ARJ3** | Transmembrane protein 236 OS=Mus musculus OX=10090 GN=Tmem236 PE=3 SV=1 | 1 | 1 |
| **Q9JMA7** | Cytochrome P450 3A41 OS=Mus musculus OX=10090 GN=Cyp3a41a PE=1 SV=2 | 1 | 1 |
| **A0A0G2JDW6** | Aminoacyl tRNA synthase complex-interacting multifunctional protein 1 (Fragment) OS=Mus musculus OX=10090 GN=Aimp1 PE=1 SV=4 | 1 | 1 |
| **A0A1Y7VNP4** | Eukaryotic translation initiation factor 5 (Fragment) OS=Mus musculus OX=10090 GN=Eif5 PE=1 SV=1 | 1 | 1 |
| **F6TIL5** | von Willebrand factor A domain-containing protein 5A (Fragment) OS=Mus musculus OX=10090 GN=Vwa5a PE=1 SV=1 | 1 | 1 |
| **P62911** | 60S ribosomal protein L32 OS=Mus musculus OX=10090 GN=Rpl32 PE=1 SV=2 | 1 | 1 |
| **Q8C129** | Leucyl-cystinyl aminopeptidase OS=Mus musculus OX=10090 GN=Lnpep PE=1 SV=1 | 1 | 1 |
| **Q922Q8** | Leucine-rich repeat-containing protein 59 OS=Mus musculus OX=10090 GN=Lrrc59 PE=1 SV=1 | 1 | 1 |
| **Q80TE4** | Signal-induced proliferation-associated 1-like protein 2 OS=Mus musculus OX=10090 GN=Sipa1l2 PE=1 SV=3 | 1 | 1 |

| | | | |
|---|---|---|---|
| **P27046** | Alpha-mannosidase 2 OS=Mus musculus OX=10090 GN=Man2a1 PE=1 SV=2 | 1 | 1 |
| **P70697** | Uroporphyrinogen decarboxylase OS=Mus musculus OX=10090 GN=Urod PE=1 SV=2 | 1 | 1 |
| **Q61316** | Heat shock 70 kDa protein 4 OS=Mus musculus OX=10090 GN=Hspa4 PE=1 SV=1 | 1 | 1 |
| **G5E8F1** | Integrin alpha-M OS=Mus musculus OX=10090 GN=Itgam PE=1 SV=2 | 1 | 1 |
| **Q64523** | Histone H2A type 2-C OS=Mus musculus OX=10090 GN=Hist2h2ac PE=1 SV=3 | 4 | 0 |
